# Pervasive species-specific repulsion among adult tropical trees

**DOI:** 10.1101/2022.02.12.480223

**Authors:** Michael Kalyuzhny, Jeffrey K. Lake, S. Joseph Wright, Annette M. Ostling

## Abstract

For species to coexist, performance must decline as the density of conspecific individuals increases. Although evidence for such Conspecific Negative Density Dependence (CNDD) exists in forests, the spatial repulsion it should produce has not been demonstrated in adults. Here we show that in comparison to a null model of stochastic birth, death and limited dispersal, the adults of dozens of tropical forest tree species show almost ubiquitous and strong spatial repulsion, some to surprising distances of ~100 meters. We use simulations to show such strong repulsion can only occur if CNDD considerably exceeds Heterospecific Negative Density Dependence, an even stronger condition required for coexistence, and that large-scale repulsion can indeed result from small-scale CNDD. These results highlight the power of limited dispersal spatial null models.

**One-Sentence Summary:** Spatial distributions of tropical trees reflect a strong negative effect of conspecifics, exceeding that of heterospecifics

## Main Text

How can hundreds of species coexist on small spatial scales? Some tropical forests contain over 250 species of trees in a single hectare, or over 1,100 species of trees and shrub in 25 hectares (*1*), begging for mechanistic explanation. According to coexistence theory, a necessary condition for species coexistence is that the performance of species decreases as they become more common, preventing any one species from taking over, a phenomenon known as Conspecific Negative Density Dependence (CNDD) (*2, 3*). A stronger necessary condition is that CNDD is stronger than Heterospecific Negative Density Dependence (HNDD) (*2, 3*), implying that species mostly ‘suffer’ from their own kind due to niche differences, that stabilize population size. Coexistence is assured if, in addition to the aforementioned condition, species are sufficiently similar in their competitive ability (*3, 4*).

Considerable evidence has been presented for CNDD acting on early life stages in tropical and temperate forests, and variation in CNDD has been hypothesized to underlie the latitudinal gradient from high tree species diversity at the equator to lower diversity at higher latitudes (*3, 5–7*). For example, the abundance of seedlings, seeds and adults have been shown by numerous works to reduce the number and survival of seedlings and saplings (*8–11*) and several works have also presented evidence that CNDD is stronger than HNDD (*8, 11*). Nevertheless, the validity of many of these findings has been questioned on methodological grounds (*2, 10*). Crucially, current evidence suggests that CNDD has very limited effect on adults (*12, 13*). Moreover, adult tropical trees can live for hundreds of years (*14*) and integrate density dependent effects throughout their life. CNDD at one life stage can be offset by positive effects at another (*2, 15*), or have a weak effect on lifetime performance, begging the question – what is the overall integrated effect of conspecifics on populations? Importantly, this integrated effect is what matters for coexistence (2).

A potential approach to evaluating the integrated effect of CNDD across life stages is through the study of the spatial distribution of adult trees. Because CNDD has been shown to operate at short scales around adults (*3, 8, 9*), it is theorized to generate spatial ‘repulsion’ between adults, potentially leading to spatial overdispersion (*16, 17*). Alas, classical studies of the spatial distributions of adult trees found them to be almost ubiquitously *aggregated* in tropical forests (*18–20*). Hence, individuals are typically closer to each other than expected, especially at small scales where CNDD is expected to be the strongest (*18–20*). Furthermore, many spatial patterns in tropical forests resemble the expectations of neutral models (*21–23*), or reflect habitat heterogeneity (*24–26*), with little signs of CNDD. These findings suggest that sensitivity to CNDD at early life stages is severely diluted, overwhelmed or obscured before reaching adulthood by processes that create aggregation such as habitat specificity, facilitation and dispersal limitation (*2, 3*). If CNDD at early life stages does not propagate to affect the adults (as evidence for this is lacking (*27*)), how can it maintain species diversity?

Classical studies on spatial patterns of adult trees compared their distributions to the expectation of the Complete Spatial Randomness (CSR) null model (*16, 18, 19*). This model distributes individuals with equal probability across the (often arbitrary) survey area, inevitably assigning some to sites that they would not be able to reach due to dispersal limitation (Fig. 1a,b). We believe that a more ecologically sound null model is Dispersal Limitation (DL, spatially explicit neutral theory) (*21, 29*), which assumes that abundance and distribution are determined by stochastic births and deaths and random, yet limited, dispersal. Using DL as a null model should incorporate the aggregation created by dispersal limitation into the null expectations (Fig 1c, Fig. S1-2), potentially revealing signals of biotic interactions and habitat specificity.

**Fig. 1.**
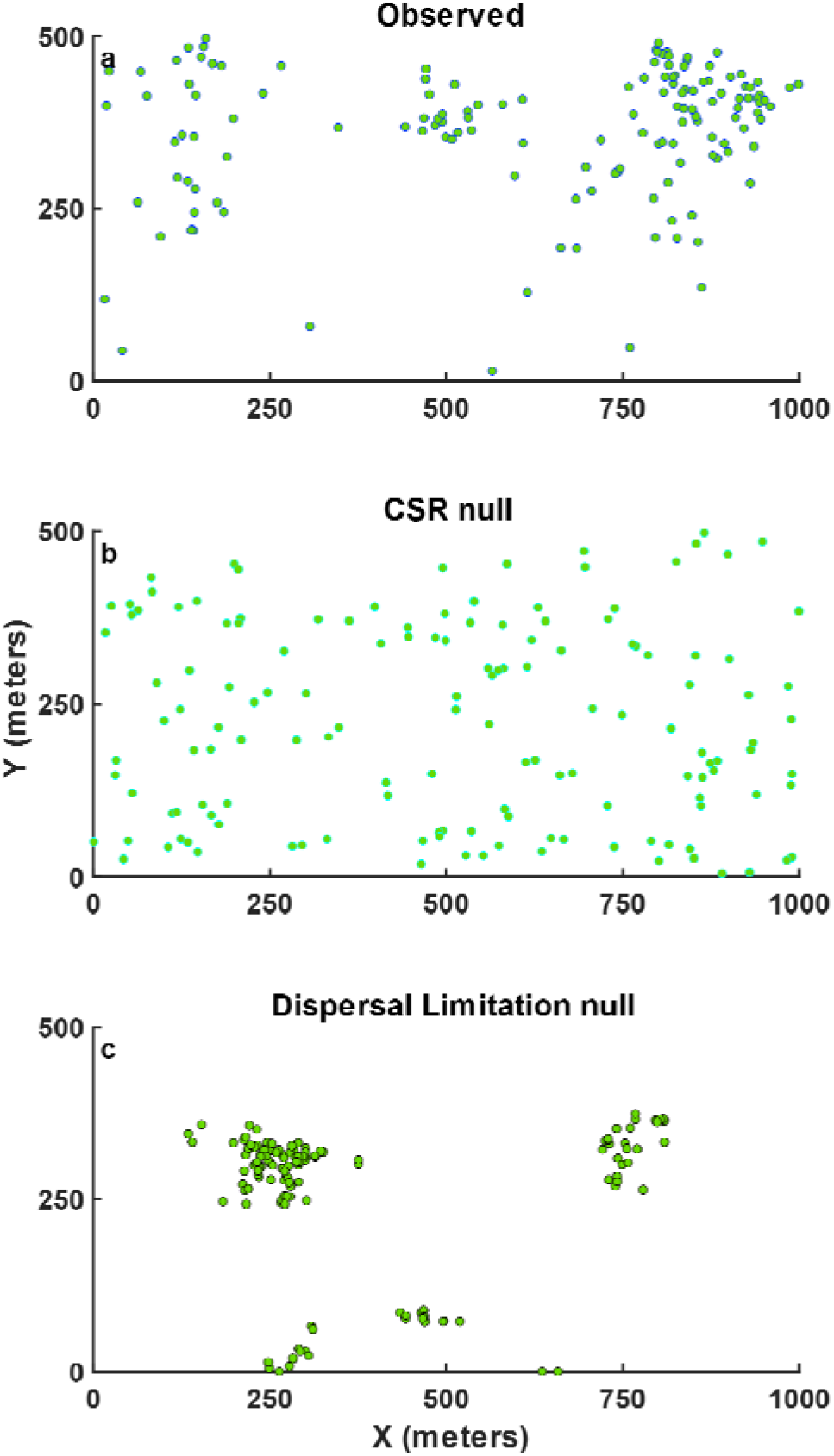
Spatial point patterns of adult *Beilschmiedia pendula* trees; The (**A**) observed pattern appears aggregated compared to (**B**) realizations of the Complete Spatial Randomness (CSR) null model, but overdispersed compared to (**C**) the standard variant of the Dispersal Limitation (DL) null model. Both nulls use the same abundance as the observed. The mean dispersal distance of *B. pendula* is estimated at 10.7 meters (*28*). Fig. S1 presents realizations of the other variants of DL, while Fig. S2 zooms in on the clump of trees at the top-left of (**C**).

Here we compare the spatial patterns of tropical tree species in a Panamanian forest to their expectation under DL, using previously estimated species-specific dispersal distance distributions (*28, 30*). Our goal is to examine the prevalence, magnitude and scale of overdispersion and aggregation with respect to DL. Since spatial repulsion is the signature of CNDD (*16, 31*), which is caused by local interactions and has been shown to operate on small spatial scales (~20 m., *3, 8, 9*), we hypothesize that some species, for which CNDD propagates to the adult stage, would show overdispersion at small scales. We then use a spatially explicit simulation model to examine whether the spatial patterns we find provide insight into the relative strength of CNDD versus HNDD. Finally, we examine the implications of our findings for tropical tree diversity.

We perform the analysis with 41 species that have reliable estimates of dispersal distance distributions, obtained using known locations of seed-bearing trees and seed arrivals (*28*) in the Barro Colorado Island 50 ha forest plot (BCI) (*32*). These estimates have been corroborated using genotyped seeds and adults (*33*), highlighting their reliability. DL simulations, parameterized with the dispersal distance distributions, are run for every species separately, and the observed spatial distribution of adults in 2015 is compared to the expectation under DL. For that comparison we develop analogs of the two most used point-pattern statistics: the Clark-Evans nearest neighbor statistic and the relative neighborhood density (a.k.a. pair correlation function) (*16, 18, 19*) statistic. We compare the observed mean (or median, see methods) distance of trees to their nearest conspecific neighbor, *NND_obs_*, to its value under the null (*NND_null_*) using the Excess Distance (*ED*) statistic:

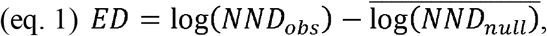

where the averaging (denoted by an overbar in eqs. 1 and 2) is over samples of the DL model. We also compute the Excess neighborhood Abundance (*EA*(*r*)), namely the observed number of conspecific neighbors of an average tree at distances *r* to *r* + Δ*r*, *N_obs_*(*r*), relative to its value under the null *N_null_*(*r*):

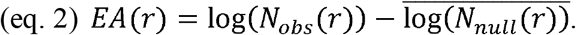

To examine the robustness of our results (‘Standard’), we repeat this analysis for the standard DL null hypothesis and five additional variants of the DL null model. The five variants 1) use earlier estimates of dispersal, for a broader set of 81 species (‘H2001’ (*30*)); 2) add 10% of global dispersal, representing widespread long-distance dispersal (‘LDD’); these variants consider different types of dispersal distance distributions and increase the dispersion under the null due to longer dispersal distances, making the analysis more conservative; 3) incorporate a lag of one generation from dispersal to recruitment, likewise increasing dispersion under the null since parent and offspring often do not co-occur; 4) start with the distribution of conspecific trees in 1985 and examine only the recruits to the adult class appearing by 2015 (‘Recruits’); 5) Fix the observed locations of all adult trees (‘Fixed’), while allowing their species identity to vary dynamically, controlling the spatial distribution of trees of all species. Finally, we also show that clustered dispersal that may result from animals depositing clusters of seeds (*34*) would typically produce aggregation compared to our null models with independent seeds (see Supplementary Text).

Contrary to previous findings based on CSR (*18, 19*), BCI tree species are almost ubiquitously and strongly overdispersed on short scales (~20 meters) when compared to all six variants of the Dispersal Limitation null model (Table 1). On average, trees have 50% - 80% fewer conspecific neighbors than expected within 20 meters, and their nearest conspecific neighbor is 1.8 - 2.5 times more distant than expected (these numbers are the values of the mean statistics in Table 1, exponentiated). Even the most common shrubs, treelets and trees (e.g. *Trichilia tuberculata*, Fig. 2), are overdispersed, as are relatively rare species (Data S1) and these patterns are qualitatively similar under the different DL null model variants (Table 1). Fig. 2 shows typical, monotonically increasing *EA*(*r*)*s* for four species, along with their distribution maps in comparison with the expectations of the DL null, while Fig. S3-4 show *EA*(*r*) for all non-rare species we analyzed. Not surprisingly, some light-demanding and habitat specialist species deviate from this typical pattern, exhibiting increased aggregation reflective of the limited spatial extent of the conditions they require (Fig. S5).

**Table 1.**
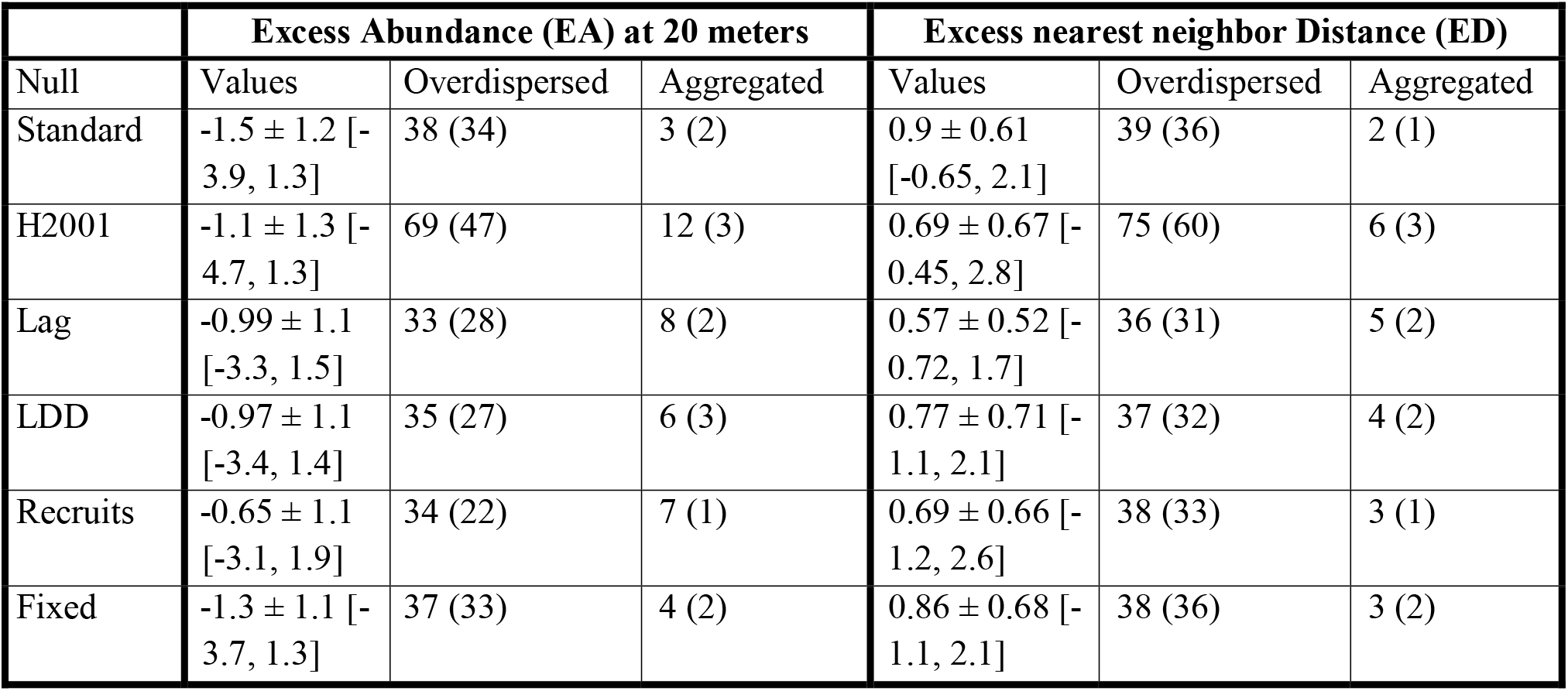
**Distributions and significance of the spatial statistics of Barro Colorado tree species in** 2015 using the six variants of the DL null model as references. We compare the abundance of conspecific neighbors within 20 meters (left, eq. 2) and the nearest neighbor distance (right, eq. 1) to their expectations. For abundances (left), negative values represent repulsion and vice versa. For distances (right), positive values represent repulsion. Exponentiating the statistics would give the factor by which the observed density or distance exceeds expectations. The mean ± SD and the range (in square brackets) of the statistics over all species are presented, as well as the number of overdispersed and aggregated species, along with the number of statistically significant (with *α* = 0.05) species in parentheses. For the LDD and Recruits nulls, median nearest neighbor distance is used, while the mean is calculated in the other cases.

|  | Excess Abundance (EA) at 20 meters |  |  | Excess nearest neighbor Distance (ED) |  |  |
| --- | --- | --- | --- | --- | --- | --- |
| Null | Values | Overdispersed | Aggregated | Values | Overdispersed | Aggregated |
| Standard | $-1.5 \pm 1.2$ [-3.9, 1.3] | 38 (34) | 3 (2) | $0.9 \pm 0.61$ [-0.65, 2.1] | 39 (36) | 2 (1) |
| H2001 | $-1.1 \pm 1.3$ [-4.7, 1.3] | 69 (47) | 12 (3) | $0.69 \pm 0.67$ [-0.45, 2.8] | 75 (60) | 6 (3) |
| Lag | $-0.99 \pm 1.1$ [-3.3, 1.5] | 33 (28) | 8 (2) | $0.57 \pm 0.52$ [-0.72, 1.7] | 36 (31) | 5 (2) |
| LDD | $-0.97 \pm 1.1$ [-3.4, 1.4] | 35 (27) | 6 (3) | $0.77 \pm 0.71$ [-1.1, 2.1] | 37 (32) | 4 (2) |
| Recruits | $-0.65 \pm 1.1$ [-3.1, 1.9] | 34 (22) | 7 (1) | $0.69 \pm 0.66$ [-1.2, 2.6] | 38 (33) | 3 (1) |
| Fixed | $-1.3 \pm 1.1$ [-3.7, 1.3] | 37 (33) | 4 (2) | $0.86 \pm 0.68$ [-1.1, 2.1] | 38 (36) | 3 (2) |

**Fig. 2.**
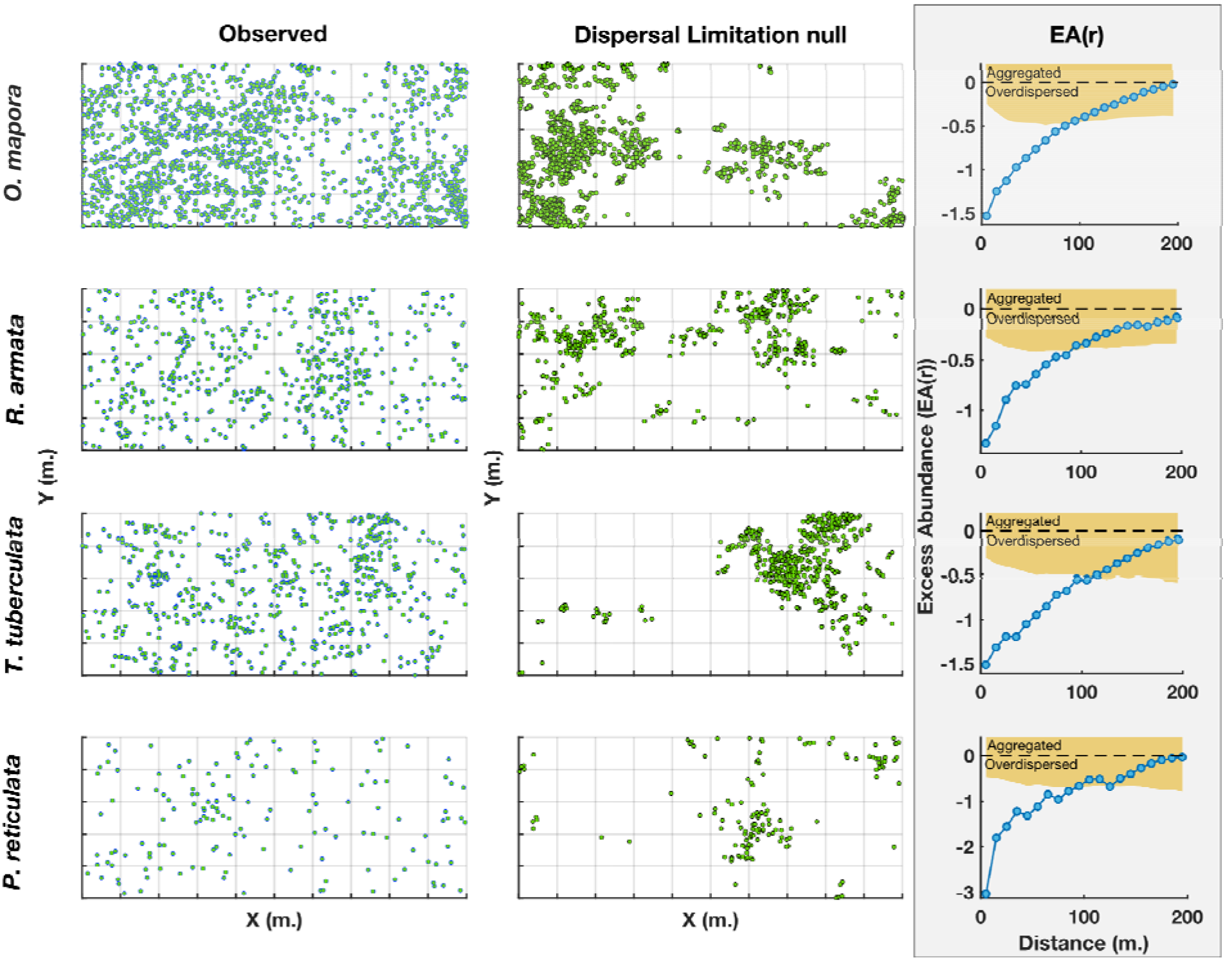
**Spatial distributions of four typical species and their Excess neighbor abundance at distance *r***, *EA*(*r*) (eq. 2). Each row presents a different species, while the columns present for each species the observed distribution (left) versus a single realization of the Standard DL null (center). *EA*(*r*) is presented in the rightmost, shaded column, with the orange region indicating the 95% simulation envelope of the DL null. *EA*(*r*) of 0 indicates agreement with DL while values < 0 indicate overdispersion. All distribution maps are 1000 by 500 meters. Compared to the DL null, trees have considerably fewer neighbors than expected at distances up to 100 m.

What underlies this widespread, strong and robust (under different assumptions of the null models) overdispersion? Theoretical studies show that while aggregation can be caused by dispersal limitation (*17, 35*), habitat specificity (*31*) and facilitation (*36*), overdispersion is the signature of negative density dependence (*16, 17*). Hence, we infer that the overdispersion we detect reflects the spatial signature of density dependence acting on juveniles, for which much evidence on BCI exists (*8, 10, 11, 37*), propagating to affect the adults.

Importantly, we show here using a simulation model that the observed overdispersion is only consistent with a scenario when CNDD >> HNDD. Figures 3 and S6 show the degree of overdispersion generated in spatially explicit stochastic simulations of community dynamics where CNDD and HNDD both reduce the survival of dispersing juveniles and are set to be strong out to a distance of about 7m. (Fig. S10). First, when the magnitude of CNDD = HNDD (along the diagonal in Fig. 3), overdispersion in the simulations quickly asymptotes with the magnitude of density dependence, at values indicating much lower overdispersion than observed on BCI: while *EA*(20) is ~ −1.5 in BCI (Table 1), the simulated *EA*(20) asymptotes at much lower values (~ −0.18 or ~−0.55, Fig. 3, S7-8). Intuitively, when HNDD = CNDD, the competitive effect of neighbors is equal regardless of their species, and so density dependence cannot generate a strong repulsion from conspecifics. Second, only the scenario CNDD >> HNDD (upper left corners in Fig. 3) is compatible with the magnitude of overdispersion observed on BCI. Finally, our results using the ‘Fixed’ null variant (Table 1), in which the locations of all adults are preserved, are similar to the analysis using the standard null variant. The ‘Fixed’ variant preserves the spacing between all trees of all species, resulting from competition for light, soil resources and other non-species-specific factors, hence incorporating HNDD and some CNDD into the null expectation. The strong overdispersion compared with this variant further confirms the weakness of HNDD compared to CNDD.

**Fig. 3.**
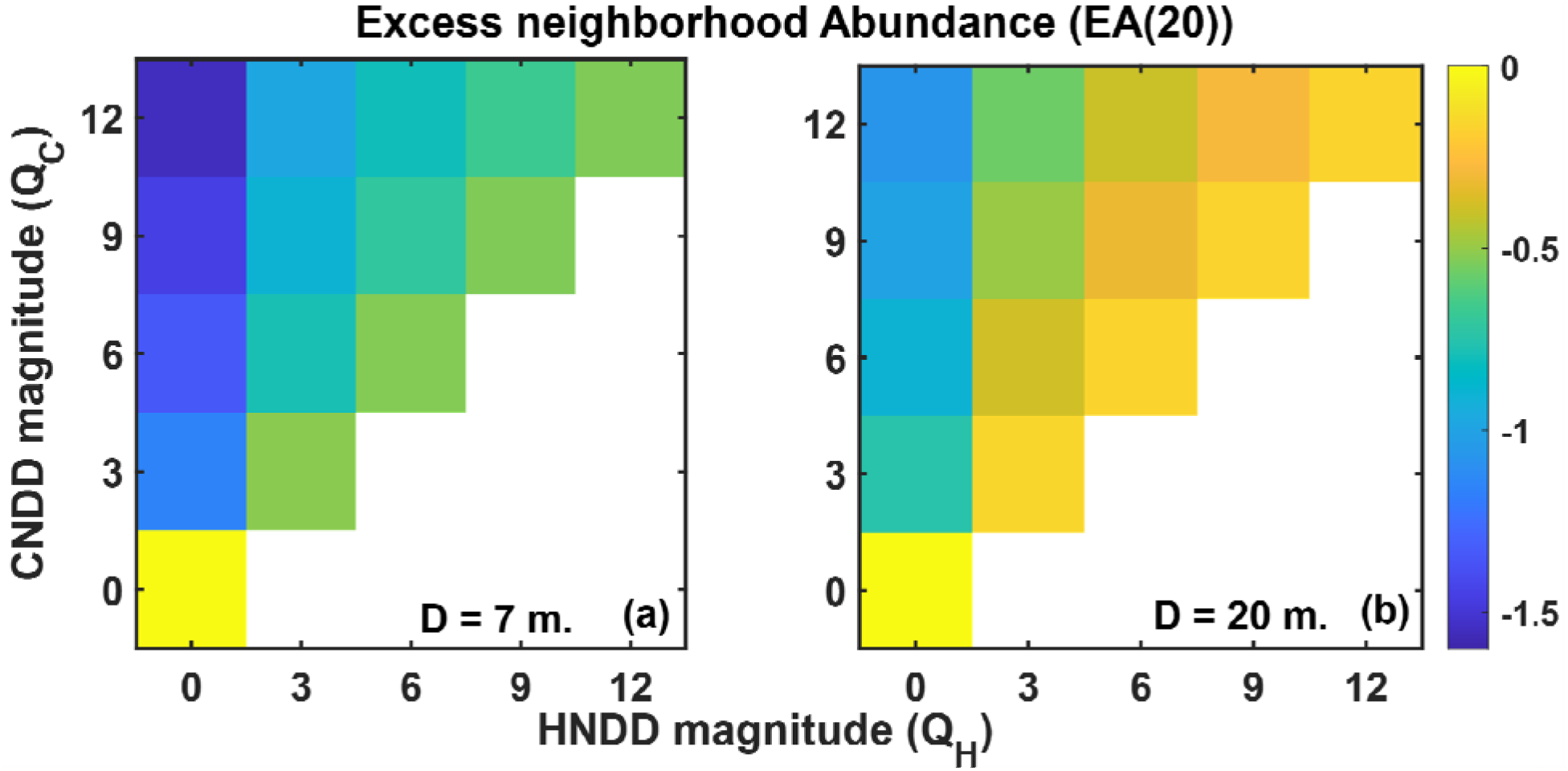
Excess neighborhood Abundance at 20 meters, *EA*(**20**) (eq. 2) in simulations. In each simulation, the magnitude of CNDD (*Q_C_*, Y axis) and HNDD (*Q_H_*, X axis) were controlled, and the color of each square represents the mean *EA*(20) of simulated populations. In the simulations, seeds are dispersed an average distance of *D* = 7 meters (**a**) or *D* = 20 meters (**b**) and the survival of each is determined by both the density of conspecific adults and the density of heterospecific adults in a way that gives large weight to adults within ~ 7 meters of where the seed lands. For perspective, a seed landing right near a single conspecific would have a recruitment probability of 1/(1+*QC*), and more nearby conspecifics and heterospecifics would reduce this probability further. See Methods and eq. S3 for more details. Results are not shown for cases where HNDD > CNDD (below the diagonal) since priority effects lead to one species taking over these communities. When CNDD = HNDD (on the diagonal) overdispersion is weak, and the strongest overdispersion is observed when CNDD ≫ HNDD.

While we expected overdispersion to occur primarily at the short distances at which CNDD operates (*8, 11*), we find several species are significantly and substantially overdispersed at distances up to ~100 meters, with *all* species having on average ~ 45% fewer neighbors 75 – 125 meters away from a typical tree (under the Standard null and all variants except the ‘LDD’ or ‘Recruits’ null, Figure 2, Table S1). Our simulations show that such long-distance overdispersion can result from short-distance CNDD (acting to seven meters) (Fig. S9). In hindsight, this result appeals to intuition. While each tree “repels” only its neighbors, they in turn repel their neighbors and so forth, creating large-scale patterns from short-distance interactions.

Previous analyses aimed at detecting CNDD from spatial patterns have reached contradictory conclusions. Many did not detect evidence of CNDD (*21–26*). Several revealed spatial repulsion of juveniles from conspecific adults (e.g., *7, 9*). Others fit complex process-based models to all individuals (including subadults) finding evidence for CNDD (*35, 38*). However, none of these studies provide evidence that the effects of CNDD propagate to affect the adult stage. Focusing on larger trees and community-level spatial pattern, May et al. (*21*) found that *all* trees ≥ 10 CM DBH (*not only conspecifics*) are overdispersed compared to CSR at distances < 10 m, indicating general density dependence among all trees of all species. Several studies found that conspecific aggregation decreases with tree size (*18, 20, 39*), but those that considered more than a handful of species found significant reductions in only 15% - 22% of species studied (*18, 20*). In contrast, we found that almost all species are overdispersed at the adult stage. This difference might occur as the effects of CNDD accumulate through ontogeny with the strongest effects at early life stages (*12*) before the sapling stage considered by references 18, 20. This highlights the utility of comparison to the DL null model, which uses seed rain as the reference. Finally, no previous study of spatial pattern evaluated the relative strength of CNDD and HNDD, while we show that species-specific spatial patterns of adults are only consistent with a scenario when CNDD >> HNDD across life stages.

These findings have important implications for the maintenance of species diversity. First, our finding that CNDD must be substantially greater than HNDD for many species implies large niche differences among species, which may stabilize diversity because species that increase in abundance suffer reduced fitness. It has recently been debated whether differences in species-level fitness can overwhelm the stabilizing effects of CNDD (*40, 41*), so the potential of CNDD to maintain species diversity needs further investigation. However, our work provides a key missing ingredient in this line of research on diversity maintenance in forests — namely evidence that the overall forces of density dependence *across life stages* set up substantial stabilizing niche differences, a key requirement for stable coexistence. Moreover, given that the forest is overall saturated with individuals (*29*), the finding that individuals of most species have fewer *conspecific* neighbors up to long distances implies an increase in *heterospecific* neighbors and an increase in compositional turnover at these distances. This is consistent with the reported increased spatial turnover (beta diversity) at scales of 20-200 meters in BCI compared with a neutral model (*22*). Beta diversity has been extensively studied and is believed to be driven mostly by spatial heterogeneity, dispersal limitation and interspecific competition (*22, 42, 43*). Our results suggest that the role of CNDD in shaping turnover should also be carefully considered.

More broadly, patterns of species distributions at local scales have been investigated and compared to CSR for over one hundred years (*44*), often with the aim of detecting the signals of habitat specificity and biotic interactions (*18, 19, 26, 45*). However, theoretical models predict that dispersal limitation alone could override and obscure the signal of biotic interactions and dominate spatial patterns, creating ubiquitous aggregated distributions (*17*). Indeed, spatial patterns have been found to be ubiquitously aggregated with respect to the CSR null (*18, 19, 45*). In contrast, a richness of spatial patterns is revealed with respect to the DL null: most species are overdispersed, owing to CNDD, while some habitat and gap specialists are more aggregated (Fig. S5). This demonstrates the power of the DL null to filter out much of the effect of dispersal limitation and to reveal the overall effect of biotic interactions and habitat specificity on the spatial distributions of adult individuals. Our work thus promotes a complementary definition of the classical terms “aggregation” and “overdispersion”, rooted in ecological processes and theory. We believe that a more widespread use of this approach will have much capacity to shed light on the processes shaping the assembly of ecological communities.

## Acknowledgments

We thank R. Condit and R. Kadmon for comments. BCI forest dynamics research project was made possible by National Science Foundation grants to Stephen P. Hubbell: DEB-0640386, DEB-0425651, DEB-0346488, DEB-0129874, DEB-00753102, DEB-9909347, DEB-9615226, DEB-9615226, DEB-9405933, DEB-9221033, DEB-9100058, DEB-8906869, DEB-8605042, DEB-8206992, DEB-7922197, support from the Forest Global Earth Observatory, the Smithsonian Tropical Research Institute, the John D. and Catherine T. MacArthur Foundation, the Mellon Foundation, the Small World Institute Fund, and numerous private individuals, and through the hard work of over 100 people from 10 countries over the past three decades. The plot project is part the Forest Global Earth Observatory (ForestGEO), a global network of large-scale demographic tree plots.

## Funding

Michigan Life Sciences Fellowship (MK)

Zuckerman STEM Leadership Program (MK)

Sabbatical support from the University of Michigan and Adrian College (JKL)

Mcubed, The University of Michigan (MK, AMO)

Associate Professor Support Fund (MK, AMO)

## Author contributions

Conceptualization: MK

Planning: MK, JKL, AMO

Data generation: SJW

Data analysis: MK

Writing – original draft: MK

Interpreting results: MK, JKL, SJW, AMO

Writing – review & editing: MK, JKL, SJW, AMO

## Data and materials availability

All data used in this analysis is publicly available and referenced to. All the code used to generate the results is included in the supplementary.

## Supplementary Materials

Materials and Methods

Supplementary Text

Figs. S1 to S11

Tables S1

References (*46–65*)

Data S1

