## Supplementary material for "Pervasive species-specific repulsion among adult tropical trees": Methods

This PDF file includes:

Materials and Methods

Supplementary Text

Figs. S1 to S11

Table S1

Other Supplementary Materials for this manuscript include the following:

Data S1 – Species level data

Materials and Methods

All code used in this analysis is provided in the code appendix.

Dispersal Estimates

A dispersal kernel describes the probability that a seed would travel and land in a specific unit area at a distance *r* from a tree, while a dispersal distance distribution describes the probability the seed would land *somewhere* at a distance *r* (*30, 46*). Generally, the dispersal distance distribution is obtained from the estimated dispersal kernel through radial integration, but for our simulations we use the distance distributions with parameters that were estimated using the kernels.

We use two sets of dispersal estimates that have been obtained previously for tropical trees in Barro Colorado Island (*32, 47, 48*) using inverse-modeling with seed trap and adult census data. The main source we used (‘H-2008’) estimated the single parameter of a 2DT dispersal kernel with 3 degrees of freedom (*46, 49*), which was chosen as the best overall, for 41 species (*28*). This work also incorporated year to year variation in fecundity and dispersal from outside the census plot, finding a mean dispersal distance of 27 meters across species. Importantly, a comparison of the methodology used to obtain the ‘H-2008’ kernels with estimates based on genotyped seeds and adults revealed remarkable similarity (*33*), boosting our confidence in these estimates. The second set, published earlier in a dissertation (‘H-2001’)(*30*), is comprised of 81 species (including 40 species from the first set), used a Weibull dispersal kernel and has a mean dispersal distance of 40 meters. Since this is an earlier estimate that had not gone through peer review, we consider results obtained with this set as a “robustness test” of the main results with a broader set of species, larger mean dispersal distance and an alternative functional form of the kernel.

Because of the evidence that some species may have surprisingly high levels of long-distance dispersal (*50*), we also considered a mixed kernel, with 90% of the seeds dispersed according to the H-2008 kernel (*28*), and 10% distributed completely at random.

These two alternative kernels have larger mean dispersal distance, which leads to larger distancing and lower densities of individuals under the null. Compared to these nulls, observed distances may be less deviant, making the analyses using them more conservative for detecting repulsion.

A few comments about the estimates of dispersal are in order. First, the primary estimates of dispersal distance (‘H-2008’) include some uncertainty, yet the credibility intervals for dispersal distance are not very large. Moreover, despite the validation of the H-2008 estimates with an alternative method, there may still be uncertainty about the proper model for the kernel. We address this source of uncertainty by considering kernels with alternative forms and with larger mean dispersal distances, for which the analyses are more conservative.

Furthermore, our null models assume that seeds are dispersed independently of trees other than their parent and of each other. In reality, both assumptions could be inaccurate. First, seed dispersal could be density dependent, with more seeds dispersed to locations where adult density is high. This could be caused by animal foragers congregating in such areas (*34*), thereby increasing the chances seeds would be dispersed there. Moreover, such foragers could potentially become satiated, leaving many of the seeds without effective dispersal vectors (*34*). The second assumption could also be violated, since seeds are often dispersed in clumps, mostly by animal vectors (*28, 30, 51*). Regarding density dependent seed dispersal – this mechanism would increase the local densities of seeds around clumps of adults, making our null model conservative in terms of detecting adult repulsion. Similarly, clumped dispersal creates clumps of multiple seeds, and the recruitment of multiple seeds from the same clump increases the aggregation of adults. The effect of clumped dispersal is analyzed in Supplementary Text 1. Hence, in most scenarios, the independence assumption we use is conservative for detecting repulsion.

Finally, the subset of species that have dispersal data is not a random subset. These species exclude shrubs, for which not all reproductive adults are recorded, and have seeds large enough to remain in the mesh nets of the seed traps. This subset includes multiple rare species, with 13 out of the 41 species having abundance < 1 adult per hectare. The rare species turn out not less overdispersed (data S1).

The Dispersal Limitation null model

The observed spatial distributions of species are compared to the spatial point patterns generated by the Dispersal Limitation (DL) null model, a spatially-explicit, continuous time and space version of neutral theory (*21, 29*). Hence, the null model incorporates dispersal limitation and drift and eliminates all forms of niche processes. We use several versions of this null model, representing alternative assumptions. The null models are run for each of the 41 or 81 species we analyze separately, using its species-specific estimated dispersal kernel. To generate many samples of the process for each focal analyzed species, under all versions except for ‘Recruits’, a community of species with *identical* dispersal is simulated and different populations of different ‘species’ in the simulation are considered as independent realizations of the null model for the focal analyzed species. We run simulations separately for each analyzed species because interspecific differences in dispersal within a simulation would create differences in fitness and unwarranted selection.

The DL null is calibrated and run separately for each species with two empirical inputs:

a) the overall number of individuals in the model (*J*) – to obtain *J* we compute the density of all individuals in the BCI 50-ha plot (*32*) that have a Diameter at Breast Height (DBH) that equals or exceeds the reproductive threshold of the focal species (*28, 30*), averaged over the surveys from 1985 – 2015. In line with previous works, we use 2/3 of the estimated reproductive threshold (*28, 30*). This density is transformed to *J*, to accommodate the larger area that is simulated.
b) The dispersal distance distribution

Additional empirical inputs that are used in variants of the DL null are mentioned in their descriptions below.

In the simulations, the community consists of a fixed number of *J* individuals on a continuous landscape of 1200 x 1200 meters. In all versions accept for ‘Recruits’, the simulations are initialized by drawing *J* individuals from a species pool of 300 arbitrary equally common species and assigning them a random location. Each time step, a random individual is chosen to die and a new individual is recruited. The new individual is an immigrant from the regional pool with probability *m* = 10/*J* (or 2/*J* in the ‘Fixed’ version) or an offspring of a randomly chosen local individual with probability 1 – *m*. Immigration is incorporated to avoid fixation due to drift. Immigrants land at a random location. Offspring of randomly chosen local individuals disperse from their parent tree a random distance drawn from the dispersal distance distribution in a random direction, with torus boundary conditions (so that seeds dispersing “off the edge” of the landscape appear on the other side). These boundary conditions, as well as simulating a landscape larger than BCI, account for dispersal across the plot boundary from outside of the plot. Species-specific dispersal kernels (selected from the H-2008, H-2001 and LDD options described in the ‘Dispersal Estimates’ section) are used for each species in the simulation. Hence, all species in the simulation are neutral since they have identical dispersal and demography. The simulations are run for 5000 generations (each consisting of *J* time steps) to equilibrate, and then sampled every 10 generations, with 1000 samples overall. Each species-sample combination is treated as an independent sample of a spatial point pattern, and those with abundance similar to the observed species are used for the comparison (see more on abundance binning below). All samples are truncated to an observation window of 1000 x 500 meters, which are the dimensions of the BCI forest plot.

Three additional versions of the null model are used – in the first (‘Lag’), each recruited individual “waits” for *S* time steps before it is recruited, representing the time it takes to grow from seed to adult. We used *S* = *J* for all species since a generation consists of *J* time steps. This incorporates some distancing between individuals due to the lag between dispersal and recruitment to the adult stage, because adults may die before their offsprings, that are often dispersed nearby, reach maturity. Notice that while species differ greatly in their generation time, some growing and dying fast and some growing and dying slowly (*52, 53*), this should not affect our results because our analysis is performed separately for each species and the lag in the simulations is measured in units of the (species-specific) generation time.

The second version (‘Recruits’) considers recruitment to the adult class given the initial distribution observed in the data and the observed demographic rates. Hence, this version requires two additional inputs:

a) The initial spatial distribution of adults of the focal species – we used 1985 as the initial state.
b) The number of recruits (to the adult class) and number of deaths of adults that took place in the time frame of the data (calculated by comparing consecutive surveys).

In the ‘Recruits’ version, we simulated a single species each time on a landscape of 1000 x 500 meters. The ‘Recruits’ simulation is initiated from the observed distribution of adults in 1985 and each realization reshuffles the number of recruitment and mortality events observed over 30 years in a random order. At each time step, a demographic event (adult mortality or recruitment) takes place according to the reshuffling, with recruits dispersing from adult trees. 2∙10^4^ realizations are run, and realizations where a species went extinct are discarded and re-run. Only distances from and densities around new recruits are analyzed. Other versions of the DL null model rely on spatial patterns that emerge over long periods of time in the simulation, while this version refrains from making any assumptions on long-term dynamics, relying on the observed initial distribution. This version also incorporates any demographic trends or large fluctuations that species undergo and that could influence their dispersion.

In the third version (‘Fixed’), we run the simulation using the observed coordinates of all trees of all species with DBH that equals or exceeds the reproductive threshold of the focal species in the 2015 survey. A tree is first chosen to die and is immediately replaced (at the same coordinates) by a recruit. The parent of this recruit is found by drawing a random angle and distance from the replaced tree using the dispersal distance distribution. The tree nearest to the drawn location (using torus boundary conditions on a 1000 x 500 m. landscape) is set to be the parent. This version is set to preserve the distancing of trees from each other that results from competition for space and light.

Overall, we use six different variants of the DL null model analysis: 1) the basic DL model with the standard H-2008 kernel (‘Standard’); 2) The basic DL model with the H-2001 kernel (‘H2001’); 3) the ‘Lag’ DL model with the standard H-2008 kernel; 4) the basic DL model with the kernel incorporating extra long-range dispersal (‘LDD’); 5) the ‘Recruits’ null model, initialized with the observed spatial pattern in 1985, using the standard H-2008 kernel; 6) the ‘Fixed’ null model, initialized with the locations of trees in 2015, using the standard H-2008 kernel. Variants 2-6 of the null model analysis are used as robustness tests.

It is important to note that all our DL models assume that the community has a fixed size. This implicitly incorporates some level of density dependence into the null model expectations. Other DL models may be appropriate in different cases, such as if a species invades an empty habitat.

Previous authors analyzed forest plot data by fitting Poisson cluster models (*26*), interpreting this as ‘dispersal limitation’ (*25, 54*). Poisson cluster models generate spatial distributions by randomly assigning cluster centers according to a (potentially heterogenous) Poisson process, and then distributing individuals within the clusters. These models are static, distributing all individuals at once without considering parents and offspring, and the cluster centers could appear anywhere on the (arbitrary) landscape, eliminating a key element of dispersal limitation. Moreover, observed patterns of aggregation could be due to small-scale spatial heterogeneity (e.g., light gaps) and not dispersal limitation, but the fitted Poisson cluster model would capture them nevertheless. These points emphasize the issues with interpreting the Poisson cluster model as representing dispersal limitation, making us believe that our DL model is much more suitable for this task. On top of that, our DL null is fully parameterized using independent data, thus having zero free parameters to fit.

Spatial statistics

We compared the observed spatial distributions of adults to the null spatial distributions using two analogs of classical spatial statistics: Excess neighborhood Abundance (*EA*(*r*)) and Excess Distance to conspecific neighbors (*ED*), analogs of the pair correlation function and Clark-Evans nearest neighbor statistics (*16, 18, 19, 55*), respectively. All the distances we compute incorporate torus boundary conditions.

Unlike the classical statistics, we design our statistics to be symmetric on the logarithmic scale and have a mean of zero under the null. To compute *EA*(*r*) (eq. 2), the number of conspecific neighbors within an annulus at distances *r* to *r* + Δ*r* is computed for every tree and averaged over all the observed ($N_{obs}(r))$ or simulated (($N_{null}(r))$ trees in the spatial distribution, and the logarithm of both is computed. To obtain the expectation under the DL null model, $\bar{\log\left( N_{null}(r) \right)}$, we perform this calculation for every point pattern sampled with an abundance similar to the observed (see more about abundance binning below), obtaining a distribution of log abundances within each annulus and using its mean over realizations of DL. Moreover, to evaluate the variability of *EA*(*r*) under the null, we construct 95% simulation envelopes by examining the 2.5 and 97.5 percentiles of the distribution of simulated log abundances (*16*). This is presented as an orange area in Fig. 2 and Extended Data Figures 3 - 4. *EA*(*r*) is computed at distance bins of 0 - 10, 10 - 20, 20 - 30, …, 190 – 200 meters. Moreover, to obtain a single overdispersion statistic for each species, we compute *EA*(*r*) for a single circle with radius 20 meters (equivalent to a single distance bin of 0 – 20 m, *EA*(20)), which is often believed to be roughly the radius of interactions (*8, 56, 57*). If no individuals are observed within some distance bin, and to avoid taking the logarithm of zero, we assume that a single tree has a single neighbor at this distance. This equals half of the minimal possible strictly positive number of neighbors, since in real data at least two trees would have each other in this neighborhood (or no trees at all). Values of *EA*(*r*) < 0 indicate a higher abundance than expected at some radius, or aggregation, while *EA*(*r*) < 0 indicates overdispersion.

Notice that because the total number of individuals in the null is matched to the observed, if *EA*(*r*) is negative at small *r*, the trees ‘absent’ at a short distance must appear at (are “repelled” to) larger distances, creating positive *EA*(*r*) at these larger distances. The analogous issue appears with the relative neighborhood density statistic (pair-correlation function) (*16, 18*). Still, these patterns are typically interpreted at short distances (*16, 18, 58*) since it is difficult to interpret patterns at scales larger than roughly half the shortest dimension of the plot (*16*) and since direct effects of trees on each other occur at short distances, through overlapping canopies, root systems or enemies that disperse short distances.

We also study the Excess Distance (*ED*) of trees to their nearest neighbors. For every tree, the distance to the nearest conspecific neighbor is calculated, and the mean ($\bar{NND}_{obs},$sometimes median, ${\hat{\mathrm{NND}}}_{\mathrm{obs}}$, see below) of this distribution over the trees is compared to the expectation under the null, in one of two ways:

(eq. S1)	${ED}_{mean}=\log\left( {\bar{\mathrm{NND}}}_{\mathrm{obs}} \right)-\bar{\log\left( {\bar{\mathrm{NND}}}_{\mathrm{null}} \right)}$,

where ${\bar{\mathrm{NND}}}_{\mathrm{obs}}$ is the observed mean distance of a tree to its nearest neighbor and ${\bar{\mathrm{NND}}}_{\mathrm{null}}$ is this mean for a sample of the null, whose log is averaged over the samples. Alternatively, we use medians instead of means, computing the logarithm of the median distance of a tree to its nearest neighbor, and taking the median over samples of the null:

(eq. S2)	${ED}_{median}=\log\left( {\hat{\mathrm{NND}}}_{\mathrm{obs}} \right)-\hat{log({\hat{\mathrm{NND}}}_{\mathrm{null}})}$,

where “hat” indicates median. Values of *ED* > 0 mean that the distances to neighbors are larger than expected and the distribution is overdispersed, while *ED* < 0 indicates aggregation. To preserve the analogy with the Clark-Evans statistic, we typically used ${ED}_{mean}$ for analyses, accept when using the LDD and Recruits variants of DL. We used medians with the LDD variant because there is an arbitrary proportion of seeds that are dispersed very long distances, and the median is less sensitive to this. For the Recruits variant, we computed only nearest neighbor distances of recruits to all trees, because other trees are preserved under the null. The distribution of nearest neighbor distances in this case is very skewed, and our simulations showed that using the mean (but not the median) would lead to deviations from neutrality when the method is applied to neutral simulations (results not shown). Hence, we used *ED_median_* here as well. Intuitively, if *ED* or *EA*(*r*) have a value of *x*, it indicates that the nearest neighbor distance or number of conspecific neighbors at distance *r*, respectively, are larger by e*^x^* then their expectation, which is the geometric mean (or median) over realizations of the null. For example, the mean *EA*(20) across species is -1.5 under the standard null (Table 1). From eq. 2 of the main text, this indicates $N_{obs}\left( 20 \right)=e^{-1.5}geomean(N_{null}\left( 20 \right))$, so an observed tree $\mathrm{has}$ $(1-e^{-1.5})$∙100% = 78% fewer neighbors than expected based on the null.

The significance of both *EA*(20) and *ED* are evaluated by comparing the logarithm of the observed nearest neighbor distance or abundance with the distribution of their expectations under the nulls with a two-sided test (with *α* = 0.05). Importantly, both nearest neighbor distances and neighborhood density depend strongly on abundance. For this reason, the observed patterns must be compared with samples of similar abundance. Therefore, for all null models except for Recruits (which is initialized with the observed abundance), a species of abundance *N* is compared to null samples with abundances in the range (round(*N* – *B*), round(*N* + *B*)), where *B*, the bin width, is 0.3**N*^0.75^. This non-standard binning meant that a species with abundance < 9 will be compared only to samples of its exact abundance, species with an abundance of 20 will be compared to species with abundance in the range (17,23) and a species with abundance 500 will be compared with the range (468, 532).

Detecting CNDD – general considerations

Our main goal is to detect the spatial signature of CNDD, which should come in the form of repulsion – larger distances and lower densities compared to a sensible null that does not incorporate CNDD. Here we discuss mechanisms and scenarios that can make our analysis too conservative or too liberal.

In general, any mechanism that increases distances between adults (dispersion) and is not incorporated into the null would lead to the false detection of CNDD. To address this possibility, we incorporated several such processes into alternative versions of the null model. Specifically, the time lag between dispersal and recruitment can increase distancing, because the (likely adjacent) parent will often die before recruitment. This mechanism is incorporated into the ‘Lag’ version of the null model. Furthermore, demographic trends or large population fluctuations (beyond the expectations of the drift we incorporated into all null versions) can increase distancing between trees (or decrease it). To address this possibility, the ‘Recruits’ version incorporates the observed number of demographic events, and is initialized with the observed spatial distribution in 1985, therefore making no assumptions about long-term dynamics. The latter property also makes this version weaker, since the information encapsulated in the initial distribution is not used. For similar reasons, underestimating dispersal distance would also lead to false detection of CNDD. To address this, we consider two alternative dispersal kernels, both having longer dispersal distances. All these analyses show qualitatively similar results.

On the other hand, mechanisms that decrease distances between adults and are not incorporated into the null would make our analysis conservative. One such mechanism is spatial habitat specificity, which would generate aggregation and is not incorporated into our null. Other mechanisms include increased dispersal into areas with high adult density due to the attraction of dispersal vectors (*34*), and also clumped seed dispersal (*51*).

Finally, some level of density dependence is incorporated into the null model due to its zero-sum assumption. Even stronger density dependence is incorporated into the ‘Fixed’ version, that preserves the overall spacing between trees without regard to their identity. These assumptions also make our analysis conservative.

Simulation model

We ran simulations to assess the potential degree of overdispersion that results from varying the levels of CNDD and HNDD. To do this we modified our standard null model simulations described above to include density-dependent survival of recruits. We simulate a community of sessile organisms on a landscape with edge length *L* = 600 m and toroidal boundary conditions. The landscape is inhabited by a fixed number of *J* = 5,500 adults. This density corresponds to the density of trees with Diameter at Brest Height (DBH) of 20 cm (the mode and median of the reproductive thresholds of the species we analyzed) and above in the BCI 50-ha forest plot. Each time step, with probability *m* = 10^-3^, a seed arrives from a uniform species pool of *S_reg_* = 300 species at a random location, or, with probability 1 – *m,* a random local individual is chosen to reproduce. The offspring is dispersed with a 2DT (3df) dispersal kernel (*28*). Two mean distances were used: 20 meters, corresponding to the median of the dispersal distance among the species we analyzed, and seven meters, which is close to the minimal distance observed for BCI species. The probability an offspring would establish equals $\frac{1}{1+NCI}$, where *NCI* is the Neighborhood Crowding Index, or the summed contribution of the competitive effects of neighbors, *C_i_*. In line with previous analyses (*57, 59, 60*), we assume that only neighbors within 20 meters have any effect on the focal individual, and for these neighbors the competitive effect of individual *i* at distance *D_i_* on the focal individual is:

(eq. S3)	$C_{i}\left( D_{i} \right)=I_{C_{i}}\frac{Q_{C}}{1+\left( \frac{D_{i}}{b_{1}} \right)^{b_{2}}}+ I_{H_{i}}\frac{Q_{H}}{1+\left( \frac{D_{i}}{b_{1}} \right)^{b_{2}}}$,

where $Q_{C}$and $Q_{H}$ represent the magnitudes of CNDD and HNDD, respectively; $I_{C_{i}}$ and $I_{H_{i}}$ are indicators variables for whether individual *i* is a conspecific or heterospecific (respectively); *b_1_* is the distance at which CNDD is reduced in half, and *b_2_* governs how fast competitive effects decline with distance for both CNDD and HNDD. This functional form enables setting competition to be relatively fixed up to some distance, followed by a sharp decline at longer distances (when *b_2_* is large, see Fig. S10). This causes the effect of density dependence to be quite fixed up to a distance of roughly a crown diameter, followed by a rapid decline. We set *b_1_* = 7 meters, representing a diameter somewhat larger than the crown diameter of adult trees, and b_2_ = 6, so that at 20 meters the effect drops to ~ 0.002Q (see Fig. S10).. If an offspring establishes, it immediately becomes an adult, preserving the species identity of its parent, and a random adult is chosen to die.

The main simulations were run with *b1* = 7 meters all combinations of $Q_{C}$and $Q_{H}$ of 0, 3, 6, 9, 12 each (Results in Figure 3 and S6). Moreover, to examine the special situation when $Q_{C}$ = $Q_{H}$, we ran additional simulations with *b1* = 7 and $Q_{C}$ = $Q_{H}$ = 24, 48, 96 and 192 (Results in Figure S7). To check robustness to the scale of density dependence we further ran simulations with *b_1_* = 20 meters and $Q_{C}$ = $Q_{H}$ = 0, 2, 4, 6, …, 14 (Results in Figure S8), considering the effects of all individuals up to 50 meters. All simulations were initiated as random samples from the pool and run for 1000 generations (*J* timesteps each) to equilibrate, and then sampled –500 - 1000 times, every ten generations. The spatial distribution of all species with more than five individuals was compared to a Dispersal Limitation null model with similar parameters, but where *Q* = 0, using a procedure similar to the empirical analyses. We present the mean *EA*(20) and *ED* across all the species (with more than four individuals) in a simulation. We further analyzed *EA*(*r*) at 10-m distance bins, as in the empirical data, with results presented in Fig. S9, to examine the scale of overdispersion that is generated by density dependence acting at a short scale.

Note that the *m* parameter we chose for the theoretical simulations is lower than values estimated for BCI (*61, 62*), because we wanted to avoid the spatially homogenizing effect of random immigration from outside the plot.

Supplementary Text

Clustered dispersal model analysis

The simulation models we use as nulls for empirical analyses, as well as the models we use for theoretical investigations, all assume that seeds are dispersed independently of each other. However, this assumption is often false: animals congregate in specific areas or bury seeds in caches (*63*), generating clumps of dispersed seeds. How would ignoring this process impact our results, that rely on the independence of seeds?

To address this question, we run and analyze simulation models with clustered seed dispersal. The model considers a single species’ population of size *N_t_* on a continuous toroidal landscape of 600X600 meters with discrete time steps. Every time step, the following events happen:

1. Every tree produces Poisson(*C*) clusters of seeds, where *C* is the expected number of clusters per tree. The cluster centers are dispersed according to the dispersal kernel (2DT with 3DF) with mean distance *D*.
2. Each adult is killed with probability *𝛿*. Hence, an average of *𝛿N_t_* trees die every time step, and a generation constitutes 1/*𝛿* time steps. If the number of adults drawn to be killed exceeds *N_t_*, no adults are killed, preventing the extinction of the population.
3. Poisson(*𝛿N_t_*) seeds are produced. Each seed ‘picks’ a cluster where it appears. The seeds are dispersed around the center of their cluster according to a bivariate gaussian distribution with a mean of zero and SD of *R*, the cluster radius.
4. The dispersed seeds are recruited to the adult stage.
5. If the new number of adults exceeds *J* = 5,500, random adults are killed to cap the population at *J*.

To resemble the likely situation in a tropical forest, we considered the following parameter values. We used mean dispersal distances (*D*) of 7 meters (close to the estimated minimum across species), 20 meters (the median across species) and 60 meters. *𝛿* was set at either 0.02, representing one year (roughly the reported mean yearly mortality rate in Barro Colorado Island, (*63, 64*)), or 0.1, representing five years. Recruitment every several years can be viewed as representing masting. We studied *C* values of 0.02, 0.1 and 1. Larger values would eliminate the ‘clustering’ mechanism, which happens when multiple seeds are recruited in each cluster, i.e. when *𝛿/C* is not small. We considered cluster radii of 1 m., 2.5 m., 5 m. and 10 meters. Values of 2.5-5 meters are the most likely, since Wiegand et al. (*58*) found saplings to have clusters of ~ 2 m. in radius and Detto et al. (*10*) found only some correlation in seedfall between seed traps that are a few meters apart.

Populations were initialized at 100 individuals randomly distributed across the landscape. Models were allowed 1000 generations to equilibrate and then sampled every 10 generations for a total of 4∙10^4^ samples.

The observed populations were compared to a DL null where seeds are dispersed independently of each other, but with a distance distribution equal to what it would be in the clustered simulations if each seed were dispersed independently (rather than in a cluster). Hence, in the DL null, seeds were first dispersed according to a 2DT (3df) kernel with a mean dispersal distance *D*, and then additionally seeds were ‘shifted’ according to a bivariate normal distribution with a SD of *R* and mean of zero. The comparison was made by the same procedure as in other theoretical analyses (using binning by abundance).

Figure S11 presents the *EA*(20) of the simulated populations under the various parameter regimes (examining *ED* reveals similar results). Under all scenarios, the resulting pattern of adults is clustered compared to DL, due to the recruitment of multiple seeds within the same cluster.

This analysis implies that our choice to ignore clumped seed dispersal in the null models we use makes our analysis conservative for detecting overdispersion. Had we incorporated clumped seed dispersal into the DL null, we would have obtained shorter nearest neighbor distances and higher densities under the null, making observed patterns even more overdispersed compared to this hypothetical null.

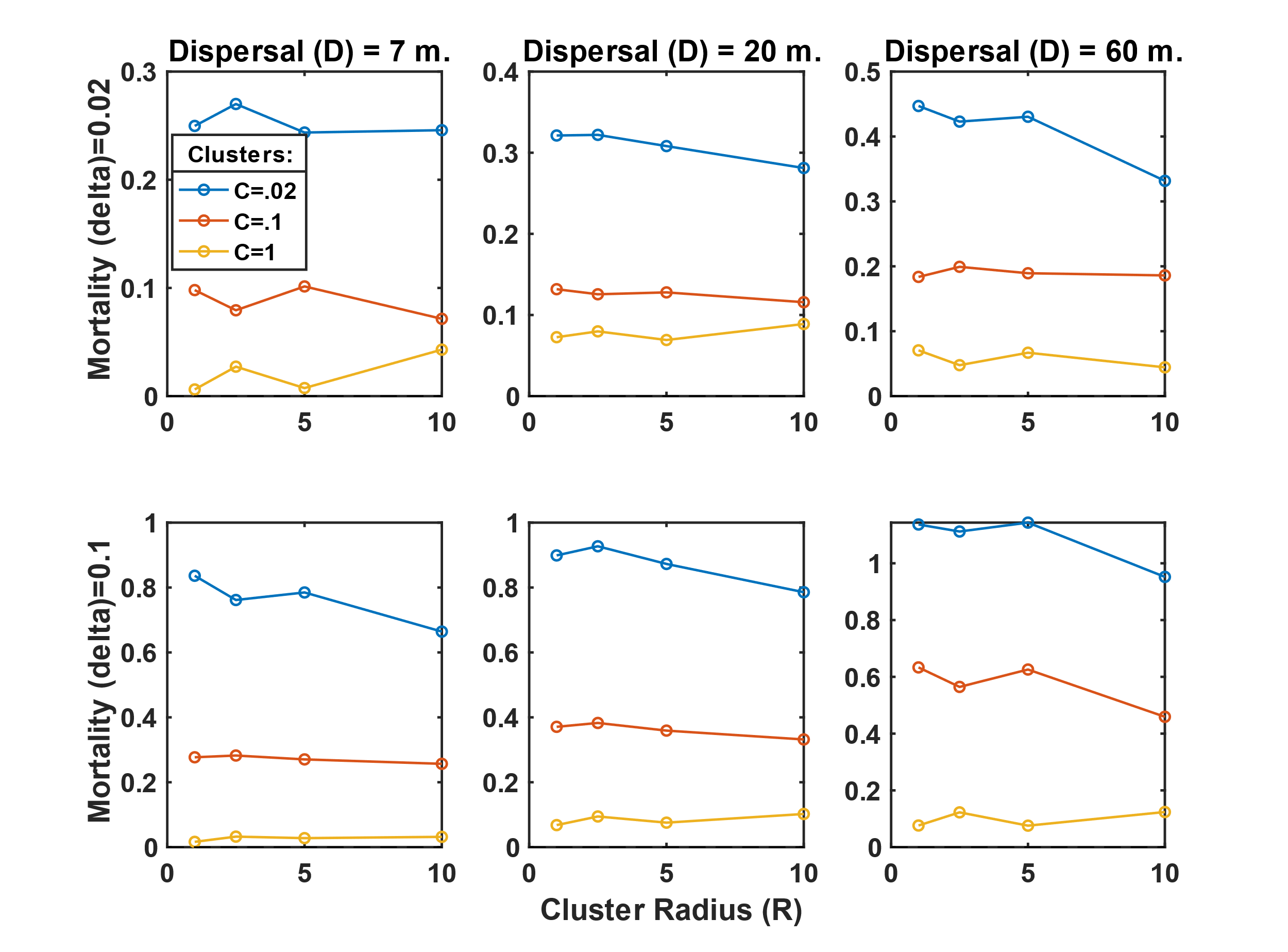

**Fig. S11. Excess neighborhood abundance within 20 meters** (*EA*(20), Y axes) **in simulations with clustered dispersal**. In the simulations, every time step, a proportion *𝛿* of adults is killed, then new adults are recruited from seeds that were dispersed with mean distance *D* into clusters of radius *R*. *C* is the mean number of clusters each tree produces. The resulting populations are compared with DL, so that zero indicates no deviation from DL and positive numbers indicate aggregation compared to DL.

Supplementary Figures

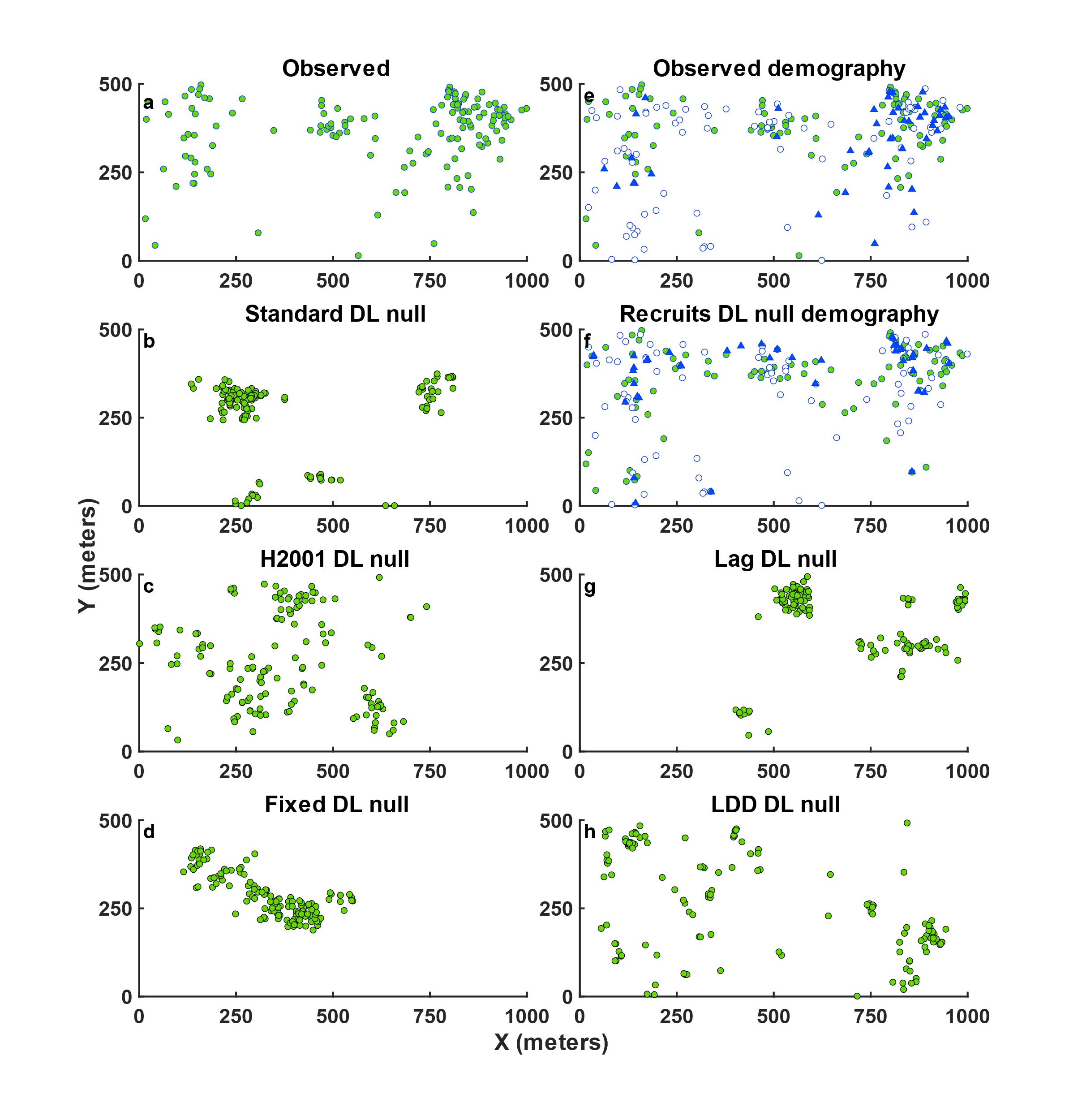

Fig. S1. Spatial point patterns of adult *Beilschmiedia pendula* trees – observed (a, e), compared to realizations of the null models (b-d, f-h). The nulls fix the observed abundance, while the Recruits null fixes the number of observed recruitment (blue triangle) and mortality (empty circle) events from 1985 to 2015.

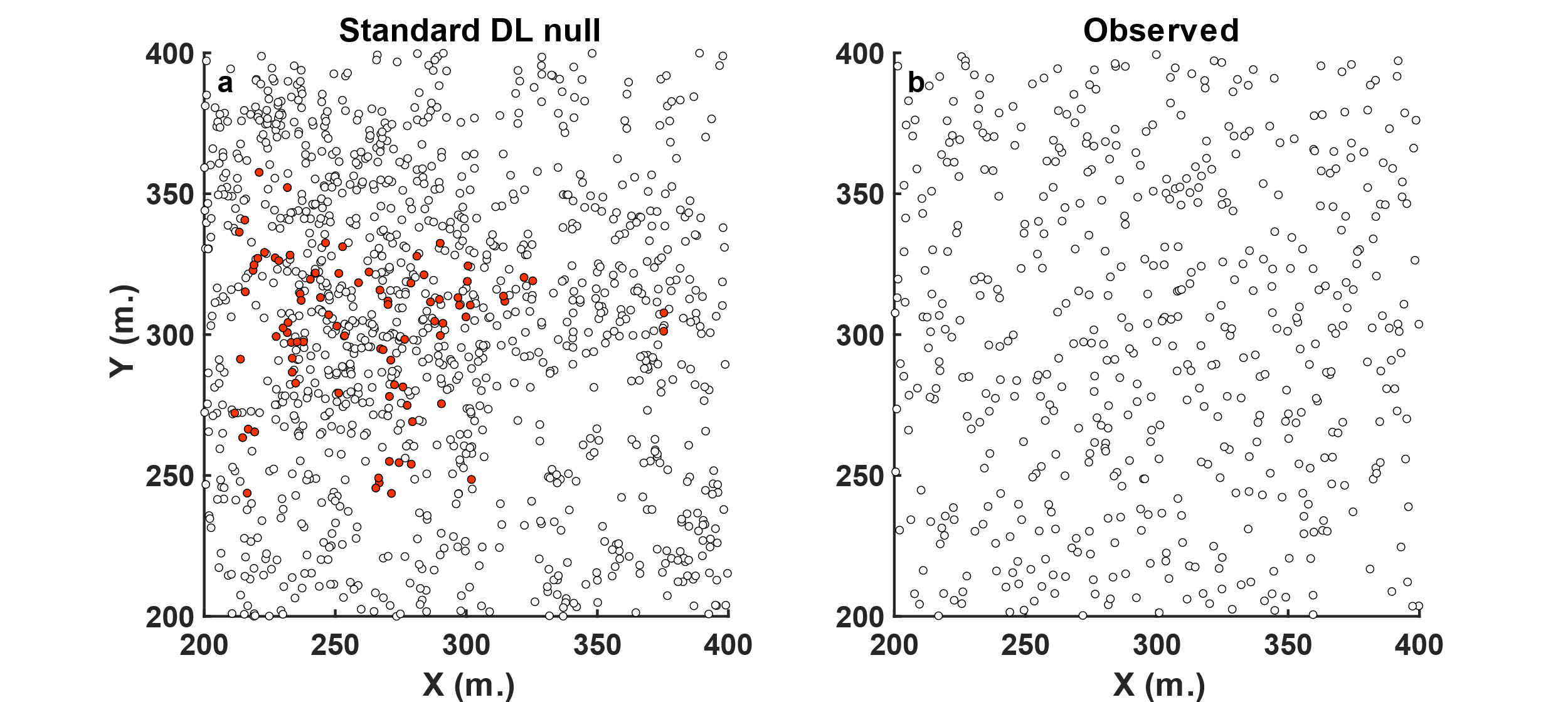

Fig. S2. Observed vs. null spatial patterns, zoomed in. (a) shows a piece of Fig. 1c, zooming on a clump of *Beilschmiedia pendula* trees (red) under the Standard DL null, shown along with heterospecifics (hollow black). This is compared to (b), showing the observed pattern of trees in the same area. The density of trees under the null in this area is somewhat higher than the observed, but it is lower in other parts of t­­he landscape.

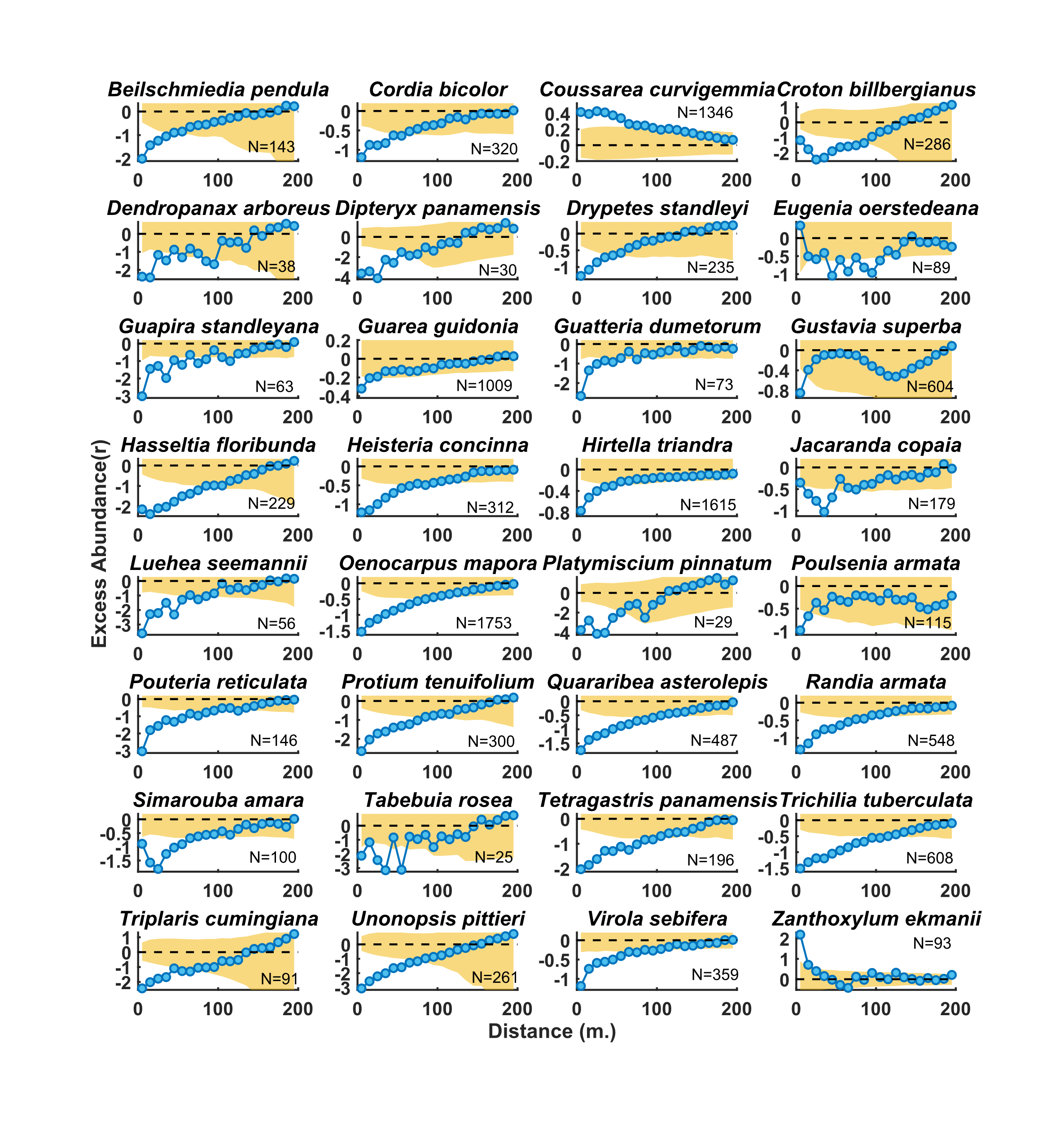
Fig. S3. Excess neighborhood Abundance at distance *r* (*EA*(r)), for all species with data on dispersal distance (in ref. (*28*)) in 2015 with at least 25 individuals. (*N*, shown in each panel). The shaded region represents the 95% simulation envelope of the standard DL null. Note that for less common species, noisy patterns are to be expected. Negative values represent overdispersion w.r.t DL.

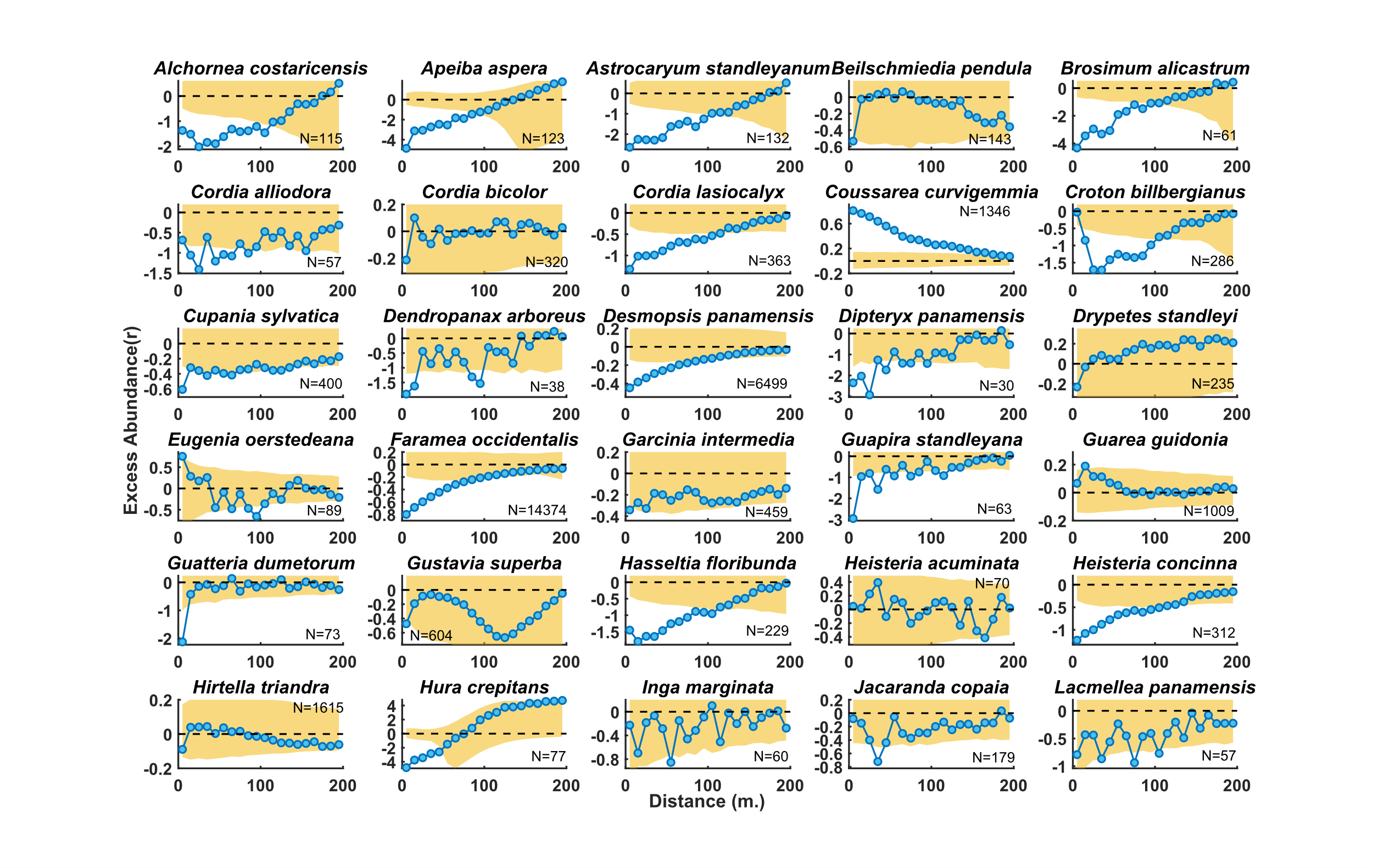

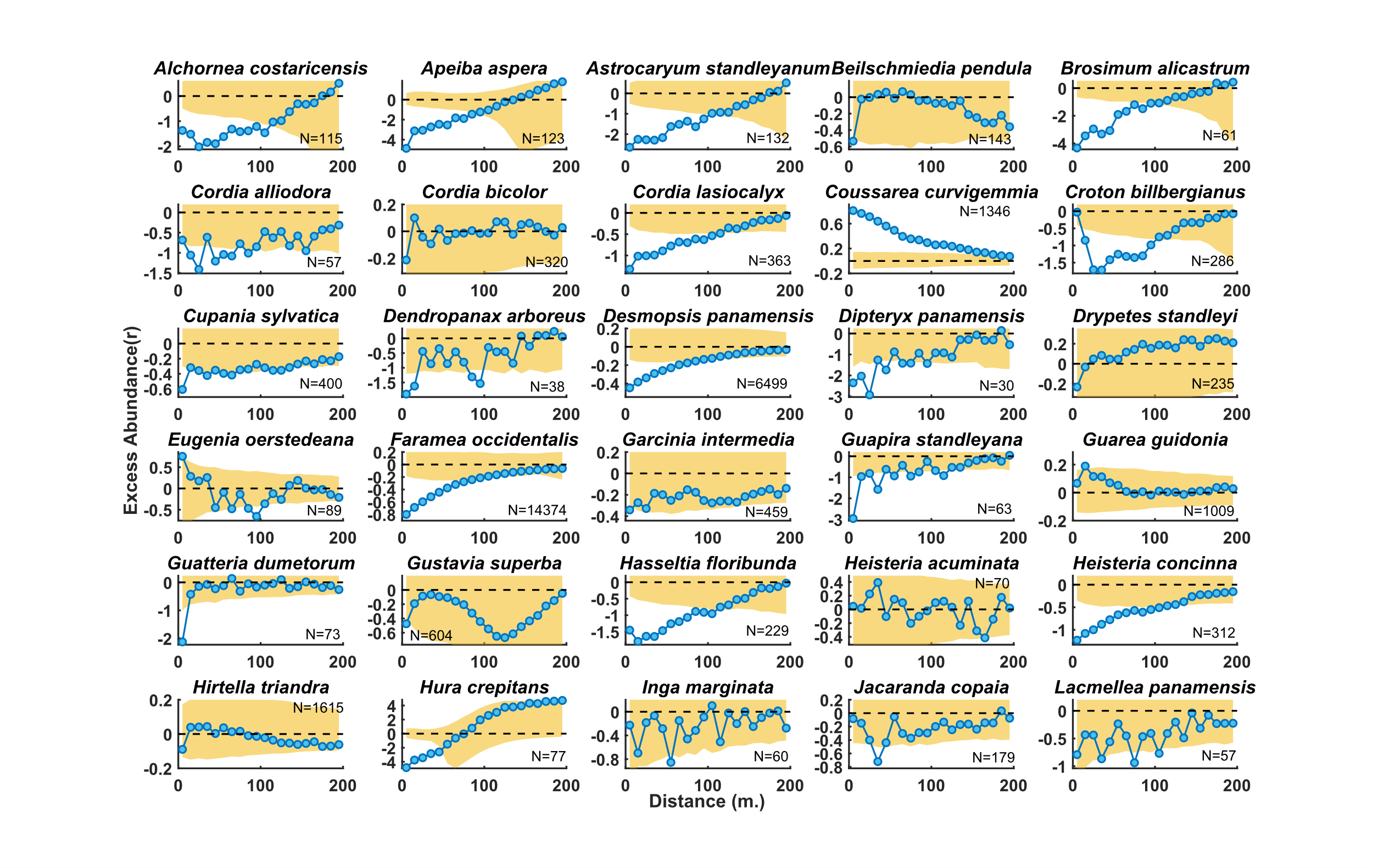

Fig. S4. Excess neighborhood Abundance at distance *r,* *EA*(*r*), for all species with data on dispersal distance (in ref. (*30*)) in 2015 with at least 25 individuals (*N*, shown in each panel). The shaded region represents the 95% simulation envelope of the H2001 DL null. Note that for less common species, noisy patterns are to be expected. Negative values represent overdispersion w.r.t DL.

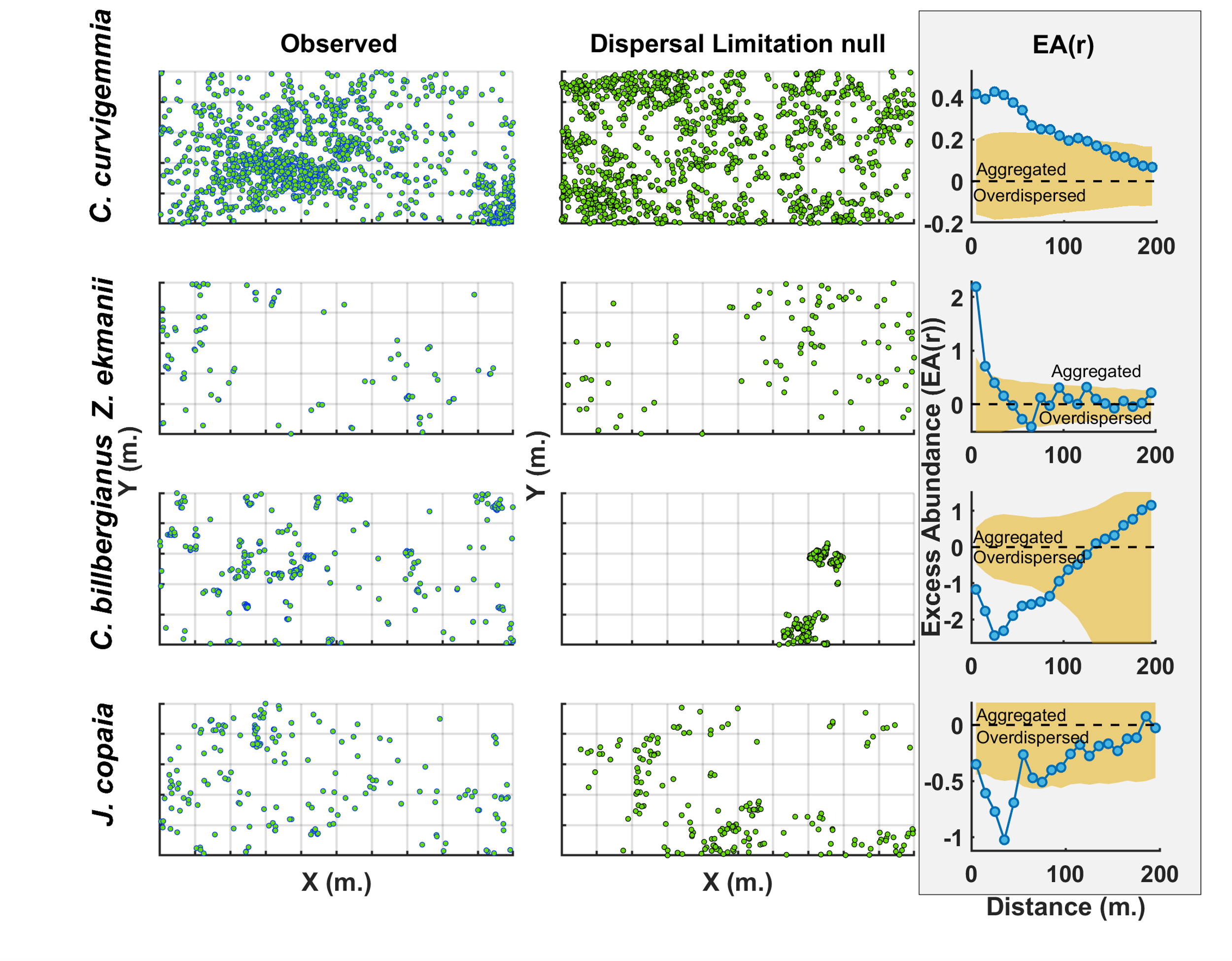

Fig. S5. Spatial distributions of four atypical species and their Excess neighborhood Abundance at distance *r*, *EA*(*r*) (eq. 2). Each row presents a different species, while the columns present for each species the observed distribution (left) versus a single realization of the Standard DL null (center). *EA*(*r*) is presented in the rightmost, shaded column, with the orange region indicating the 95% simulation envelope of the DL null. *EA*(*r*) of 0 corresponds to the expectation of DL while values < 0 indicate overdispersion. All distribution maps are 1000 by 500 meters. *Coussarea curvigemmia* is a habitat specialist with a strong positive affiliation with the large low plateau habitat and negative associations with the large high plateau and slope habitats (*65*), leading to aggregation up to large distances. The three other species, *Zanthoxylum ekmanii*, *Croton billbergianus* and *Jacaranda copaia*, are gap specialists, as is evident from the very high relative growth rate of their fasters-growing juveniles (*53*). They all have increased aggregation at short scales (0 – 30 meters), typical of gaps, compared with intermediate scales. *Z. ekmanii* is also estimated to have a very large dispersal distance (105 meters)(*28*) leading to wide expected dispersion under DL. Compared with this expected dispersion, the observed clumps of trees lead to the high positive values of *EA*(*r*) at small scales for this species.

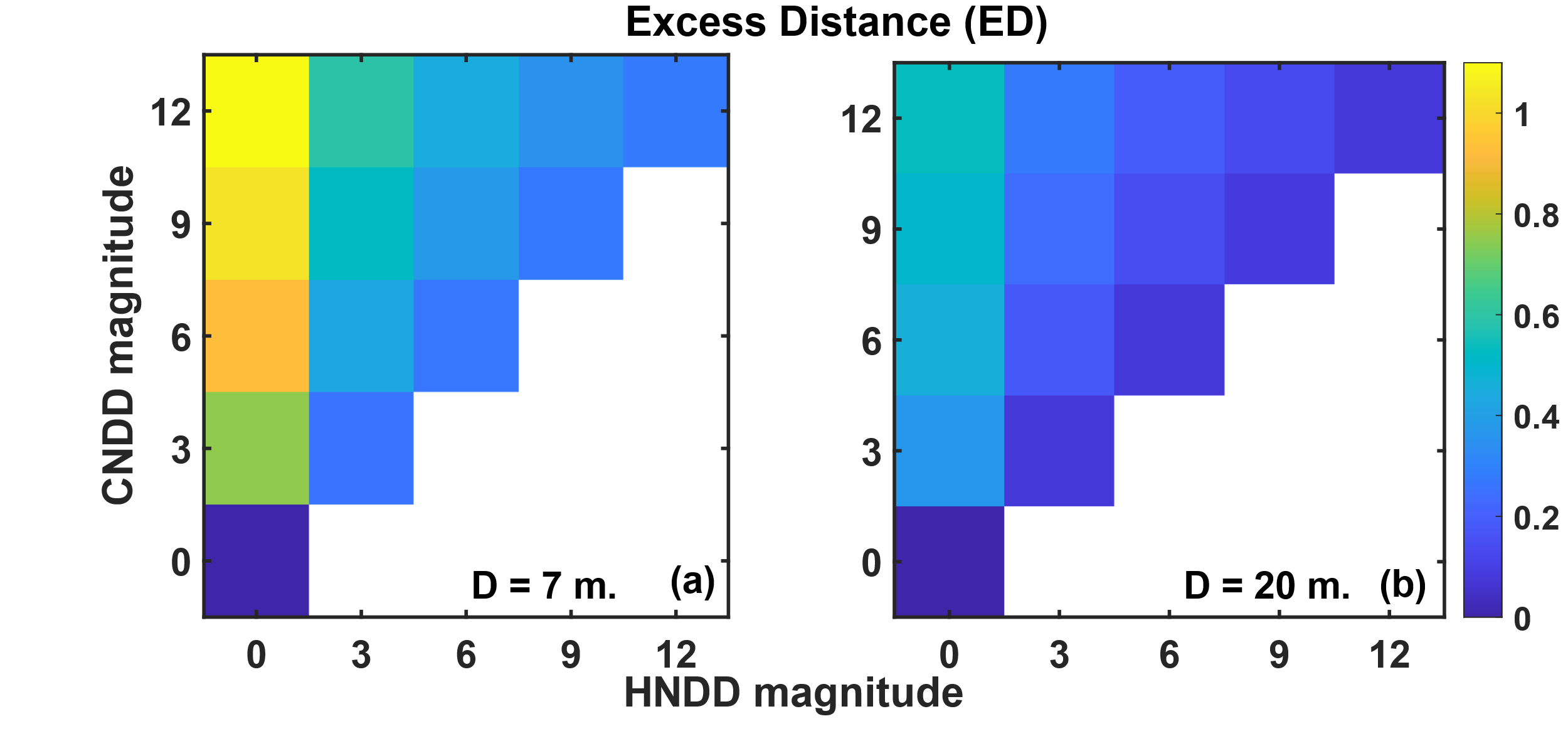

**Fig. S6.** **Excess nearest neighbor Distance** (*ED*) (eq. 1) **in theoretical simulations**. In each simulation, the magnitude of CNDD (*Q_C_*, Y axis) and HNDD (*Q_H_*, X axis) were controlled, and the color of each square represents the mean ED of simulated populations. In the simulations, seeds are dispersed an average distance of *D* = 7 meters (**a**) or *D* = 20 meters (**b**) and the survival of each is determined by both the density of conspecific adults and the density of heterospecific adults in a way that gives large weight to adults within ~ 7 meters of where the seed lands. For perspective, a seed landing right near a single conspecific would have a recruitment probability of 1/(1+*Q_C_*), and more nearby conspecifics and heterospecifics would reduce this probability further. See Methods and eq. S3 for more details. Results are not shown for cases where HNDD > CNDD (below the diagonal) since priority effects lead to one species taking over these communities. When CNDD = HNDD (on the diagonal) overdispersion is weak, and the strongest overdispersion is observed when CNDD >> HNDD.

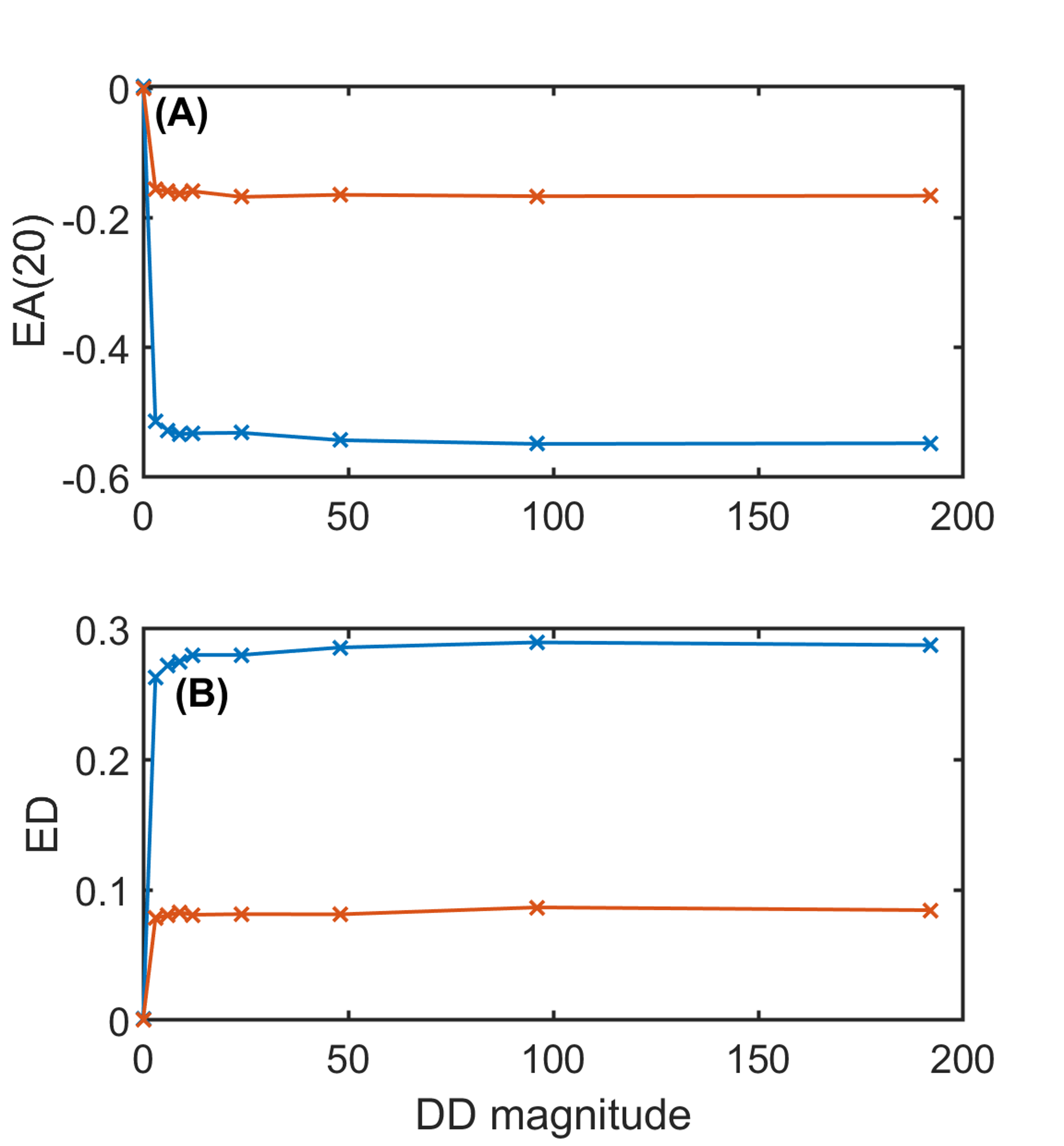

**Fig. S7.** **Spatial statistics in theoretical simulations where CNDD = HNDD = DD magnitude** (X axis) and density dependence operates to ~ 7 meters (*b1* = 7 m.). (**A**) presets the Excess neighborhood abundance within 20 meters (*EA*(20)) while (**B**) presents Excess nearest neighbor distance (ED). In the simulations, seeds are dispersed a distance of 7 meters (blue) or 20 meters (orange) and their survival is determined by the density of trees near the site where they land. See Methods for more details. The overdispersion quickly asymptotes with the magnitude of density dependence, at values considerably lower than observed on BCI (See Table 1).

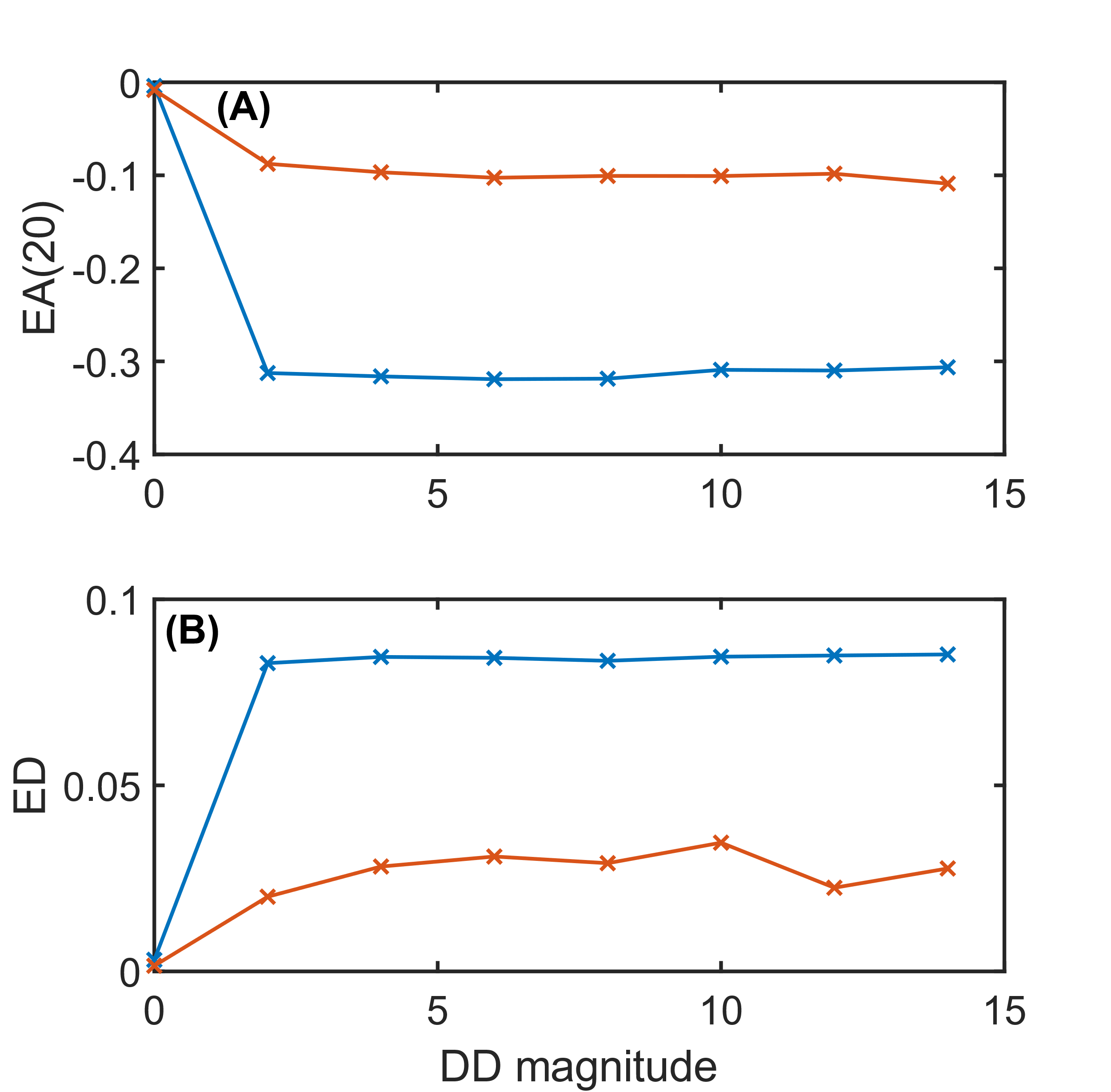

**Fig. S8.** **Spatial statistics in theoretical simulations where CNDD = HNDD = DD magnitude** (X axis) and density dependence operates to ~ 20 meters (*b_1_* = 20 m.). (**A**) presets the Excess neighborhood abundance within 20 meters (*EA*(20)) while (**B**) presents Excess nearest neighbor distance (ED). In the simulations, seeds are dispersed a distance of 7 meters (blue) or 20 meters (orange) and their survival is determined by the density of trees near the site where they land. See Methods for more details. The overdispersion quickly asymptotes with the magnitude of density dependence, at values considerably lower than observed on BCI (See Table 1).

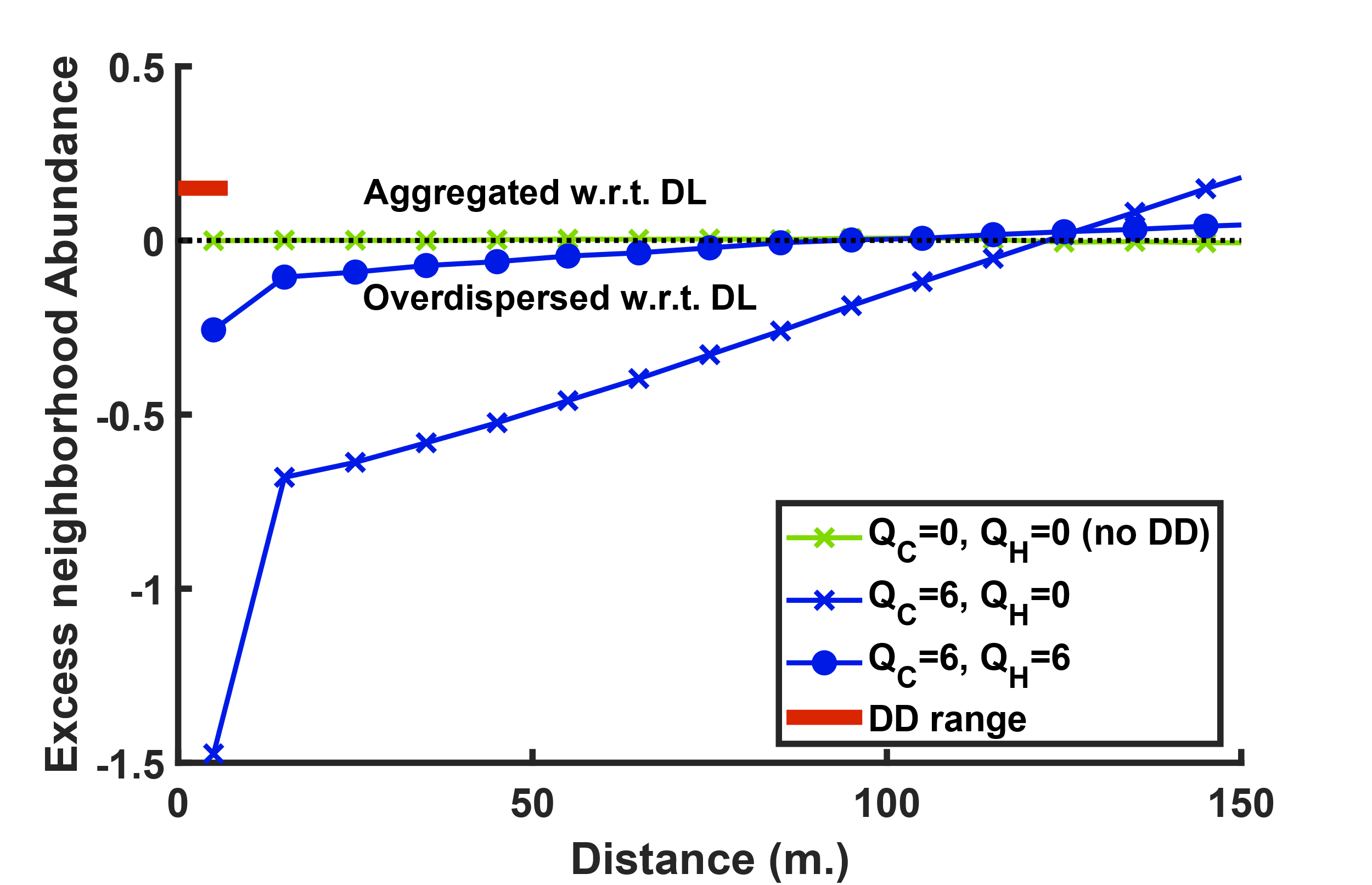

**Fig. S9.** **Excess neighborhood Abundances at different distances *r*** (*EA*(*r*), eq. 1) **in theoretical simulations**. In the simulations presented here, dispersal was set to a mean of 20 meters, the magnitude of CNDD was *Q_C_* = 0 (green) or *Q_C_* = 6 (blue) and the magnitude of HNDD was *Q_H_* = 0 (‘x’) or *Q_H_* = 6 (‘o’). *EA*(*r*) is calculated in ten-meter distance bins. The red line denotes the seven-meter range of density dependence. For more details on the simulation procedure, see method. In the absence of density dependence, *EA*(*r*) is zero for any *r*, while density dependence generates overdispersion up to long distances, even when *Q_C_* = *Q_H_*. The positive values of *EA*(*r*) for large *r*s are an inevitable result of the total number of individuals being fixed in the analyzed pattern and the respective null; hence, if there are fewer individuals than expected at some distance, there must be more than expected at another distance.Density dependence ‘repels’ individuals a large distance away from focal individuals.

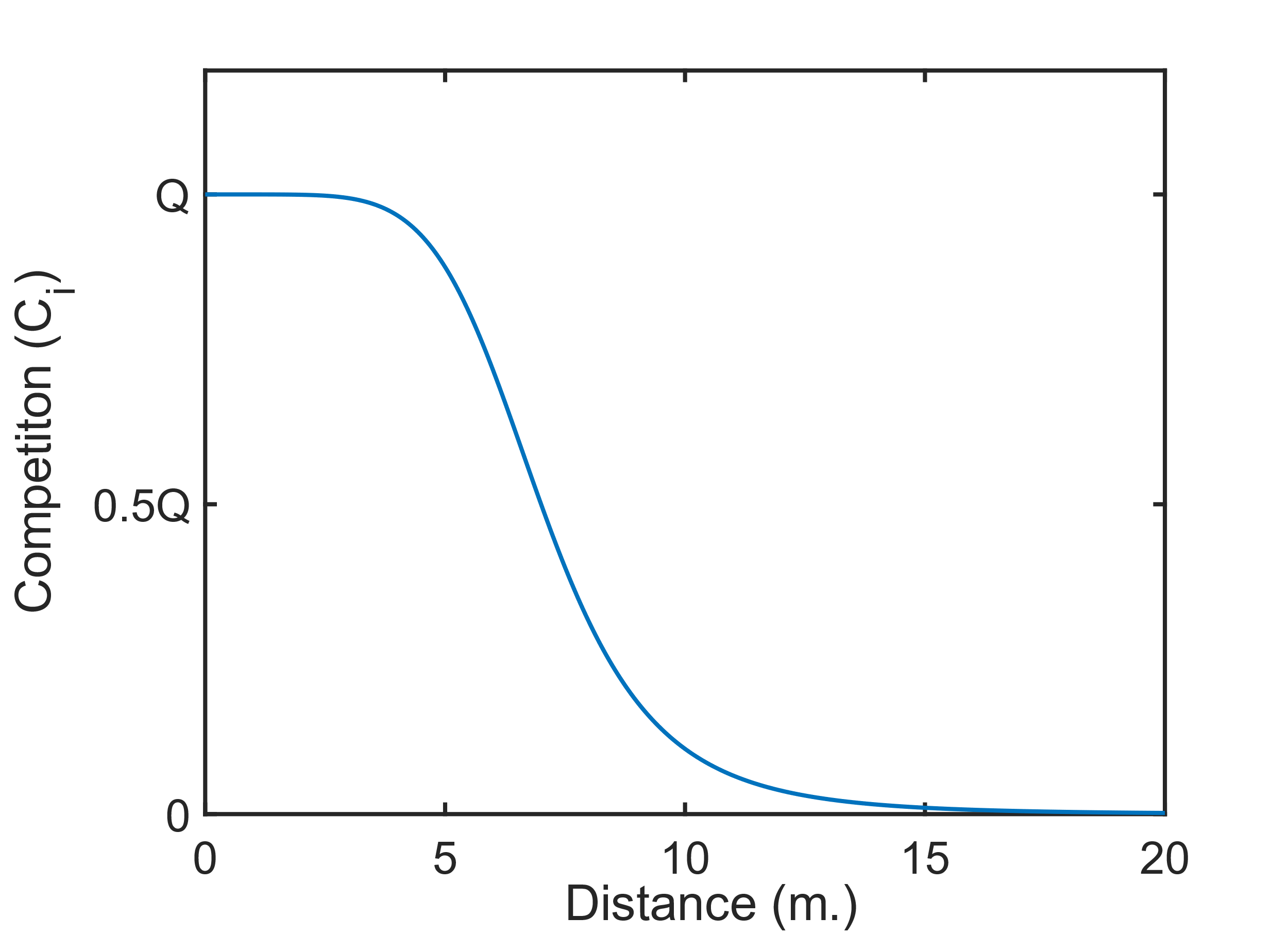

Fig. S10. Competition as a function of distance in the simulation models. Eq. S3 is plotted with *b_1_* = 7 meters and *b_2_* = 6, which were used in most simulations. *Q* represents the maximal magnitude of either CNDD (*Q_C_*) or HNDD (*Q_H_*) since they have similar functional form.

|  | Excess Abundance (EA) at 75-125 meters | | |
| --- | --- | --- | --- |
| Null | Values | Overdispersed | Aggregated |
| Standard | -0.59 ± 0.51 [-2.7, 0.57] | 38 (8) | 3 (1) |
| H2001 | -0.43 ± 0.74 [-2.3, 3.6] | 67 (23) | 14 (3) |
| Lag | -0.49 ± 0.55 [-2.5, 0.46] | 35 (5) | 6 (1) |
| LDD | 0.004 ± 0.5 [-2.3, 0.66] | 12 (2) | 29 (9) |
| Recruits | -0.021 ± 0.22 [-0.59, 0.36] | 14 (6) | 18 (3) |
| Fixed | -0.67 ± 0.57 [-2.9, 0.55] | 38 (13) | 3 (0) |

Table S1. Distributions and significance of Excess Abundance within 75 – 125 meters for Barro Colorado tree species in 2015 using the six variants of the DL null model as references. We compare the abundance of conspecific neighbors within 75 - 125 meters to its expectation on a log scale, hence, negative values represent overdispersion and exponentiating the statistics would give the factor by which the observed density exceeds expectations. The mean ± SD and the range (in square brackets) of EA(75-125) are presented, as well as the number of overdispersed and aggregated species, along with the number of statistically significant (with α = 0.05) results in brackets. Overdispersion is the norm accept in the Recruits and LDD models. The former indicates that recruitment at long distances is comparable to expectations. The latter could be a result of the (arguably) artificially high level of seeds dispersed with no dispersal limitation (10%).
