## Supplementary material for "Pervasive species-specific repulsion among adult tropical trees": Code appendix: Analyze_all_stats3_log.m

```
%%%%%%%%%%%%%%%%%%%%%%%%%%%%%%%%%%%%%%%%%%%%%%%%%%%%%%%%%%%%%%%%%%%%%%%%%%%%%%%%%
% Perform the full analysis of statistics and add them to species data table
%%%%%%%%%%%%%%%%%%%%%%%%%%%%%%%%%%%%%%%%%%%%%%%%%%%%%%%%%%%%%%%%%%%%%%%%%%%%%%%%%
%
% Author: Michael Kalyuzhny
% Date created: 20/01/2021
% Date last modified: 24/01/2022
%
%%%%%%%%%%%%%%%%%%%%%%%%%%%%%%%%%%%%%%%%%%%%%%%%%%%%%%%%%%%%%%%%%%%%%%%%%%%%%%%%%%%%%%%%%%%%


load('species_data1.mat','sp_dat')

all_stats_names = {'R20l_08', 'R20l_08_pval', 'R75_125l_08','R75_125l_08_pval', 'ED_mean_08', 'ED_mean_08_pval', 'ED_med_08', 'ED_med_08_pval', 'null08_N', ...
                   'R20l_01', 'R20l_01_pval', 'R75_125l_01','R75_125l_01_pval', 'ED_mean_01', 'ED_mean_01_pval', 'ED_med_01', 'ED_med_01_pval', 'null01_N', ...
                   'R20l_08_lag', 'R20l_08_lag_pval', 'R75_125l_08_lag','R75_125l_08_lag_pval', 'ED_mean_08_lag', 'ED_mean_08_lag_pval', 'ED_med_08_lag', 'ED_med_08_lag_pval', 'null08_lag_N', ...
                   'R20l_08_LDD', 'R20l_08_LDD_pval', 'R75_125l_08_LDD','R75_125l_08_LDD_pval', 'ED_mean_08_LDD', 'ED_mean_08_LDD_pval', 'ED_med_08_LDD', 'ED_med_08_LDD_pval', 'null08_LDD_N', ...
                   'R20l_08_dem', 'R20l_08_dem_pval', 'R75_125l_08_dem','R75_125l_08_dem_pval', 'ED_mean_08_dem', 'ED_mean_08_dem_pval', 'ED_med_08_dem', 'ED_med_08_dem_pval', ...
                   'R20l_08_fixed', 'R20l_08_fixed_pval', 'R75_125l_08_fixed','R75_125l_08_fixed_pval', 'ED_mean_08_fixed', 'ED_mean_08_fixed_pval', 'ED_med_08_fixed', 'ED_med_08_fixed_pval', 'null08_fixed_N'};
stats = array2table(nan(size(sp_dat,1),length(all_stats_names)),"VariableNames",all_stats_names);

bin_edges = 0:10:200;

% Analyze basic null with 2008 kernel:
disp('Bacis Null')
species_analyze_2008 = find(~isnan(sp_dat{:,'HML2008AlphaFitted'})); %these are the species to run the simulation for
filenames_nulls = [repmat('Results/null1_2008_', length(species_analyze_2008), 1), char(sp_dat{species_analyze_2008,'SpeciesCode'}) repmat('.mat', length(species_analyze_2008), 1) ]; %the filenames of the community samples files to analyze now


[stats{species_analyze_2008, 'R20l_08'}, stats{species_analyze_2008, 'R20l_08_pval'}, stats{species_analyze_2008, 'R75_125l_08'}, stats{species_analyze_2008, 'R75_125l_08_pval'}, ~, ~,  stats{species_analyze_2008, 'ED_mean_08'}, stats{species_analyze_2008, 'ED_mean_08_pval'}, ...
   stats{species_analyze_2008, 'ED_med_08'}, stats{species_analyze_2008, 'ED_med_08_pval'}, stats{species_analyze_2008, 'null08_N'}, dist_bin_centers] = Analyze_nulls3_1(species_analyze_2008, filenames_nulls, 1000, 500, bin_edges, 'Results/Null1_2008_stats.mat',true);

% Null with lag:
disp('Null with lag')
filenames_nulls = [repmat('Results/null2_2008_', length(species_analyze_2008), 1), char(sp_dat{species_analyze_2008,'SpeciesCode'}) repmat('.mat', length(species_analyze_2008), 1) ]; %the filenames of the community samples files to analyze now

[stats{species_analyze_2008, 'R20l_08_lag'}, stats{species_analyze_2008, 'R20l_08_lag_pval'}, stats{species_analyze_2008, 'R75_125l_08_lag'}, stats{species_analyze_2008, 'R75_125l_08_lag_pval'},  ~, ~,  stats{species_analyze_2008, 'ED_mean_08_lag'}, stats{species_analyze_2008, 'ED_mean_08_lag_pval'}, ...
   stats{species_analyze_2008, 'ED_med_08_lag'}, stats{species_analyze_2008, 'ED_med_08_lag_pval'}, stats{species_analyze_2008, 'null08_lag_N'}, ~] = Analyze_nulls3_1(species_analyze_2008, filenames_nulls, 1000, 500, bin_edges, 'Results/Null2_2008_stats.mat',true);

% Null with extra LDD:
disp('Null with LDD')
filenames_nulls = [repmat('Results/null3_2008_', length(species_analyze_2008), 1), char(sp_dat{species_analyze_2008,'SpeciesCode'}) repmat('.mat', length(species_analyze_2008), 1) ]; %the filenames of the community samples files to analyze now

[stats{species_analyze_2008, 'R20l_08_LDD'}, stats{species_analyze_2008, 'R20l_08_LDD_pval'}, stats{species_analyze_2008, 'R75_125l_08_LDD'}, stats{species_analyze_2008, 'R75_125l_08_LDD_pval'}, ~, ~,  stats{species_analyze_2008, 'ED_mean_08_LDD'}, stats{species_analyze_2008, 'ED_mean_08_LDD_pval'}, ...
   stats{species_analyze_2008, 'ED_med_08_LDD'}, stats{species_analyze_2008, 'ED_med_08_LDD_pval'}, stats{species_analyze_2008, 'null08_LDD_N'}, ~] = Analyze_nulls3_1(species_analyze_2008, filenames_nulls, 1000, 500, bin_edges, 'Results/Null3_2008_stats.mat',true);

% Null with 2001 kernel:
disp('Null with 2001 kernel')
species_analyze_2001 = find(~isnan(sp_dat{:,'HML2001Distance'})); %these are the species to run the simulation for
filenames_nulls = [repmat('Results/null1_2001_', length(species_analyze_2001), 1), char(sp_dat{species_analyze_2001,'SpeciesCode'}) repmat('.mat', length(species_analyze_2001), 1) ]; %the filenames of the community samples files to analyze now

[stats{species_analyze_2001, 'R20l_01'}, stats{species_analyze_2001, 'R20l_01_pval'}, stats{species_analyze_2001, 'R75_125l_01'}, stats{species_analyze_2001, 'R75_125l_01_pval'}, ~, ~,  stats{species_analyze_2001, 'ED_mean_01'}, stats{species_analyze_2001, 'ED_mean_01_pval'}, ...
   stats{species_analyze_2001, 'ED_med_01'}, stats{species_analyze_2001, 'ED_med_01_pval'}, stats{species_analyze_2001, 'null01_N'}, ~] = Analyze_nulls3_1(species_analyze_2001, filenames_nulls, 1000, 500, bin_edges, 'Results/Null1_2001_stats.mat',true);

% Null examining recruitment:
disp('Null examining recruitment')
filenames_nulls = [repmat('Results/null4_2008_', length(species_analyze_2008), 1), char(sp_dat{species_analyze_2008,'SpeciesCode'}) repmat('.mat', length(species_analyze_2008), 1) ]; %the filenames of the community samples files to analyze now

[stats{species_analyze_2008, 'R20l_08_dem'}, stats{species_analyze_2008, 'R20l_08_dem_pval'}, stats{species_analyze_2008, 'R75_125l_08_dem'}, stats{species_analyze_2008, 'R75_125l_08_dem_pval'}, ~, ~,  stats{species_analyze_2008, 'ED_mean_08_dem'}, stats{species_analyze_2008, 'ED_mean_08_dem_pval'}, ...
   stats{species_analyze_2008, 'ED_med_08_dem'}, stats{species_analyze_2008, 'ED_med_08_dem_pval'}, ~] = Analyze_nulls_dem3_1(species_analyze_2008, filenames_nulls, 1000, 500, bin_edges, 'Results/Null4_2008_stats.mat');
    
% Null with Fixed trees:
disp('Null with Fixed Trees')
filenames_nulls = [repmat('Results/null6_Fiexd_Trees_2008_', length(species_analyze_2008), 1), char(sp_dat{species_analyze_2008,'SpeciesCode'}) repmat('.mat', length(species_analyze_2008), 1) ]; %the filenames of the community samples files to analyze now

[stats{species_analyze_2008, 'R20l_08_fixed'}, stats{species_analyze_2008, 'R20l_08_fixed_pval'}, stats{species_analyze_2008, 'R75_125l_08_fixed'}, stats{species_analyze_2008, 'R75_125l_08_fixed_pval'}, ~, ~,  stats{species_analyze_2008, 'ED_mean_08_fixed'}, stats{species_analyze_2008, 'ED_mean_08_fixed_pval'}, ...
   stats{species_analyze_2008, 'ED_med_08_fixed'}, stats{species_analyze_2008, 'ED_med_08_fixed_pval'}, stats{species_analyze_2008, 'null08_fixed_N'}, ~] = Analyze_nulls3_1(species_analyze_2008, filenames_nulls, 1000, 500, bin_edges, 'Results/Null6_fixed_trees_2008_stats.mat',false);

% Concatenate tables and save:

save('null_stats.mat','stats') %THIS INTERMEDIATE FILE IS PROVIDED
sp_dat = [sp_dat stats];      
save('species_data3.mat','sp_dat') %THIS INTERMEDIATE FILE IS PROVIDED


%% Short summary:
%load('species_dat3.mat','sp_dat','stats')

clc
disp('Standard:')
nanmean(stats.R75_125l_08)
sum(stats.R75_125l_08_pval<0.05)

disp('HML2001:')
nanmean(stats.R75_125l_01)
sum(stats.R75_125l_01_pval<0.05)

disp('LDD:')
nanmean(stats.R75_125l_08_LDD)
sum(stats.R75_125l_08_LDD_pval<0.05)

disp('Demography:')
nanmean(stats.R75_125l_08_dem)
sum(stats.R75_125l_08_dem_pval<0.05)

disp('Fixed:')
nanmean(stats.R75_125l_08_fixed)
sum(stats.R75_125l_08_fixed_pval<0.05)
```
