## Supplementary material for "Pervasive species-specific repulsion among adult tropical trees": Code appendix: Analyze_nulls_dem3.m

```
%%%%%%%%%%%%%%%%%%%%%%%%%%%%%%%%%%%%%%%%%%%%%%%%%%%%%%%%%%%%%%%%%%%%%%%%%%%%%%%%%
% Compute statistics for a null model for a set of species - with demography
%%%%%%%%%%%%%%%%%%%%%%%%%%%%%%%%%%%%%%%%%%%%%%%%%%%%%%%%%%%%%%%%%%%%%%%%%%%%%%%%%
%
% This version codes the computation for a null that examins new recruits
%
% Author: Michael Kalyuzhny
% Date created: 17/01/2020
% Date last modified: 28/02/2020
%
% Input: 
%   1. Lx, Ly - sizes of landscape
%   2. dist_bind_edges - the left edge of the bins of the Relative Neighborhood density R(r). The last number is the
%   right edge
%   3. species_analyze - the species to run the analysis for
%   4. filenames_nulls - the filenames of the null model runs 
%   5. fileneme_save - filename to save the results
%
% Output:
%    R(20), ED_mean, ED_median and their p values; R(r) and it's 95% simulation envelope; the centers of distance bins
% 
% Procedure:
%   1. Compute the set of statistics for every one of the observed species (each represented by a set of points)
%   2. Import the Null data for that species and compute the distributions of these statistics (including simulation
%   envelopes) for a species with a similar binning
%   3. Compute the normalized statistics
%%%%%%%%%%%%%%%%%%%%%%%%%%%%%%%%%%%%%%%%%%%%%%%%%%%%%%%%%%%%%%%%%%%%%%%%%%%%%%%%%%%%%%%%%%%%

function [R20l, R20l_pval, R75_125l, R75_125l_pval, R, R_sim_envelope, ED_mean, ED_mean_pval, ED_median, ED_median_pval, dist_bin_centers] = Analyze_nulls_dem3(species_analyze, filenames_nulls, Lx, Ly, dist_bin_edges, fileneme_save)

%% Precalculations:

% Temporart version of the input:
load('species_data1.mat','sp_dat')

% % Example input:
% species_analyze = find(~isnan(sp_dat{:,'HML2008AlphaFitted'})); %these are the species to run the simulation for
% filenames_nulls = [ repmat('Results/null4_2008_', length(species_analyze), 1), char(sp_dat{species_analyze,'SpeciesCode'}) repmat('.mat', length(species_analyze), 1) ];
% Lx = 1000; %length (in meters) of the X dimension
% Ly = 500;
% dist_bin_edges = 10:10:200; %'dist_bin_edges' array of the left edge of every bin. last value is the final edge.
% fileneme_save = [];
% species_analyze = species_analyze_2008;

num_sp = length(species_analyze);
num_dist_bins = length(dist_bin_edges) - 1;
dist_bin_centers = dist_bin_edges(1:end-1) + diff(dist_bin_edges)/2; %centers of distance bins

%% Compute observed (raw) statistics:

load('Results/spatial_distributions.mat')

% Empty arrays of results:
R20_raw = nan(num_sp,1);
R75_125_raw = nan(num_sp,1);
R_raw = nan(num_sp, num_dist_bins);
NND_mean_raw = nan(num_sp,1);
NND_median_raw = nan(num_sp,1);

for sp = 1:num_sp
    % data:
    is_new = ~ismember(point_2015{species_analyze(sp)}, point_1985{species_analyze(sp)},'rows');
    new_inds = sum(is_new); %how many new individuals?
    
    points_analyze = [point_2015{species_analyze(sp)}(is_new,:) ; point_2015{species_analyze(sp)}(~is_new,:)]; %reorganize so that the new points are new_inds (at the end):

    %points = point_2015{species_analyze(sp)};
    this_abd = size(points_analyze,1);
    
    % generate distance matrix:
    xdists = abs(points_analyze(:,1)'-points_analyze(1:new_inds,1)); %distance of the first points to every point
    xdists = min(xdists,Lx-xdists); %torus!
    
    ydists = abs(points_analyze(:,2)'-points_analyze(1:new_inds,2)); %distance between every pair of points on a torus along y axis
    ydists = min(ydists,Ly-ydists); %torus!
    
    dd = sqrt(xdists.^2+ydists.^2); %distances of first 'num_samples' individuals to all 

    %replace diagonal with Inf:
    dd(1:(new_inds+1):(new_inds^2)) = Inf;

    %find minimum in every row:
    mins = min(dd,[],2); 
    
    % Calculate neighborhood density:
    dists_binned = nan(new_inds,num_dist_bins); %the thing to average on

    for pp = 1:new_inds
        [dists_binned(pp,:) , ~] = histcounts(dd((pp),:),dist_bin_edges); %for every indivisual, count neighbors within bins of distance
    end
    
    % Compute raw statistics: 
    R20_raw(sp) = log(mean(sum(dd<20))); %number of neighbors within 20 meters
    R75_125_raw(sp) = log(mean(sum( (dd>74)&(dd<125) )));
    R_raw(sp,:) = log(mean(dists_binned,1)); % neighborhood density ~ distance
    NND_mean_raw(sp) = log(mean(mins)); % mean nearest neighbor distance
    NND_median_raw(sp) = log(median(mins)); % median nearest neighbor distance
    
    % Correct Rs that are zeros: assume 1 individual had a neighbor (in real data minimum is 2):
    if R20_raw(sp)==-Inf
        R20_raw(sp) = log(1/this_abd);
    end
    R_raw(sp,~isfinite(R_raw(sp,:))) = log(1/this_abd);
    
    if R75_125_raw(sp)==-Inf
        R75_125_raw(sp) = log(1/this_abd);
    end
end

%% For every species: analyze it's Null and then compute relativized statistics:

% Empty arrays of results:
R20l = nan(num_sp,1);
R20l_pval = nan(num_sp,1);
R75_125l = nan(num_sp,1);
R75_125l_pval = nan(num_sp,1);
R = nan(num_sp, num_dist_bins);
R_sim_envelope = nan(2, num_dist_bins, num_sp); %first dimension: lower/upper, second: distance bin, 3rd: species index
ED_mean = nan(num_sp,1);
ED_mean_pval = nan(num_sp,1);
ED_median = nan(num_sp,1);
ED_median_pval = nan(num_sp,1);

parfor sp = 1:num_sp %run over all species
    sp
    
    % Load null and remove edges:  
    l = load(filenames_nulls(sp,:)); %load simulation results for this species

    num_resamples = length(l.points_final);
    
    % Empty arrays of resamples:
    R20_res = nan(num_resamples,1);
    R75_125_res = nan(num_resamples,1);
    R_res = nan(num_resamples, num_dist_bins);
    NND_mean_res = nan(num_resamples,1);
    NND_median_res = nan(num_resamples,1);
    
    %run over comparable samples in the null model:
    for rr = 1:num_resamples
        
        points_analyze_res = [l.points_final{rr}(l.is_new{rr},:) ; l.points_final{rr}(~l.is_new{rr},:)]; %reorganize so that the new points are first:
        new_inds = sum(l.is_new{rr});
        
        if new_inds>0 %only if any events took place
            % generate distance matrix:
            xdists = abs(points_analyze_res(:,1)'-points_analyze_res(1:new_inds,1)); %distance of the first points to every point
            xdists = min(xdists,Lx-xdists); %torus!

            ydists = abs(points_analyze_res(:,2)'-points_analyze_res(1:new_inds,2)); %distance between every pair of points on a torus along y axis
            ydists = min(ydists,Ly-ydists); %torus!

            dd = sqrt(xdists.^2+ydists.^2); %distances of first 'num_samples' individuals to all 

            %replace diagonal with Inf:
            dd(1:(new_inds+1):(new_inds^2)) = Inf;

            %find minimum in every row:
            mins = min(dd,[],2); 

            % Calculate neighborhood density:
            dists_binned = nan(new_inds,num_dist_bins); %the thing to average on

            for pp = 1:new_inds
                [dists_binned(pp,:) , ~] = histcounts(dd((pp),:),dist_bin_edges); %for every indivisual, count neighbors within bins of distance
            end

            % Compute resampled statistics:
            R20_res(rr) = log(mean(sum(dd<20))); %number of neighbors within 20 meters
            R75_125_res(rr) = log(mean(sum( (dd>75)&(dd<125) )));
            R_res(rr,:) = log(mean(dists_binned,1)); %
            NND_mean_res(rr) = log(mean(mins)); % mean nearest neighbor distance
            NND_median_res(rr) = log(median(mins)); % median nearest neighbor distance
            
            if R20_res(rr)==-Inf
                R20_res(rr) = log(1/this_abd);
            end
            R_res(rr,~isfinite(R_res(rr,:))) = log(1/this_abd);
            
            if R75_125_res(rr)==-Inf
                R75_125_res(rr) = log(1/this_abd);
            end
        end
    end
    
    % Finally, compute all statistics for this species:
    R75_125l(sp) = R75_125_raw(sp) - mean(R75_125_res);
    R75_125l_pval(sp) = dist_pval(R75_125_res, R75_125_raw(sp));
    R20l(sp) = R20_raw(sp) - nanmean(R20_res);
    R20l_pval(sp) = dist_pval(R20_res, R20_raw(sp));
    R(sp,:) = R_raw(sp,:) - nanmean(R_res);
    ED_mean(sp) = NND_mean_raw(sp) - nanmean(NND_mean_res);
    ED_mean_pval(sp) = dist_pval(NND_mean_res, NND_mean_raw(sp));
    ED_median(sp) = NND_median_raw(sp) - nanmedian(NND_median_res);
    ED_median_pval(sp) = dist_pval(NND_median_res, NND_median_raw(sp));

    %Compute dimulation envelopes:
    s = sort(R_res);
    bounds = [round(0.025*num_resamples) round(0.975*num_resamples)]; %the 2.5 and 97.5 percentiles of the resampled densitites
    R_sim_envelope(:,:,sp) = [s(bounds(1),:) - mean(R_res) ; s(bounds(2),:) - mean(R_res)]; %the simulation envelope
end

% Save results:

if ~isempty(fileneme_save)
    save(fileneme_save, 'R20l', 'R20l_pval', 'R75_125l', 'R75_125l_pval', 'R', 'R_sim_envelope', 'ED_mean', 'ED_mean_pval', 'ED_median', 'ED_median_pval')
end

end
```
