## Supplementary material for "Pervasive species-specific repulsion among adult tropical trees": Code appendix: analyze_sim_clustered_dispersal2.m

```
%% Analysis of clustered dispersal model
% Author: Michael Kalyuzhny
% 
% Created: 22/06/22
% 
% Last modified: 13/01/23

%% Preparing the analysis results:
Run_sim_clustered_dispersal1; %In practice - divide into pieces to obtain
%all results

%% Compute stats

% Load references:
load("inp_clustered_null1.mat");
refs = cell(size(inp_null_c));
for dd = 1:length(dist_vals)
    for rr = 1:length(R_vals)
        refs{dd,rr} = load([inp_null_c(dd,rr).output_file '_analysis_results.mat']);
    end
end

load('inp_clustered1.mat')

R_levels = length(R_vals);
delta_levels = length(delta_vals);
C_levels = length(C_vals);
disp_levels = length(dist_vals);

ED = nan(R_levels, C_levels, delta_levels, disp_levels);
EA20 = nan(R_levels, C_levels, delta_levels, disp_levels);

for dd = 1:disp_levels
    for cc = 1:C_levels
        for dede = 1:delta_levels
            for rr = 1:R_levels
               analysis_res_now = inp_all(dd,cc,dede,rr);
               
               if isfile([analysis_res_now.output_file '.mat_analysis.mat']) %if results were obtained
                   analysis_res = load([analysis_res_now.output_file '.mat_analysis.mat']);
                   
                   ED(rr, cc, dede, dd) = nansum((analysis_res.mean_log_dist - refs{dd,rr}.mean_log_dist).*analysis_res.samps_per_bin)/sum(analysis_res.samps_per_bin);
                   EA20(rr, cc, dede, dd) = nansum((analysis_res.log_N20 - refs{dd,rr}.log_N20).*analysis_res.samps_per_bin)/sum(analysis_res.samps_per_bin);

               end
               
            end
        end
    end
end

save('clustered_results.mat','delta_levels', 'R_levels', 'C_levels', 'disp_levels', 'R_vals', 'delta_vals', 'C_vals', 'dist_vals', 'ED', 'EA20') %THIS FILE IS PROVIDED
%% Plot results:

load('clustered_results.mat')

% EA:

figure()

for dede = 1: delta_levels
    for dd = 1: disp_levels
        s1 = subplot(delta_levels, disp_levels, (dede-1)*disp_levels + dd);
        plot(R_vals,EA20(:,:,dede,dd)','-o','LineWidth',1.5)

        line([0 10],[0 0],'LineStyle','--','LineWidth',1,'Color','black')
        
        if dede == 1 %first row
            title(['Dispersal (D) = ' num2str(dist_vals(dd)) ' m.'])
        end
        
        if dd == 1
            ylabel(['Mortality (delta)=' num2str(delta_vals(dede))])
        end
        
        if (dede==2)&(dd==2)
            xlabel('Cluster Radius (R)')
            lgd=legend({'C=.02','C=.1','C=1'});
            lgd.Title.String='Clusters:';
        end
        
        set(s1,'FontSize',16,'FontWeight','bold','LineWidth',1.5);

    end
end
% 
% ED

figure()

for dede = 1: delta_levels
    for dd = 1: disp_levels
        subplot(delta_levels, disp_levels, (dede-1)*disp_levels + dd)
        plot(R_vals,ED(:,:,dede,dd)','-o','LineWidth',1.5)

        line([0 10],[0 0],'LineStyle','--','LineWidth',1,'Color','black')
        
        if dede == 1 %first row
            title(['Dispersal distance (D) = ' num2str(dist_vals(dd)) ' m.'])
        end
        
        if dd == 1
            ylabel(['Mortality (delta)=' num2str(delta_vals(dede))])
        end
        
        if (dede==2)&(dd==2)
            xlabel('Cluster Radius (R)')
            lgd=legend({'C=.02','C=.1','C=1'});
            lgd.Title.String='Clusters:';
        end
    end
end
```
