## Supplementary material for "Pervasive species-specific repulsion among adult tropical trees": Code appendix: Run_sim_clustered_dispersal1.m

```
%%%%%%%%%%%%%%%%%%%%%%%%%%%%%%%%%%%%%%%%%%%%%%%%%%%%%%%%%%%%%%%%%%%
%%% Run simulations for clustered dispersal
%%%%%%%%%%%%%%%%%%%%%%%%%%%%%%%%%%%%%%%%%%%%%%%%%%%%%%%%%%%%%%%%%%%
%
% Author: Michael Kalyuzhny
%
% Date created: 20/06/2022
% Date last modified: 20/06/2022
%
%%%%%%%%%%%%%%%%%%%%%%%%%%%%%%%%%%%%%%%%%%%%%%%%%%%%%%%%%%%%%%%%%%%
% Description: runs the clustered dispersal models and their appropriate nulls

% Dimensions: 1. dispersal distance (a); 2. cluster number (C); 3.
% mortality (delta); 4. Cluster Radius (R)
%
%% Run simultions 

inp_all = struct;

inp_all.L = 600; %landscape edge (meters)
inp_all.b = 2; %'b' parameter of the 2DT kernel (sensu Nathan 2012), = p+1 (sensu Clark 1999), representing heavy tailedness
inp_all.a = 25; %second parameter of the 2DT
inp_all.N0 = 100; %initial number of individuals
inp_all.C = 0.1; %number of seed clusters per adult individual
inp_all.R = 5; %cluster radius
inp_all.delta = 0.02; %death probability
inp_all.N_max = 5500; %maximum abundance

% Time and sampling parameters (time in sweeps):
inp_all.samps_take = 40000; %how many samples to take?
inp_all.samp_freq = 10; %how many generations between samples?
inp_all.first_samp = 1000; %after how many generations to take first sample?
inp_all.print_freq = 10000;
inp_all.output_file = 'sim_clustered.mat';

R_vals = [1 2.5 5 10]; %cluster radii
delta_vals = [0.02 0.1];
C_vals = [0.02 0.1 1];
dist_vals = [7 20 60];

inp_all = repmat(inp_all,length(dist_vals),length(C_vals),length(delta_vals), length(R_vals));

for dd=1:length(dist_vals)
    for cc = 1:length(C_vals)
        for dede = 1:length(delta_vals)
            for rr = 1:length(R_vals)
                inp_all(dd,cc,dede,rr).R = R_vals(rr);
                inp_all(dd,cc,dede,rr).C = C_vals(cc);
                inp_all(dd,cc,dede,rr).delta = delta_vals(dede);
                inp_all(dd,cc,dede,rr).a = dist_vals(dd)*2*gamma(1)/(sqrt(pi)*gamma(0.5)); %set disperdal distance
                inp_all(dd,cc,dede,rr).output_file = ['clustered_disp1_D_' sprintf('%0.4g',dist_vals(dd)) '_C_' sprintf('%0.4g',C_vals(cc)) '_delta_' sprintf('%0.4g',delta_vals(dede)) '_R_' sprintf('%0.4g',R_vals(rr))];
            end
        end
    end
end

save('inp_clustered1.mat','inp_all', 'R_vals', 'delta_vals', 'C_vals', 'dist_vals')

abd_bin_edges = [1 5:2:51 55:5:100 110:10:200 250:50:600 700:100:1000 1250 1500 5500];
dist_bin_edges = 0:10:200;

%% Run:

% I used 'for' instead of 'parfor' since the latter caused a wiered memory leak causing Matlab to crash.
% in practice this can be broken down to pieces to accelerate.

for ii=1:numel(inp_all) 
    inp = inp_all(ii);
    tmp = rand(1,ii); %create differences between simulations with similar seed

    [~, ~] = Sim_clustered(inp);
end

%% Analyze:

parfor ii=1:numel(inp_all)
    filename_load = [inp_all(ii).output_file '.mat'];

    [~, ~, ~, ~, ~, ~, ~, ~, ~, ~, ~, ~] = Stats_regime_clustered_disp(filename_load,1, abd_bin_edges, dist_bin_edges,  5, 600, [filename_load '_analysis.mat']);
end

%% Set the nulls of clustered dispersal:

load('inp_clustered1.mat','inp_all', 'R_vals', 'delta_vals', 'C_vals', 'dist_vals')

% Set general stuff:
inp_null_c = struct;
inp_null_c.J = 5500; %number of individuals in the forest
inp_null_c.L = 600; %landscape edge (meters)
inp_null_c.S_reg = 300; %number of species in regional pool
inp_null_c.b = 2; %'b' parameter of the 2DT kernel (sensu Nathan 2012), = p+1 (sensu Clark 1999), representing heavy tailedness
inp_null_c.imm_prob = 0.001; %number of immigrating seeds to add

% Time and sampling parameters (time in sweeps):
inp_null_c.samps_tot = 5000; %how many samples to take?
inp_null_c.samp_freq = 10; %how many sweeps between samples?
inp_null_c.first_samp = 5000; %after how many sweeps to take first sample?
inp_null_c.print_freq = 1000;

inp_null_c = repmat(inp_null_c,length(dist_vals),length(R_vals));

for dd=1:length(dist_vals)
    for rr = 1:length(R_vals)
        inp_null_c(dd,rr).R = R_vals(rr);        
        inp_null_c(dd,rr).a = dist_vals(dd)*2*gamma(1)/(sqrt(pi)*gamma(0.5)); %set disperdal distance
        inp_null_c(dd,rr).output_file = ['clustered_null1_D_'  sprintf('%0.4g',dist_vals(dd)) '_R_' sprintf('%0.4g',R_vals(rr))];
    end
end

save('inp_clustered_null1.mat','inp_null_c', 'R_vals', 'dist_vals')

%% Run the nulls of clustered dispersal:

parfor ii=1:numel(inp_null_c)
    inp = inp_null_c(ii);
    tmp = rand(1,ii); %create differences between simulations with similar seed

    [~, ~] = sim2_NC(inp);
end

%% Analyze null:

abd_bin_edges = [1 5:2:51 55:5:100 110:10:200 250:50:600 700:100:1000 1250 1500 5500];
dist_bin_edges = 0:10:200;

parfor ii=1:numel(inp_null_c)
    filename_load = [inp_null_c(ii).output_file '.mat'];
    l = load(filename_load);

    Stats_regime3(l.com_samp, abd_bin_edges, dist_bin_edges, 0.25, 5, l.inp.L, [l.inp.output_file '_analysis_results.mat']); %run analysis for this regime and save result separately

end
```
