## Supplementary material for "Pervasive species-specific repulsion among adult tropical trees": Code appendix: sim2_NC.m

```
%%%%%%%%%%%%%%%%%%%%%%%%%%%%%%%%%%%%%%%%%%%%%%%
% Forest simulator - Dispersal Limitation null with TWO STAGED DISPERSAL
%%%%%%%%%%%%%%%%%%%%%%%%%%%%%%%%%%%%%%%%%%%%%%%
%
% Author: Michael Kalyuzhny
%
% First written: 21/08/2020
% Last Modified: 23/09/2022
%
%% Description
%  Main community matrix - 3 columns: x,y,species ID. Seeds are generated
% (with same attributes) and dispersed with torus BC. immigrants are
%  added. All time is in sweeps. Fixed number of samples is taken, after equilibration. 
% Samples are saved in a preset folder and then merged
%
% Input: 'inp' struct with various fields
% output - spatial community 3D array: first dimension is the invividual, second dimension has three elements:
% x,y,species ID, third dimension is the sample. So for example, com_samp(100,:,20) will return the attributes of the
% 100th individual in the 20th sample: [x y species_ID].
%
% THIS VERSION ADDS A SECOND DISPERSAL STAGE

function [com_samp, time_samp] = sim2_NC(inp)

% %% Input patameters - example
% % This can be uncommented when transforming this function into a script.
% 
% inp.J = 1600; %number of individuals in the forest
% inp.L = 200; %landscape edge (meters)
% inp.S_reg = 30; %number of species in regional pool
% inp.b = 2; %'b' parameter of the 2DT kernel (sensu Nathan 2012), = p+1 (sensu Clark 1999), representing heavy tailedness
% inp.a = 25; %second parameter of the 2DT
% inp.imm_prob = 0.01; %number of immigrating seeds to add
% inp.R = 3; %radius of second dispersal 
% 
% % Time and sampling parameters (time in sweeps):
% inp.samps_tot = 100; %how many samples to take?
% inp.samp_freq = 10; %how many sweeps between samples?
% inp.first_samp = 0; %after how many sweeps to take first sample?
% inp.print_freq = 10;
% inp.output_file = 'sim2x_1.mat';

%% Initialization

%all time is in sweeps (=generations, J events)
t_now = 0;
next_samp_time = inp.first_samp;
next_samp_ind = 1; %index of next sample
stop_samp_ind = inp.samps_tot + 1; %when this is the next sample - stop the run
next_print = inp.print_freq;
mkdir(inp.output_file) % create a temporary directory for the samples
time_add = 1/inp.J; %the amount of time (in sweeps) that = a time step
time_samp = NaN(1,inp.samps_tot,'single'); %when were the samples taken?


% Generate initial community
com = zeros(inp.J,3,'single'); %this is the community composition, main dynamic variable. first two columns are coordinates. 3rd is species ID, 4th is quadrat (using linear indexing)
com(:,1:2) = rand(inp.J,2,'single')*inp.L; %set uniform random coordinates
com(:,3) = randi(inp.S_reg,inp.J,1,'single'); %draw species from uniform dist.


%% Main Loop:
tic

while next_samp_ind < stop_samp_ind

    
    if rand > inp.imm_prob % No immigration: generate and disperse seed:
        seed = com(randi(inp.J,'single'),:); %choose a random parent
        u = rand; %temporary variable, to be transformed into distance:
        r = sqrt(((1-u)./(inp.a.^(2*inp.b-2))).^(1./(1-inp.b)) - inp.a^2 ); %transform to 2DT distance distribution
        theta = rand*2*pi; %direction
        seed(1) = seed(1) + r*cos(theta); %perform seed displacement - x coord.
        seed(2) = seed(2) + r*sin(theta); %perform seed displacement - y coord.
        
        seed(1:2) = seed(1:2) + normrnd(0,inp.R,1,2); % ADD SECOND DISPERSAL COMPONENT!

        % Enforce torus boundaries:
        seed(1:2) = rem(seed(1:2),inp.L); %for coordinates > L - performs wrapping, also shortens negative values to be > -L
        seed(seed<0) = inp.L + seed(seed<0); %for negative values
    else % Immigration:
        seed = [rand(1,2,'single')*inp.L randi(inp.S_reg,'single')]; %generate immigrating seeds
    end
    
    com(randi(inp.J,'single'),:) = seed; % replace a random tree with the seed
    
    %print time:
    if t_now >= next_print
        disp(['Runtime: ' sprintf('%0.8g',toc) ' sec., Time: ' sprintf('%0.8g',t_now) ' gen.'])
        next_print = next_print + inp.print_freq;
    end
  
    if t_now >= next_samp_time %if it's time to sample
        time_samp(next_samp_ind) = t_now; %record time
        
        save([inp.output_file '/s_' num2str(next_samp_ind) '_' inp.output_file '.mat'],'com');
        
        %next sampling:
        next_samp_time = next_samp_time + inp.samp_freq;
        next_samp_ind = next_samp_ind + 1;
    end
    
    t_now = t_now + time_add; %progress time
end

% Compose the pieces and save:

com_samp = NaN(inp.J,3,inp.samps_tot,'single');

for ss = 1:inp.samps_tot
    l = load([inp.output_file '/s_' num2str(ss) '_' inp.output_file '.mat'],'com');
    com_samp(:,:,ss) = l.com;
end

save([inp.output_file '.mat'], '-v7.3')
rmdir(inp.output_file, 's') % delete temporary directory for the samples
pause(3); %wait until delete is done, just in case

disp(['Finished! Runtime: ' sprintf('%0.8g',toc) ' sec., Time: ' sprintf('%0.8g',t_now) ' gen.'])

end
```
