## Supplementary material for "Pervasive species-specific repulsion among adult tropical trees": Code appendix: Sim_clustered.m

```
%%%%%%%%%%%%%%%%%%%%%%%%%%%%%%%%%%%%%%%%%%%%%%%
% Clustered dispersal model
%%%%%%%%%%%%%%%%%%%%%%%%%%%%%%%%%%%%%%%%%%%%%%%
%
% Author: Michael Kalyuzhny
%
% Base First written: 21/08/2020
% First written: 13/06/2022
%
%%%%%%%%%%%%%%%%%%%%%%%%%%%%%%%%%%%%%%%%%%%%%% 
%% Description
%
% ######################
%
% The population is represented by a matrix of N by 2 ('pop'), which changes its
% size in real time when the population size changes.
%
%%%%%%%%%%%%%%%%%%%%%%%%%%%%%%%%%%%%%%%%%%%%%%%%%%%
%%
function [pop_samps, pop_size_samps] = Sim_clustered(inp)

% %% Input patameters
% 
% inp.L = 600; %landscape edge (meters)
% inp.b = 2; %'b' parameter of the 2DT kernel (sensu Nathan 2012), = p+1 (sensu Clark 1999), representing heavy tailedness
% inp.a = 25; %second parameter of the 2DT
% inp.N0 = 20; %initial number of individuals
% inp.C = 0.1; %number of seed clusters per adult individual
% inp.R = 5; %cluster radius
% inp.delta = 0.02; %death probability
% inp.N_max = 5500; %maximum abundance
% 
% % Time and sampling parameters (time in sweeps):
% inp.samps_take = 10000; %how many samples to take?
% inp.samp_freq = 100; %how many generations between samples?
% inp.first_samp = 1000; %after how many generations to take first sample?
% inp.print_freq = 10000;
% inp.output_file = 'sim_clustered.mat';

%% Initialization

%all time is in sweeps (J events)
t_now = 0;
time_tot = inp.samps_take*inp.samp_freq + inp.first_samp;
next_samp_time = inp.first_samp;
next_samp_ind = 1; %index of next sample
next_print = inp.print_freq;
pop_samps = cell(1,inp.samps_take);
pop_size_samps = nan(1,inp.samps_take);

% Generate initial population
pop = rand(inp.N0,2,'single')*inp.L; %set uniform random coordinates
pop_size = size(pop,1);


%% Main Loop:
tic

while t_now < time_tot

    % How many clusters does every tree produce?
    clusters_per_tree = poissrnd(inp.C,pop_size,1);
    clusters_tot = sum(clusters_per_tree);
    if clusters_tot == 0 %make sure there is at least one cluster to recruit from
        clusters_tot = 1;
        clusters_per_tree(unidrnd(pop_size)) = 1; %one random tree has a single cluster
    end
    
    % Generate cluster_centers and disperse them:
    
    cluster_centers = repelem(pop,clusters_per_tree,1); %initial location: at the mother tree
            
    u = rand(clusters_tot,1); %temporary variable, to be transformed into distance:
    r = sqrt(((1-u)./(inp.a.^(2*inp.b-2))).^(1./(1-inp.b)) - inp.a^2 ); %transform to 2DT distance distribution
    theta = rand(clusters_tot,1)*2*pi; %direction
    cluster_centers(:,1) = cluster_centers(:,1) + r.*cos(theta); %perform seed displacement - x coord.
    cluster_centers(:,2) = cluster_centers(:,2) + r.*sin(theta); %perform seed displacement - y coord.

    % Kill each adult tree with probability delta
    kill = logical(rand(pop_size,1,'single') < inp.delta);
    
    if sum(kill) >= pop_size
        kill = false(pop_size,1); %if the population will go extinct: don't kill anyone.
    end
    
    pop(kill,:) = [];
    
    % How many should be recruited, and from which clusters?

    to_rec = poissrnd((pop_size*inp.delta)/clusters_tot,clusters_tot,1); %how many recruits per cluster
    total_rec = sum(to_rec);
    
    if total_rec > 0 %anything to recruit
        
        recruits = repelem(cluster_centers,to_rec,1); %generate recruits, pre dispersal within cluster
        recruits = recruits + normrnd(0,inp.R,total_rec,2); %disperse seeds within clusters

        % Enforce torus boundaries:
        recruits(:,1:2) = rem(recruits(:,1:2),inp.L); %for coordinates > L - performs wrapping, also shortens negative values to be > -L
        recruits(recruits<0) = inp.L + recruits(recruits<0); %for negative values

        pop = [pop; recruits];
    end
    
    pop_size = size(pop,1);
    
    if pop_size > inp.N_max %truncate population at N_max
        to_rem_num = pop_size - inp.N_max; %number of individuals to be removed
        
        % Draw (witout replacement) individuals to remove:
        tmp = randperm(pop_size); 
        to_rem = tmp(1:to_rem_num);
        pop(to_rem,:) = [];
        
        pop_size = size(pop,1);
    end
    
    %print time:
    if t_now >= next_print
        disp(['Runtime: ' sprintf('%0.8g',toc) ' sec., Time: ' sprintf('%0.8g',t_now) ' gen.'])
        next_print = next_print + inp.print_freq;
    end
  
    if t_now >= next_samp_time %if it's time to sample
        
        pop_samps{next_samp_ind} = pop;
        pop_size_samps(next_samp_ind) = pop_size;
        
        %next sampling:
        next_samp_time = next_samp_time + inp.samp_freq;
        next_samp_ind = next_samp_ind + 1;
    end
    
    t_now = t_now + inp.delta ; %progress time
end

save([inp.output_file '.mat'], '-v7.3')
disp(['Finished! Runtime: ' sprintf('%0.8g',toc) ' sec., Time: ' sprintf('%0.8g',t_now) ' gen.'])

end
```
