## Supplementary material for "Pervasive species-specific repulsion among adult tropical trees": Code appendix: Stats_regime3.m

```
%%%%%%%%%%%%%%%%%%%%%%%%%%%%%%%%%%%%%%%%%%%%%%%%%%%%%%%%%%%%%%%%%%%%%%%%%%%%%%%%%
% Compute multiple statistics for the samples of a parameter regime
%%%%%%%%%%%%%%%%%%%%%%%%%%%%%%%%%%%%%%%%%%%%%%%%%%%%%%%%%%%%%%%%%%%%%%%%%%%%%%%%%
%
% Author: Michael Kalyuzhny
% Date created: 29/12/2020
% Date last modified: 28/07/2021
%
% Input: 
%       1. 'com_samp' array, where first dimension is the invividual, second dimension has three elements: x,y,species ID, third dimension is the sample.
%       2. 'abd_bin_edges' array - the left edge of each abundance bin. first entry should be one, last should be
%       community size
%       3. 'dist_bin_edges' array of the left edge of every bin. last value is the final edge.
%       4. 'prealloc_size' - fraction of total observations to preallocate bin
%       5. 'min_abd' - the minimal abundance to consider
%       6. 'L' - the edge length of the landscape
%       7. 'filename' - if not NaN, the output will be saved to this location.
%c
% Output:
%       1. 'log_mean_dist' - the mean of the log conspecific nearest neighbor distance in each abundance bin
%       2. 'mean_log_dist - the log of the mean conspecific nearest neighbor distance in each abundance bin
%       3. 'CE' - the mean Clark-Evans statistic in each abundance bin
%       4. 'N20' - the number of neighbors within 20 meters, by abundance bins
%       5. log_N20 the log number of individuals 
%       6. 'ND' - the number of neighbors per distance bin (neighbor distance)
%       7. log_ND - abundnace bins by distance
%       8. 'samps_per_bin' - how many samples were obtained for each abundance bin?
%       9. 'abd_bin_centers' - the centers of abundance bins
%       10. 'dist_bin_centers' - the centers of distance bins
%       11. 'PCF' - the pair correlation function averaged for all species with abundnace above min_abd
%       12. RND20 - relative neighborhood density at 20 meters
%
% Procedure:
%       1. Compute the composition througth time matrix - 'composition_ts', where species are columns and samples (time)
%       are rows
%       2. For every entry that has sufficient abundance, calculate log_dist, CE, N20 and ND. Comments:
%           - ND is a matrix of samples by distance bins
%           - for all others - create a matrix of samples by species and then allocate to bins
%
%%%%%%%%%%%%%%%%%%%%%%%%%%%%%%%%%%%%%%%%%%%%%%%%%%%%%%%%%%%%%%%%%%%%%%%%%%%%%%%%%%%%%%%%%%%%

function [mean_log_dist, log_mean_dist, CE, log_N20, N20, RND20, log_ND, ND, PCF, samps_per_bin, abd_bin_centers, dist_bin_centers] = Stats_regime3(com_samp, abd_bin_edges, dist_bin_edges, prealloc_size, min_abd, L, filename)
% This function computes the statistics for a parameter regime

% % Example input:
% abd_bin_edges = [1 5:40 45:5:100 110:10:200 220:20:400 400:50:750 750:100:1050 1200 1500 5500]; %left edges
% dist_bin_edges = 0:10:200;
% prealloc_size = 0.25; 
% min_abd = 5;
% L = 600;
% filename = 'Stats_regime1_test.mat';

% Precalculations:
dist_bin_centers = dist_bin_edges(1:end-1) + diff(dist_bin_edges)/2; %centers of distance bins
abd_bin_centers = abd_bin_edges(1:end-1) + (diff(abd_bin_edges)-1)/2; %centers of abundance bins
abd_bin_edges_use = abd_bin_edges - 0.01;
abd_bins = length(abd_bin_edges) - 1;
dist_bins = length(dist_bin_edges) - 1;
con_area = pi*dist_bin_edges.^2; %the area of annuli with radii 'dist_bin_edges'
one_over_prop_area = (L^2)./diff(con_area); %what is the proportion of area (out of the total) in each annulus?
one_over_prop_area20 = (L^2)./(pi*400);

% Find all cases when a species is present:
composition_ts = sum_pop(com_samp); % generate a matrix of abundances in samples (rows) by species (columns)
[year,sp] = find(composition_ts>=min_abd); %pairs of year, sp where there are observations
ly = length(year);
ly100= round(ly/100); %one percent of the total calculations to do. Calculated to show progress.

% Preallocate: 
ND_prealloc = zeros(abd_bins, dist_bins); %preallocate neighborhood density at a longer scale
N20_prealloc = nan(round(ly*prealloc_size), abd_bins); %preallocate neighborhood density at one short range
RND20_prealloc = nan(round(ly*prealloc_size), abd_bins); %preallocate rel. neighborhood density at one short range
log_ND_prealloc = zeros(abd_bins, dist_bins); %preallocate neighborhood density at a longer scale
log_N20_prealloc = nan(round(ly*prealloc_size), abd_bins); %preallocate neighborhood density at one short range
nnd_mean_prealloc = nan(round(ly*prealloc_size), abd_bins); %preallocate nearest neighbor distance
PCF_prealloc = nan(round(ly*prealloc_size),dist_bins);

counters = ones(1,abd_bins); %next empty index

%run over all species-year combinations:
disp(['total samples: ' num2str(ly)])
for ii = 1:ly
    
    % Some initial calculations for this species:
    sp_inds = com_samp(:,3,year(ii)) == sp(ii); %find the indexes of this species in this year
    this_abd = sum(sp_inds);
    points = com_samp(sp_inds,1:2,year(ii));
    this_bin = find(abd_bin_edges_use-this_abd>0,1)-1;
    
    % Compute nearest neighbor distances:
    
    xdists = abs(points(:,1)-points(:,1)'); %distance between every pair of points on a torus along x axis
    xdists = min(xdists,L-xdists); %torus!
    ydists = abs(points(:,2)-points(:,2)'); %distance between every pair of points on a torus along y axis
    ydists = min(ydists,L-ydists); %torus!
    dd = sqrt(xdists.^2+ydists.^2); %distances of first 'num_samples' individuals to all 

    %replace diagonal with Inf (so that your distance to yourseld does not get counted in any distance bin):
    dd(1:(this_abd+1):(this_abd^2)) = Inf;

    %find minimum in every row:
    min_dists = min(dd,[],2);
    
    % Calculate neighborhood density:
    dists_binned = nan(this_abd,dist_bins); %the thing to average on

    for pp = 1:this_abd
        [dists_binned(pp,:) , ~] = histcounts(dd((pp),:),dist_bin_edges); %for every indivisual, count neighbors within bins of distance
    end
    
    ND_now = mean(dists_binned,1); %mean neighbors in rings (over individuals)
    log_ND_now = log(ND_now);
    
    %assign to preallocated arrays:
    nnd_mean_prealloc(counters(this_bin),this_bin) = mean(min_dists);
    N20_prealloc(counters(this_bin),this_bin) = mean(sum(dd<20));
    RND20_prealloc(counters(this_bin),this_bin) = N20_prealloc(counters(this_bin),this_bin)*one_over_prop_area20/this_abd;
    ND_prealloc(this_bin,:) = ND_prealloc(this_bin,:) + ND_now;
    log_N20_prealloc(counters(this_bin),this_bin) = log(N20_prealloc(counters(this_bin),this_bin));
    PCF_prealloc(ii,:) = ND_now.*one_over_prop_area/this_abd;
    
    %Correct infinite values when no neighbors are observed:
    if ~isfinite(log_N20_prealloc(counters(this_bin),this_bin))
        log_N20_prealloc(counters(this_bin),this_bin) = log(1/this_abd);
    end
    
    log_ND_now(~isfinite(log_ND_now)) = log(1/this_abd);
    
    log_ND_prealloc(this_bin,:) = log_ND_prealloc(this_bin,:) + log_ND_now;
        
    counters(this_bin) = counters(this_bin) + 1;
    
    if rem(ii,ly100) == 0
        disp(['total samples: ' num2str(ly) ', done: ' num2str(ii)])
    end
end

%Final calculations:
samps_per_bin = counters-1;

mean_log_dist = nanmean(log(nnd_mean_prealloc));
log_mean_dist = log(nanmean(nnd_mean_prealloc));
CE = nanmean((nnd_mean_prealloc))*2.*sqrt(abd_bin_centers/(L^2));
N20 = nanmean(N20_prealloc);
ND = ND_prealloc./(counters-1)';
log_N20 = nanmean(log_N20_prealloc);
log_ND = log_ND_prealloc./(counters-1)';
PCF = nanmean(PCF_prealloc);
RND20 = nanmean(RND20_prealloc);

if ~isnan(filename) %save if required to do so
    save(filename, 'mean_log_dist', 'PCF', 'log_mean_dist', 'CE', 'N20', 'RND20', 'log_N20', 'ND', 'log_ND', 'samps_per_bin', 'abd_bin_centers', 'dist_bin_centers')
end

end
```
