## Supplementary material for "Pervasive species-specific repulsion among adult tropical trees": Code appendix: Analysis_CNDD_HNDD_Delta7.m

```
%%%%%%%%%%%%%%%%%%%%%%%%%%%%%%%%%%%%%%%%%%%%%%%%%%%%%%%%%%%%%%%%%%%
%%% what is the effect of a combination of CNDD and HNDD?
%%%%%%%%%%%%%%%%%%%%%%%%%%%%%%%%%%%%%%%%%%%%%%%%%%%%%%%%%%%%%%%%%%%
%
% Author: Michael Kalyuzhny
%
% Date created: 01/10/2020
% Date last modified: 11/01/2023
%
%%%%%%%%%%%%%%%%%%%%%%%%%%%%%%%%%%%%%%%%%%%%%%%%%%%%%%%%%%%%%%%%%%%
%
% 
% Many of the parameters below, aimed at creating differences beween CNDD and HNDD
% are not used.
%
%% Set main simulations 

inp_all = struct;

inp_all.J = 5500; %number of individuals in the forest
inp_all.L = 600; %landscape edge (meters)
inp_all.S_reg = 300; %number of species in regional pool
inp_all.b = 2; %'b' parameter of the 2DT kernel (sensu Nathan 2012), = p+1 (sensu Clark 1999), representing heavy tailedness
inp_all.a = 25; %second parameter of the 2DT
inp_all.Q_HNDD = 10; %magnitude of HNDD
inp_all.Q_CNDD_min = 10; %minimal value of CNDD
inp_all.Q_CNDD_max = 10;
inp_all.CNDD_dist = 7; %spatial scale at which CNDD drops
inp_all.CNDD_steep = 6; %steepness of CNDD drop with distance
inp_all.H_dist = 7;
inp_all.H_steep = 6;    
inp_all.D = 20;
inp_all.alpha = 1; %total competition
inp_all.imm_prob = 0.001; %number of immigrating seeds to add

% Time and sampling parameters (time in sweeps):
inp_all.samps_tot = 1000; %how many samples to take?
inp_all.samp_freq = 10; %how many sweeps between samples?
inp_all.first_samp = 1000; %after how many sweeps to take first sample?
inp_all.print_freq = 1000;
inp_all.output_file = 'tmp';

magnitude_vals = [0 3 6 9 12];
dist_vals = [7 20];

inp_all = repmat(inp_all,length(magnitude_vals),length(magnitude_vals),length(dist_vals));
%Dimensions: 1. CNDD magnitude; 2. HNDD magnitude; 3. dispersal distance

for cc=1:length(magnitude_vals)
    for hh = 1:length(magnitude_vals)
        for dd = 1:length(dist_vals)
            inp_all(cc,hh,dd).Q_HNDD = magnitude_vals(hh);
            inp_all(cc,hh,dd).Q_CNDD_min = magnitude_vals(cc);
            inp_all(cc,hh,dd).Q_CNDD_max = magnitude_vals(cc);
            inp_all(cc,hh,dd).a = dist_vals(dd)*2*gamma(1)/(sqrt(pi)*gamma(0.5)); %set disperdal distance
            
            inp_all(cc,hh,dd).output_file = ['CNDD_HNDD_QC_'  sprintf('%0.4g',magnitude_vals(cc)) '_QH_' sprintf('%0.4g',magnitude_vals(hh)) '_D_' sprintf('%0.4g',dist_vals(dd))];
        end
    end
end

save('inp_CNDD_HNDD1.mat','inp_all', 'magnitude_vals', 'dist_vals')

%% Run:
parfor cc=1:numel(inp_all)
    inp = inp_all(cc);
    tmp = rand(1,cc); %create differences between simulations with similar seed

    [~, ~, ~] = sim3(inp);
end

%% Set supplementary simultions with CNDD=HNDD

inp_all2 = struct;

inp_all2.J = 5500; %number of individuals in the forest
inp_all2.L = 600; %landscape edge (meters)
inp_all2.S_reg = 300; %number of species in regional pool
inp_all2.b = 2; %'b' parameter of the 2DT kernel (sensu Nathan 2012), = p+1 (sensu Clark 1999), representing heavy tailedness
inp_all2.a = 25; %second parameter of the 2DT
inp_all2.Q_HNDD = 10; %magnitude of HNDD
inp_all2.Q_CNDD_min = 10; %minimal value of CNDD
inp_all2.Q_CNDD_max = 10;
inp_all2.CNDD_dist = 7; %spatial scale at which CNDD drops
inp_all2.CNDD_steep = 6; %steepness of CNDD drop with distance
inp_all2.H_dist = 7;
inp_all2.H_steep = 6;    
inp_all2.D = 20;
inp_all2.alpha = 1; %total co
inp_all2.imm_prob = 0.001; %number of immigrating seeds to add

% Time and sampling parameters (time in sweeps):
inp_all2.samps_tot = 500; %how many samples to take?
inp_all2.samp_freq = 10; %how many sweeps between samples?
inp_all2.first_samp = 1000; %after how many sweeps to take first sample?
inp_all2.print_freq = 1000;
inp_all2.output_file = 'tmp';

magnitude_vals = [24 48 96 192];
dist_vals = [7 20];

inp_all2 = repmat(inp_all2,length(magnitude_vals),length(dist_vals));

for ch=1:length(magnitude_vals)
    for dd = 1:length(dist_vals)
        inp_all2(ch,dd).Q_HNDD = magnitude_vals(ch);
        inp_all2(ch,dd).Q_CNDD_min = magnitude_vals(ch);
        inp_all2(ch,dd).Q_CNDD_max = magnitude_vals(ch);
        inp_all2(ch,dd).a = dist_vals(dd)*2*gamma(1)/(sqrt(pi)*gamma(0.5)); %set disperdal distance

        inp_all2(ch,dd).output_file = ['CNDD_HNDD_QC_'  sprintf('%0.4g',magnitude_vals(ch)) '_QH_' sprintf('%0.4g',magnitude_vals(ch)) '_D_' sprintf('%0.4g',dist_vals(dd))];
    end
end

save('inp_CNDD_HNDD2.mat','inp_all2', 'magnitude_vals', 'dist_vals')

%% Run supplementary simulations:
parfor cc=1:numel(inp_all2)
    inp = inp_all2(cc);
    tmp = rand(1,cc); %create differences between simulations with similar seed

    [~, ~, ~] = sim3(inp);
end

%% Run null:

inp_null = reshape(inp_all(1,1,:),1,2);

for cc = 1:length(dist_vals)
    inp_null(cc).samps_tot = 5000; %how many samples to take?
    inp_null(cc).samp_freq = 10; %how many sweeps between samples?
    inp_null(cc).first_samp = 5000; %after how many sweeps to take first sample?
    inp_null(cc).print_freq = 1000;
    inp_null(cc).output_file = ['ref_HNDD_D_' sprintf('%0.4g',dist_vals(cc))];
end

save('inp_null.mat','inp_null');

parfor cc=1:length(dist_vals)
    [~, ~] = sim2_N(inp_null(cc));
end

%% Analyze null:

% Set analysis parameters:
abd_bin_edges = [1 5:2:51 55:5:100 110:10:200 250:50:600 700:100:1000 1250 1500 5500];
dist_bin_edges = 10:10:200;

parfor cc=1:length(dist_vals)
    l = load(['ref_HNDD_D_' sprintf('%0.4g',dist_vals(cc)) '.mat']);
    Stats_regime3(l.com_samp, abd_bin_edges, dist_bin_edges, 0.25, 5, l.inp.L, [l.inp.output_file '_analysis_results.mat']); %run analysis for this regime and save result separately
end

%% Analyze simulation results:

% Set analysis parameters:
abd_bin_edges = [1 5:2:51 55:5:100 110:10:200 250:50:600 700:100:1000 1250 1500 5500];
dist_bin_edges = 0:10:200;

load('inp_CNDD_HNDD1.mat','inp_all')

parfor cc = 1:numel(inp_all)
    if isfile([inp_all(cc).output_file '.mat'])
        l = load(inp_all(cc).output_file);
        com_samp = l.com_samp;

        Stats_regime3(com_samp, abd_bin_edges, dist_bin_edges, 0.25, 5, inp_all(cc).L, [inp_all(cc).output_file '_analysis_results.mat']); %run analysis for this regime and save result separately
    end
end

%% Analyze complementary simulation results:

% Set analysis parameters:
abd_bin_edges = [1 5:2:51 55:5:100 110:10:200 250:50:600 700:100:1000 1250 1500 5500];
dist_bin_edges = 0:10:200;

load('inp_CNDD_HNDD2.mat','inp_all2')

parfor cc = 1:numel(inp_all2)
    if isfile([inp_all2(cc).output_file '.mat'])
        l = load(inp_all2(cc).output_file);
        com_samp = l.com_samp;

        Stats_regime3(com_samp, abd_bin_edges, dist_bin_edges, 0.25, 5, inp_all2(cc).L, [inp_all2(cc).output_file '_analysis_results.mat']); %run analysis for this regime and save result separately
    end
end
%% Compute stats ED and R20 for main simulations:
load('inp_CNDD_HNDD1.mat')

ref = cell(1,2);
for dd = 1:length(dist_vals)
    ref{dd} = load(['ref_HNDD_D_' sprintf('%0.4g',dist_vals(dd)) '_analysis_results.mat']);
end

ED_HCNDD = nan(size(inp_all));
R20_HCNDD = nan(size(inp_all));
%SRs = nan(size(inp_all));

for cc=1:length(magnitude_vals)
    for hh = 1:length(magnitude_vals)
        for dd = 1:length(dist_vals)  
            
            if isfile([inp_all(cc,hh,dd).output_file '_analysis_results.mat'])
%                 % Compute richenss:
%                 com_samp = load([inp_all(cc,hh,dd).output_file '.mat'],'com_samp');
%                 com_samp = com_samp.com_samp;
%                 pop_t = sum_pop(com_samp);
%                 SRs(cc,hh,dd) = mean(sum(pop_t>0,2));

                % Compute statistics compared to DL:
                reg_now = load([inp_all(cc,hh,dd).output_file '_analysis_results.mat']);
                ED_HCNDD(cc,hh,dd) = nansum((reg_now.mean_log_dist - ref{dd}.mean_log_dist).*reg_now.samps_per_bin)/sum(reg_now.samps_per_bin); %weighted mean
                R20_HCNDD(cc,hh,dd) = nansum((reg_now.log_N20 - ref{dd}.log_N20).*reg_now.samps_per_bin)/sum(reg_now.samps_per_bin); %weighted mean
            end
        end
    end
end

% ALL CASES WHERE HNDD > CNDD LEAD TO PRIORITY EFFECTS AND TO DOMINATION BY A SINGLE SPECIES OCCUPYINH > 97% OF
% INDIVIDUALS

save('HNCC_CNDD.mat','ED_HCNDD','R20_HCNDD', 'magnitude_vals','dist_vals') %THIS FILE IS PROVIDED (and is corrected below)

%% Compute stats ED and R20 FOR complementary simulations:
load('inp_CNDD_HNDD2.mat')

ref = cell(1,2);
for dd = 1:length(dist_vals)
    ref{dd} = load(['ref_HNDD_D_' sprintf('%0.4g',dist_vals(dd)) '_analysis_results.mat']);
end

ED_HCNDD2 = nan(size(inp_all2));
R20_HCNDD2 = nan(size(inp_all2));

for ch=1:length(magnitude_vals)
    for dd = 1:length(dist_vals)
        % Compute statistics compared to DL:
        reg_now = load([inp_all2(ch,dd).output_file '_analysis_results.mat']);
        ED_HCNDD2(ch,dd) = nansum((reg_now.mean_log_dist - ref{dd}.mean_log_dist).*reg_now.samps_per_bin)/sum(reg_now.samps_per_bin); %weighted mean
        R20_HCNDD2(ch,dd) = nansum((reg_now.log_N20 - ref{dd}.log_N20).*reg_now.samps_per_bin)/sum(reg_now.samps_per_bin); %weighted mean
    end
end

save('HNCC_CNDD2.mat','ED_HCNDD2','R20_HCNDD2', 'magnitude_vals','dist_vals') %THIS FILE IS PROVIDED

%% Eliminate everything above the diagonal:

load('HNCC_CNDD.mat','ED_HCNDD','R20_HCNDD', 'magnitude_vals','dist_vals')

for cc=1:length(magnitude_vals)
    for hh = (cc+1):length(magnitude_vals)
        for dd = 1:length(dist_vals)  
                ED_HCNDD(cc,hh,dd) = nan;
                R20_HCNDD(cc,hh,dd) = nan;
        end
    end
end

save('HNCC_CNDD.mat','ED_HCNDD','R20_HCNDD', 'magnitude_vals','dist_vals')

%% Main text figure:

load('HNCC_CNDD.mat','ED_HCNDD','R20_HCNDD', 'magnitude_vals','dist_vals')


figure()
s1 = subplot(1,2,1);
h = imagesc(R20_HCNDD(5:-1:1,:,1), [-1.6 0]); 
set(h,'AlphaData',~isnan(R20_HCNDD(5:-1:1,:,1)));
colormap parula
set(s1,'FontSize',16,'FontWeight','bold','LineWidth',1.5);
ylabel('CNDD magnitude')
xlabel('HNDD magnitude')
xticklabels({'0','3','6','9', '12'})
xticks([1 2 3 4 5])
yticklabels({'12','9','6','3', '0'})
title('Excess neighborhood Abundance (EA(20))')
text(4.2,0.7,'(a)','FontSize',16,'FontWeight','bold')
text(3.6,1.1,'D = 7 m.','FontSize',16,'FontWeight','bold')

s1 = subplot(1,2,2);
h = imagesc(R20_HCNDD(5:-1:1,:,2), [-1.6 0]);
set(h,'AlphaData',~isnan(R20_HCNDD(5:-1:1,:,2)));
colormap parula
set(s1,'FontSize',16,'FontWeight','bold','LineWidth',1.5);
xticklabels({'0','3','6','9', '12'})
xticks([1 2 3 4 5])
yticklabels({'12','9','6','3', '0'})
text(4.2,0.7,'(b)','FontSize',16,'FontWeight','bold')
text(3.6,1.1,'D = 20 m.','FontSize',16,'FontWeight','bold')
colorbar

%% Supplamentary Figure:
figure()
s1 = subplot(1,2,1);
h = imagesc(ED_HCNDD(5:-1:1,:,1), [0 1.1]); 
set(h,'AlphaData',~isnan(ED_HCNDD(5:-1:1,:,1)));
colormap parula
set(s1,'FontSize',16,'FontWeight','bold','LineWidth',1.5);
ylabel('CNDD magnitude')
xlabel('HNDD magnitude')
xticklabels({'0','3','6','9', '12'})
xticks([1 2 3 4 5])
yticklabels({'12','9','6','3', '0'})
title('Excess Distance (ED)')
text(4.2,0.7,'(a)','FontSize',16,'FontWeight','bold')
text(3.6,1.1,'D = 7 m.','FontSize',16,'FontWeight','bold')

s1 = subplot(1,2,2);
h = imagesc(ED_HCNDD(5:-1:1,:,2), [0 1.1]);
set(h,'AlphaData',~isnan(ED_HCNDD(5:-1:1,:,2)));
colormap parula
set(s1,'FontSize',16,'FontWeight','bold','LineWidth',1.5);
xticklabels({'0','3','6','9', '12'})
xticks([1 2 3 4 5])
yticklabels({'12','9','6','3', '0'})
text(4.2,0.7,'(b)','FontSize',16,'FontWeight','bold')
text(3.6,1.1,'D = 20 m.','FontSize',16,'FontWeight','bold')
colorbar

%% Merge data from two simulation sets:

set1 = load('HNCC_CNDD.mat','ED_HCNDD','R20_HCNDD', 'magnitude_vals','dist_vals');
set2 = load('HNCC_CNDD2.mat');

all_mags = [set1.magnitude_vals set2.magnitude_vals];
all_R20 = [diag(set1.R20_HCNDD(:,:,1))' set2.R20_HCNDD2(:,1)' ;...
    diag(set1.R20_HCNDD(:,:,2))' set2.R20_HCNDD2(:,2)']; %first row: D = 7, second: D = 20
all_ED = [diag(set1.ED_HCNDD(:,:,1))' set2.ED_HCNDD2(:,1)' ;...
    diag(set1.ED_HCNDD(:,:,2))' set2.ED_HCNDD2(:,2)']; %first row: D = 7, second: D = 20

%% Supplementary figure for CNDD = HNDD:

figure()
s1 = subplot(2,1,1);
plot(all_mags,all_R20,'-x','LineWidth',1.5,'MarkerSize',8)
ylabel('EA(20)')
xlabel('DD magnitude')
set(s1,'FontSize',16,'LineWidth',1.5);
text(0.2,-0.05,'(A)','FontSize',16,'FontWeight','bold')

s1 = subplot(2,1,2);
plot(all_mags,all_ED,'-x','LineWidth',1.5,'MarkerSize',8)
ylabel('ED')
xlabel('DD magnitude')
text(0.2,0.2,'(B)','FontSize',16,'FontWeight','bold')
set(s1,'FontSize',16,'LineWidth',1.5);
```
