## Supplementary material for "Pervasive species-specific repulsion among adult tropical trees": Code appendix: Analysis_CNDD_HNDD_Delta20.m

```
%%%%%%%%%%%%%%%%%%%%%%%%%%%%%%%%%%%%%%%%%%%%%%%%%%%%%%%%%%%%%%%%%%%
%%% what is the effect of a combination of CNDD and HNDD?
%%%%%%%%%%%%%%%%%%%%%%%%%%%%%%%%%%%%%%%%%%%%%%%%%%%%%%%%%%%%%%%%%%%
%
% Author: Michael Kalyuzhny
%
% Date created: 01/10/2020
% Date last modified: 21/10/2022
%
%%%%%%%%%%%%%%%%%%%%%%%%%%%%%%%%%%%%%%%%%%%%%%%%%%%%%%%%%%%%%%%%%%%
%
% 
% Many of the parameters below, aimed at creating differences beween CNDD and HNDD
% are not used.

% Version 3 considers a case when CNDD = HNDD and both operate and 20
% meters
%

%% Run supplementary simultions with CNDD=HNDD

inp_all3 = struct;

inp_all3.J = 5500; %number of individuals in the forest
inp_all3.L = 600; %landscape edge (meters)
inp_all3.S_reg = 300; %number of species in regional pool
inp_all3.b = 2; %'b' parameter of the 2DT kernel (sensu Nathan 2012), = p+1 (sensu Clark 1999), representing heavy tailedness
inp_all3.a = 25; %second parameter of the 2DT
inp_all3.Q_HNDD = 10; %magnitude of HNDD
inp_all3.Q_CNDD_min = 10; %minimal value of CNDD
inp_all3.Q_CNDD_max = 10;
inp_all3.CNDD_dist = 20; %spatial scale at which CNDD drops
inp_all3.CNDD_steep = 6; %steepness of CNDD drop with distance
inp_all3.H_dist = 20;
inp_all3.H_steep = 6;    
inp_all3.D = 50;
inp_all3.alpha = 1; %total co
inp_all3.imm_prob = 0.001; %number of immigrating seeds to add

% Time and sampling parameters (time in sweeps):
inp_all3.samps_tot = 500; %how many samples to take?
inp_all3.samp_freq = 10; %how many sweeps between samples?
inp_all3.first_samp = 1000; %after how many sweeps to take first sample?
inp_all3.print_freq = 200;
inp_all3.output_file = 'tmp';

magnitude_vals = [0 2 4 6 8 10 12 14];
dist_vals = [7 20];

inp_all3 = repmat(inp_all3,length(magnitude_vals),length(dist_vals));

for ch=1:length(magnitude_vals)
    for hh = 1:length(magnitude_vals)
        for dd = 1:length(dist_vals)
            inp_all3(ch,dd).Q_HNDD = magnitude_vals(ch);
            inp_all3(ch,dd).Q_CNDD_min = magnitude_vals(ch);
            inp_all3(ch,dd).Q_CNDD_max = magnitude_vals(ch);
            inp_all3(ch,dd).a = dist_vals(dd)*2*gamma(1)/(sqrt(pi)*gamma(0.5)); %set disperdal distance
            
            inp_all3(ch,dd).output_file = ['CNDD_HNDD_QC_'  sprintf('%0.4g',magnitude_vals(ch)) '_QH_' sprintf('%0.4g',magnitude_vals(ch)) '_D_' sprintf('%0.4g',dist_vals(dd)) '_Delta_20'];
        end
    end
end

save('inp_CNDD_HNDD3.mat','inp_all3', 'magnitude_vals', 'dist_vals')

%% Run supplementary simulations:
parfor cc=1:numel(inp_all3)
    inp = inp_all3(cc);
    tmp = rand(1,cc); %create differences between simulations with similar seed

    [~, ~, ~] = sim3(inp);
end

%% Run null:

% THIS CAN BE OMITTED IF ALREADY DONE BEFORE

inp_null = inp_all(:,1);

for cc = 1:length(dist_vals)
    inp_null(cc).samps_tot = 5000; %how many samples to take?
    inp_null(cc).samp_freq = 10; %how many sweeps between samples?
    inp_null(cc).first_samp = 5000; %after how many sweeps to take first sample?
    inp_null(cc).print_freq = 1000;
    inp_null(cc).output_file = ['ref_HNDD_D_' sprintf('%0.4g',dist_vals(cc))];
end

save('inp_null.mat','inp_null');

parfor cc=1:length(dist_vals)
    [~, ~] = sim2_N(inp_null(cc));
end

%% Analyze null:

% Set analysis parameters:
abd_bin_edges = [1 5:2:51 55:5:100 110:10:200 250:50:600 700:100:1000 1250 1500 5500];
dist_bin_edges = 10:10:200;

parfor cc=1:length(dist_vals)
    l = load(['ref_HNDD_D_' sprintf('%0.4g',dist_vals(cc)) '.mat']);
    Stats_regime3(l.com_samp, abd_bin_edges, dist_bin_edges, 0.25, 5, l.inp.L, [l.inp.output_file '_analysis_results.mat']); %run analysis for this regime and save result separately
end

%% Analyze simulation results:

% Set analysis parameters:
abd_bin_edges = [1 5:2:51 55:5:100 110:10:200 250:50:600 700:100:1000 1250 1500 5500];
dist_bin_edges = 0:10:200;

load('inp_CNDD_HNDD3.mat','inp_all3')

parfor cc = 1:numel(inp_all3)
    if isfile([inp_all3(cc).output_file '.mat'])
        l = load(inp_all3(cc).output_file);
        com_samp = l.com_samp;

        Stats_regime3(com_samp, abd_bin_edges, dist_bin_edges, 0.25, 5, inp_all3(cc).L, [inp_all3(cc).output_file '_analysis_results.mat']); %run analysis for this regime and save result separately
    end
end


%% Compute stats ED and R20 
load('inp_CNDD_HNDD3.mat')

ref = cell(1,2);
for dd = 1:length(dist_vals)
    ref{dd} = load(['ref_HNDD_D_' sprintf('%0.4g',dist_vals(dd)) '_analysis_results.mat']);
end

ED_HCNDD3 = nan(size(inp_all3));
R20_HCNDD3 = nan(size(inp_all3));

for ch=1:length(magnitude_vals)
    for dd = 1:length(dist_vals)
        % Compute statistics compared to DL:
        reg_now = load([inp_all3(ch,dd).output_file '_analysis_results.mat']);
        ED_HCNDD3(ch,dd) = nansum((reg_now.mean_log_dist - ref{dd}.mean_log_dist).*reg_now.samps_per_bin)/sum(reg_now.samps_per_bin); %weighted mean
        R20_HCNDD3(ch,dd) = nansum((reg_now.log_N20 - ref{dd}.log_N20).*reg_now.samps_per_bin)/sum(reg_now.samps_per_bin); %weighted mean
    end
end

save('ED_HCNDD3','R20_HCNDD3', 'magnitude_vals','dist_vals') % THIS FILE IS PROVIDED

%% Supplementary figure for CNDD = HNDD with 20 m. DD:

figure()
s1 = subplot(2,1,1);
plot(magnitude_vals,R20_HCNDD3,'-x','LineWidth',1.5,'MarkerSize',8)
ylabel('EA(20)')
xlabel('DD magnitude')
set(s1,'FontSize',16,'LineWidth',1.5);
text(0.2,-0.05,'(A)','FontSize',16,'FontWeight','bold')
ylim([-.4 0])

s1 = subplot(2,1,2);
plot(magnitude_vals,ED_HCNDD3,'-x','LineWidth',1.5,'MarkerSize',8)
ylabel('ED')
xlabel('DD magnitude')
text(0.2,0.095,'(B)','FontSize',16,'FontWeight','bold')
set(s1,'FontSize',16,'LineWidth',1.5);
```
