## Supplementary material for "Pervasive species-specific repulsion among adult tropical trees": Code appendix: complete_sims.m

```
% compose the pieces of the simulations that haven't ended

%% Find all directories that have to be analyzed:

abd_bin_edges = [1 5:2:51 55:5:100 110:10:200 250:50:600 700:100:1000 1250 1500 5500];
dist_bin_edges = 0:10:200;

%assumption: if it's a folder it hasn't been deleted because the simulation hasn't finished running
D = dir;
D = D(3:end);

incomplete = find([D.isdir]); %indexes of incomplete simulations

parfor fff = 1:length(incomplete)
    ff = incomplete(fff);
    
    reg_now = D(ff).name;
    inp = load([reg_now '/inp.mat'],'inp'); %load the input file from within the folder
    inp = inp.inp;
    D2 = dir([reg_now '/']);
    next_samp = length(D2) - 2;
    
    [~, ~, ~] = sim3_continue(inp,next_samp);
    
end
```
