## Supplementary material for "Pervasive species-specific repulsion among adult tropical trees": Code appendix: Create_tab1_S1_2.m

```
%%%%%%%%%%%%%%%%%%%%%%%%%%%%%%%%%%%%%%%%%%%%%%%%%%%%%%%%%%%%%%%%%%%%%%%%%%%%%%%%%
% Summarize species table
%%%%%%%%%%%%%%%%%%%%%%%%%%%%%%%%%%%%%%%%%%%%%%%%%%%%%%%%%%%%%%%%%%%%%%%%%%%%%%%%%
%
% Author: Michael Kalyuzhny
% Date created: 20/01/2020
% Date last modified: 04/11/2020
%
%%%%%%%%%%%%%%%%%%%%%%%%%%%%%%%%%%%%%%%%%%%%%%%%%%%%%%%%%%%%%%%%%%%%%%%%%%%%%%%%%%%%%%%%%%%%

load('species_data3.mat','sp_dat')

%% Create table1A
stats_to_analyze = {'R20l_08', 'R20l_01', 'R20l_08_lag', 'R20l_08_LDD', 'R20l_08_dem', 'R20l_08_fixed'};
sig_to_analyze = {'R20l_08_pval', 'R20l_01_pval','R20l_08_lag_pval', 'R20l_08_LDD_pval', 'R20l_08_dem_pval', 'R20l_08_fixed_pval'};
    
stats_total = length(stats_to_analyze);

overdispersed = strings(stats_total, 1);
clumped = strings(stats_total, 1);
summary = strings(stats_total, 1);

for ii = 1:stats_total
    vals = (sp_dat{:,stats_to_analyze{ii}}); %the values of the statistic
    vals = vals(~isnan(vals)); %remove missing values
    pvals = (sp_dat{:,sig_to_analyze{ii}});
    pvals = pvals(~isnan(pvals));
    
    summary(ii) = [num2str(mean(vals),2) ' ' char(177) ' ' num2str(std(vals),2) ' [' num2str(min(vals),2) ', ' num2str(max(vals), 2) ']'];
   
    over = sum(vals <0 ); % overdispersed
    over_sig = sum((vals < 0) & (pvals < .05)); %overdispersed and significant
    cl = sum(vals >0 ); %clumped
    cl_sig = sum((vals > 0) & (pvals < .05)); %clumped and sig.
    overdispersed(ii) = [num2str(over) ' (' num2str(over_sig) ')'];
    clumped(ii) = [num2str(cl) ' (' num2str(cl_sig) ')'];
end

col_names = {'Values', 'Overdispersed', 'Aggregated'};
row_names = {'Standard', 'H2001', 'Lag', 'LDD', 'Recruits', 'Fixed'};

table1A = table(summary, overdispersed, clumped,'VariableNames',col_names, 'RowNames', row_names)
writetable(table1A,'table1.xls')

%% Create table1B
stats_to_analyze = {'ED_mean_08', 'ED_mean_01', 'ED_mean_08_lag', 'ED_med_08_LDD', 'ED_med_08_dem', 'ED_med_08_fixed'};
sig_to_analyze = {'ED_mean_08_pval', 'ED_mean_01_pval','ED_mean_08_lag_pval', 'ED_med_08_LDD_pval', 'ED_med_08_LDD_pval', 'ED_med_08_fixed_pval'};
    
stats_total = length(stats_to_analyze);

overdispersed = strings(stats_total, 1);
clumped = strings(stats_total, 1);
summary = strings(stats_total, 1);

for ii = 1:stats_total
    vals = (sp_dat{:,stats_to_analyze{ii}}); %the values of the statistic
    vals = vals(~isnan(vals)); %remove missing values
    pvals = (sp_dat{:,sig_to_analyze{ii}});
    pvals = pvals(~isnan(pvals));
    
    summary(ii) = [num2str(mean(vals),2) ' ' char(177) ' ' num2str(std(vals),2) ' [' num2str(min(vals),2) ', ' num2str(max(vals), 2) ']'];
   
    over = sum(vals > 0); % overdispersed
    over_sig = sum((vals > 0) & (pvals < .05)); %overdispersed and significant
    cl = sum(vals < 0); %clumped
    cl_sig = sum((vals < 0) & (pvals < .05)); %clumped and sig.
    overdispersed(ii) = [num2str(over) ' (' num2str(over_sig) ')'];
    clumped(ii) = [num2str(cl) ' (' num2str(cl_sig) ')'];
end

col_names = {'Values', 'Overdispersed', 'Aggregated'};
row_names = {'Standard', 'H2001', 'Lag', 'LDD', 'Recruits', 'Fixed'};

table1B = table(summary, overdispersed, clumped,'VariableNames',col_names, 'RowNames', row_names)
writetable(table1B,'table1B.xls')

%% Create table in supplement
stats_to_analyze = {'R75_125l_08', 'R75_125l_01', 'R75_125l_08_lag', 'R75_125l_08_LDD', 'R75_125l_08_dem', 'R75_125l_08_fixed'};
sig_to_analyze = {'R75_125l_08_pval', 'R75_125l_01_pval','R75_125l_08_lag_pval', 'R75_125l_08_LDD_pval', 'R75_125l_08_dem_pval', 'R75_125l_08_fixed_pval'};
    
stats_total = length(stats_to_analyze);

overdispersed = strings(stats_total, 1);
clumped = strings(stats_total, 1);
summary = strings(stats_total, 1);

for ii = 1:stats_total
    vals = (sp_dat{:,stats_to_analyze{ii}}); %the values of the statistic
    not_nan = ~isnan(vals);
    vals = vals(not_nan); %remove missing values
    pvals = (sp_dat{:,sig_to_analyze{ii}});
    pvals = pvals(not_nan);
    
    summary(ii) = [num2str(mean(vals),2) ' ' char(177) ' ' num2str(std(vals),2) ' [' num2str(min(vals),2) ', ' num2str(max(vals), 2) ']'];
   
    over = sum(vals < 0); % overdispersed
    over_sig = sum((vals < 0) & (pvals < .05)); %overdispersed and significant
    cl = sum(vals > 0); %clumped
    cl_sig = sum((vals > 0) & (pvals < .05)); %clumped and sig.
    overdispersed(ii) = [num2str(over) ' (' num2str(over_sig) ')'];
    clumped(ii) = [num2str(cl) ' (' num2str(cl_sig) ')'];
end

col_names = {'Values', 'Overdispersed', 'Aggregated'};
row_names = {'Standard', 'H2001', 'Lag', 'LDD', 'Recruits', 'Fixed'};

table_ED1 = table(summary, overdispersed, clumped,'VariableNames',col_names, 'RowNames', row_names)
writetable(table_ED1,'table_ED1.xls')
```
