## Supplementary material for "Pervasive species-specific repulsion among adult tropical trees": Code appendix: Full_analysis.m

```
%%%%%%%%%%%%%%%%%%%%%%%%%%%%%%%%%%%%%%%%%%%%%%%%%%%%%%%%%%%%%%%
% BCI spatial analysis - full analysis pipeline
%%%%%%%%%%%%%%%%%%%%%%%%%%%%%%%%%%%%%%%%%%%%%%%%%%%%%%%%%%%%%%%
%
% Author: Michael Kalyuzhny
% First created: 01/01/2021
% Last modified: 12/01/2023
%
% This file follows the full analysis, step by step.
% We mostly modify and analyze a data table, 'sp_dat'. First, it is created from various sources. Next, simulations of the null models are run and analyzed to obtain spatial statistics, with (most) statistics added to the table. Finally, the table is analyzed.

%% Creating the data table

generate_sp_dat_table1; %Create the basic data table

% This creates 'species_data1.mat' file, with the input to the null model
% analysis
%% Run the null model for every species, using the data from the data table:

run_sims_null1_2008;
run_sims_null1_2001;
run_sims_null2_lag_2008;
run_sims_null3_LDD_2008;
run_sims_null4_initial_2008;
run_sims_null6_2008; 

% Each of these scripts moves to the Results folder to run the analysis,
% where the simulation results are saved

%% Create a table of the resulting statistics:
Analyze_all_stats3_log;

% This uses the raw simulation results and adds the statistics to the
% 'sp_dat' table (and also saves separately), creating 'species_data2.mat'
% file, with the spatial stats

%% Theoretical analysis of CNDD and HNDD

% This section WILL take a long time. 
% If the simulations have to be terminated, 'compose pieces' can be used to compose the results from sinlge files.
% alternatively, 'complete_sims' can finish the terminated simulations from the last sample taken

cd('CNDD_HNDD')
Analysis_CNDD_HNDD_Delta7;
Analysis_CNDD_HNDD_Delta20;
cd('..')
%% Final analysis and presentations:

generate_fig1_figS1_ver2;
Create_tab1_S1_2;
Generate_Fig2_and_SI2;
generate_fig_EA_dist1; 

%% Clustered dispersal analysis:

cd('Clustered_disp')
analyze_sim_clustered_dispersal2;
cd('..')
```
