## Supplementary material for "Pervasive species-specific repulsion among adult tropical trees": Code appendix: generate_fig1_figS1_ver2.m

```
%%%%%%%%%%%%%%%%%%%%%%%%%%%%%%%%%%%%%%%%%%%%%%%
% Generate Figures 1 and S1-S2
%%%%%%%%%%%%%%%%%%%%%%%%%%%%%%%%%%%%%%%%%%%%%%%
%
% Author: Michael Kalyuzhny
% First written: 13/01/2020
% Last modified: 21/03/2020
%
%% Load data

load('species_data1.mat'); %species summary table
load('Results/spatial_distributions.mat', 'point_2015','point_1985')
%% Pick species:

%sp_code = 'oenoma'; %common, rather generalist but short dispersal VV
%sp_code = 'dendar';% VVVV fully generalist, intermediat, dispersal: 27
sp_code = 'beilpe';% intermediate, short dispersal, somewhat specialist

sp_ind = find(strcmp(sp_dat{:,'SpeciesCode'},sp_code)); %index in sp_dat
sp_abd = sp_dat{sp_ind, 'abd_adults_2015'}; %abudnance of this species

samp_num = 3; %which sample to plot (in each null)?

%% Load and analyze null1 (2008 distances):

load(['Results/null1_2008_' sp_code '.mat'],'com_samp'); %load samples

%cut edges:
for s = 1:size(com_samp,3)
   com_samp(((com_samp(:,1,s) < 100) | (com_samp(:,1,s) > 1100) | (com_samp(:,2,s) < 350) | (com_samp(:,2,s) > 850)), :, s) = nan; %remove data on tree with at least one coordinate out of bounds
   com_samp(:,1:2,s) = com_samp(:,1:2,s) - [100 350];
end
cs = sum_pop(com_samp);

[row,col] = find(cs == sp_abd);

%% Set sample:
null1_col = col(samp_num); 
null1_row = row(samp_num);
inds_in_null1 = com_samp(:, 3, null1_row) == null1_col; %the indexes of the reference in the data file of samples

%% generate Fig. 1:

figure()

s1 = subplot(3,1,1);
hold on
set(s1,'FontSize',16,'FontWeight','bold','LineWidth',1.5);
scatter(point_2015{sp_ind}(:,1), point_2015{sp_ind}(:,2), 24, ...
    'MarkerEdgeColor',[0 0.2 1], 'MarkerFaceColor',[0.4 .85 0])
xlim([0 1000])
ylim([0 500])
xticks([0 250 500 750 1000])
yticks([0 250 500 750 1000])
text(10,470,'a','FontSize',16,'FontWeight','bold')
title('Observed')


s2 = subplot(3,1,2);
rand_xy = rand(sp_abd,2)*[1000 0; 0 500];
hold on
set(s2,'FontSize',16,'FontWeight','bold','LineWidth',1.5);
scatter(rand_xy(:,1), rand_xy(:,2), 24, ...
    'MarkerEdgeColor','cyan', 'MarkerFaceColor',[0.4 .85 0])
xlim([0 1000])
ylim([0 500])
xticks([0 250 500 750 1000])
yticks([0 250 500 750 1000])
ylabel('Y (meters)')
title('CSR null')
text(10,470,'b','FontSize',16,'FontWeight','bold')


s3 = subplot(3,1,3);
hold on
set(s3,'FontSize',16,'FontWeight','bold','LineWidth',1.5);
scatter(com_samp(inds_in_null1, 1, null1_row), com_samp(inds_in_null1, 2, null1_row), 24, ...
    'MarkerEdgeColor','black', 'MarkerFaceColor',[0.4 .85 0])
xlim([0 1000])
ylim([0 500])
xticks([0 250 500 750 1000])
yticks([0 250 500 750 1000])
xlabel('X (meters)')
title('Dispersal Limitation null')
text(10,470,'c','FontSize',16,'FontWeight','bold')

%% Now proceed to Fig S1 - begin as before:

figure()

s1 = subplot(4,4,[1 2]);
hold on
set(s1,'FontSize',16,'FontWeight','bold','LineWidth',1.5);
scatter(point_2015{sp_ind}(:,1), point_2015{sp_ind}(:,2), 24, ...
    'MarkerEdgeColor',[0 0.2 1], 'MarkerFaceColor',[0.4 .85 0])
xlim([0 1000])
ylim([0 500])
xticks([0 250 500 750 1000])
yticks([0 250 500 750 1000])
text(10,450,'a','FontSize',16,'FontWeight','bold')
title('Observed')


% s2 = subplot(4,4,[5 6]);
% rand_xy = rand(sp_abd,2)*[1000 0; 0 500];
% hold on
% set(s2,'FontSize',16,'FontWeight','bold','LineWidth',1.5);
% scatter(rand_xy(:,1), rand_xy(:,2), 24, ...
%     'MarkerEdgeColor','cyan', 'MarkerFaceColor',[0.4 .85 0])
% xlim([0 1000])
% ylim([0 470])
% xticks([0 250 500 750 1000])
% yticks([0 250 500 750 1000])
% ylabel('Y (meters)')
% title('CSR null')
% text(10,450,'b','FontSize',16,'FontWeight','bold')


s3 = subplot(4,4, [5 6]);
hold on
set(s3,'FontSize',16,'FontWeight','bold','LineWidth',1.5);
scatter(com_samp(inds_in_null1, 1, null1_row), com_samp(inds_in_null1, 2, null1_row), 24, ...
    'MarkerEdgeColor','black', 'MarkerFaceColor',[0.4 .85 0])
xlim([0 1000])
ylim([0 500])
xticks([0 250 500 750 1000])
yticks([0 250 500 750 1000])
title('Standard DL null')
text(10,470,'b','FontSize',16,'FontWeight','bold')


%% Load and analyze null1 (2001 distances):

load(['Results/null1_2001_' sp_code '.mat'],'com_samp'); %load samples

%cut edges:
for s = 1:size(com_samp,3)
   com_samp(((com_samp(:,1,s) < 100) | (com_samp(:,1,s) > 1100) | (com_samp(:,2,s) < 350) | (com_samp(:,2,s) > 850)), :, s) = nan; %remove data on tree with at least one coordinate out of bounds
   com_samp(:,1:2,s) = com_samp(:,1:2,s) - [100 350];
end
cs = sum_pop(com_samp);

[row,col] = find(cs == sp_abd);

%% Set sample (2001):
null1_col = col(samp_num); 
null1_row = row(samp_num);
inds_in_null1 = com_samp(:, 3, null1_row) == null1_col; %the indexes of the reference in the data file of samples

%% Add to plot:

s4 = subplot(4,4, [9 10]);
hold on
set(s4,'FontSize',16,'FontWeight','bold','LineWidth',1.5);
scatter(com_samp(inds_in_null1, 1, null1_row), com_samp(inds_in_null1, 2, null1_row), 24, ...
    'MarkerEdgeColor','black', 'MarkerFaceColor',[0.4 .85 0])
xlim([0 1000])
ylim([0 500])
xticks([0 250 500 750 1000])
yticks([0 250 500 750 1000])
title('H2001 DL null')
text(10,470,'c','FontSize',16,'FontWeight','bold')
ylabel('Y (meters)')

%% Load and analyze null2 (lag):

load(['Results/null2_2008_' sp_code '.mat'],'com_samp'); %load samples

%cut edges:
for s = 1:size(com_samp,3)
   com_samp(((com_samp(:,1,s) < 100) | (com_samp(:,1,s) > 1100) | (com_samp(:,2,s) < 350) | (com_samp(:,2,s) > 850)), :, s) = nan; %remove data on tree with at least one coordinate out of bounds
   com_samp(:,1:2,s) = com_samp(:,1:2,s) - [100 350];
end
cs = sum_pop(com_samp);

[row,col] = find(cs == sp_abd);

%% Set sample (null2):
null1_col = col(samp_num); 
null1_row = row(samp_num);
inds_in_null1 = com_samp(:, 3, null1_row) == null1_col; %the indexes of the reference in the data file of samples

%% Add to plot:

s5 = subplot(4,4, [11 12]);
hold on
set(s5,'FontSize',16,'FontWeight','bold','LineWidth',1.5);
scatter(com_samp(inds_in_null1, 1, null1_row), com_samp(inds_in_null1, 2, null1_row), 24, ...
    'MarkerEdgeColor','black', 'MarkerFaceColor',[0.4 .85 0])
xlim([0 1000])
ylim([0 500])
xticks([0 250 500 750 1000])
yticks([0 250 500 750 1000])
title('Lag DL null')
text(10,470,'g','FontSize',16,'FontWeight','bold')


%% Load and analyze the Fixed null:

load(['Results/null6_Fiexd_Trees_2008_' sp_code '.mat'],'com_samp'); %load samples

cs = sum_pop(com_samp);

[row,col] = find(cs == sp_abd);

%% Set sample (Fixed null):
null1_col = col(samp_num); 
null1_row = row(samp_num);
inds_in_null1 = com_samp(:, 3, null1_row) == null1_col; %the indexes of the reference in the data file of samples

%% Add to plot:

s6 = subplot(4,4, [13 14]);
hold on
set(s6,'FontSize',16,'FontWeight','bold','LineWidth',1.5);
scatter(com_samp(inds_in_null1, 1, null1_row), com_samp(inds_in_null1, 2, null1_row), 24, ...
    'MarkerEdgeColor','black', 'MarkerFaceColor',[0.4 .85 0])
xlim([0 1000])
ylim([0 500])
xticks([0 250 500 750 1000])
yticks([0 250 500 750 1000])
title('Fixed DL null')
text(10,470,'d','FontSize',16,'FontWeight','bold')

%% Load and analyze null3 (LDD):

load(['Results/null3_2008_' sp_code '.mat'],'com_samp'); %load samples

%cut edges:
for s = 1:size(com_samp,3)
   com_samp(((com_samp(:,1,s) < 100) | (com_samp(:,1,s) > 1100) | (com_samp(:,2,s) < 350) | (com_samp(:,2,s) > 850)), :, s) = nan; %remove data on tree with at least one coordinate out of bounds
   com_samp(:,1:2,s) = com_samp(:,1:2,s) - [100 350];
end
cs = sum_pop(com_samp);

[row,col] = find(cs == sp_abd);

%% Set sample (null2):
null1_col = col(samp_num); 
null1_row = row(samp_num);
inds_in_null1 = com_samp(:, 3, null1_row) == null1_col; %the indexes of the reference in the data file of samples

%% Add to plot:

s6 = subplot(4,4, [15 16]);
hold on
set(s6,'FontSize',16,'FontWeight','bold','LineWidth',1.5);
scatter(com_samp(inds_in_null1, 1, null1_row), com_samp(inds_in_null1, 2, null1_row), 24, ...
    'MarkerEdgeColor','black', 'MarkerFaceColor',[0.4 .85 0])
xlim([0 1000])
ylim([0 500])
xticks([0 250 500 750 1000])
yticks([0 250 500 750 1000])
title('LDD DL null')
text(10,470,'h','FontSize',16,'FontWeight','bold')
xlabel('X (meters)')

%% Load and analyze null4 (initial state):

load(['Results/null4_2008_' sp_code '.mat']); %load samples

%% Add to plot:

unchanged = point_1985{sp_ind}(ismember(point_1985{sp_ind},point_2015{sp_ind},'rows'), :);
recruits_obs = point_2015{sp_ind}(~ismember(point_2015{sp_ind},point_1985{sp_ind},'rows'), :);
mortality_obs = point_1985{sp_ind}(~ismember(point_1985{sp_ind},point_2015{sp_ind},'rows'), :);
recruits_null = points_final{samp_num}(is_new{samp_num},:); %trees that were there in the first survey and survived
mortality_null =  point_1985{sp_ind}(~ismember(point_1985{sp_ind},points_final{samp_num},'rows'), :);
unchanged_null = point_1985{sp_ind}(ismember(point_1985{sp_ind},points_final{samp_num},'rows'), :);

s7 = subplot(4,4, [3 4]);
hold on
set(s7,'FontSize',16,'FontWeight','bold','LineWidth',1.5);

scatter(unchanged(:,1), unchanged(:,2), 24, ...
    'MarkerEdgeColor',[0 0.2 1], 'MarkerFaceColor',[0.4 .85 0]) %unchanged

scatter(mortality_obs(:,1), mortality_obs(:,2), 24, ...
    'MarkerEdgeColor',[0 0.2 1], 'MarkerFaceColor',[1 1 1]) %mortality observed

scatter(recruits_obs(:,1), recruits_obs(:,2), 24, '^', ...
    'MarkerEdgeColor',[0 0.2 1], 'MarkerFaceColor',[0 0.3 1]) %recruitment observed
xlim([0 1000])
ylim([0 500])
xticks([0 250 500 750 1000])
yticks([0 250 500 750 1000])
title('Observed demography')
text(10,470,'e','FontSize',16,'FontWeight','bold')

s8 = subplot(4,4, [7 8]);
hold on
set(s8,'FontSize',16,'FontWeight','bold','LineWidth',1.5);

scatter(unchanged_null(:,1), unchanged_null(:,2), 24, ...
    'MarkerEdgeColor',[0 0.2 1], 'MarkerFaceColor',[0.4 .85 0]) %unchanged

scatter(mortality_null(:,1), mortality_null(:,2), 24, ...
    'MarkerEdgeColor',[0 0.2 1], 'MarkerFaceColor',[1 1 1]) %mortality observed

scatter(recruits_null(:,1), recruits_null(:,2), 24, '^', ...
    'MarkerEdgeColor',[0 0.2 1], 'MarkerFaceColor',[0 0.3 1]) %recruitment observed
xlim([0 1000])
ylim([0 500])
xticks([0 250 500 750 1000])
yticks([0 250 500 750 1000])
title('Recruits DL null demography')
text(10,470,'f','FontSize',16,'FontWeight','bold')

%% Zoom on a part of the landscape in null and real data:

%% (re)Load data:

load(['Results/null1_2008_' sp_code '.mat'],'com_samp'); %load samples

%load(['Results/null6_Fiexd_Trees_2008_' sp_code '.mat'],'com_samp'); %load samples

%cut edges:
for s = 1:size(com_samp,3)
   com_samp(((com_samp(:,1,s) < 100) | (com_samp(:,1,s) > 1100) | (com_samp(:,2,s) < 350) | (com_samp(:,2,s) > 850)), :, s) = nan; %remove data on tree with at least one coordinate out of bounds
   com_samp(:,1:2,s) = com_samp(:,1:2,s) - [100 350];
end
cs = sum_pop(com_samp);

[row,col] = find(cs == sp_abd);

% Set sample:
null1_col = col(samp_num); 
null1_row = row(samp_num);
inds_in_null1 = com_samp(:, 3, null1_row) == null1_col; %the indexes of the reference in the data file of samples

load('bci_trees_data.mat','bci')

%% Plot it:

xlim_zoom = [200 400];
ylim_zoom = [200 400];

figure()
s1 = subplot(1,2,1);
hold on
set(s1,'FontSize',16,'FontWeight','bold','LineWidth',1.5);
scatter(com_samp(:, 1, null1_row), com_samp(:, 2, null1_row), 18, ...
    'MarkerEdgeColor','black', 'MarkerFaceColor',[1 1 1])
hold on
scatter(com_samp(inds_in_null1, 1, null1_row), com_samp(inds_in_null1, 2, null1_row), 18, ...
    'MarkerEdgeColor','black', 'MarkerFaceColor',[1 0.2 0])
xlim(xlim_zoom)
ylim(ylim_zoom)
xlabel('X (m.)')
ylabel('Y (m.)')
xticks([xlim_zoom(1) : 50 : xlim_zoom(2)])
yticks([ylim_zoom(1) : 50 : ylim_zoom(2)])
text(xlim_zoom(1) + 5, ylim_zoom(2) - 10,'a','FontSize',16,'FontWeight','bold')
title('Standard DL null')


%% Empirical pattern:

% Find all living trees above this size threshold:
take =   bci{7}{:,'dbh'} >= (10*sp_dat{sp_ind,'Rep_thresh_use'}) & ... %not too small
         strcmp(bci{7}{:,'status'}, 'A') & ... %alive
         ~isnan(bci{7}{:,'gx'}) & ~isnan(bci{7}{:,'gy'}); 

s1 = subplot(1,2,2);
hold on
set(s1,'FontSize',16,'FontWeight','bold','LineWidth',1.5);
scatter(bci{7}{take,'gx'}, bci{7}{take,'gy'}, 18, ...
    'MarkerEdgeColor','black', 'MarkerFaceColor',[1 1 1])
xlim(xlim_zoom)
ylim(ylim_zoom)
xlabel('X (m.)')
xticks([xlim_zoom(1) : 50 : xlim_zoom(2)])
yticks([ylim_zoom(1) : 50 : ylim_zoom(2)])
text(xlim_zoom(1) + 5, ylim_zoom(2) - 10,'b','FontSize',16,'FontWeight','bold')
title('Observed')

%% Export:
f1 = gco;
set(f1,'PaperSize',[9 12]); %set the paper size to what you want
print(f1,'ED Figure 1','-dpdf')
```
