## Supplementary material for "Pervasive species-specific repulsion among adult tropical trees": Code appendix: Generate_Fig2_and_SI_2.m

```
%%%%%%%%%%%%%%%%%%%%%%%%%%%%%%%%%%%%%%%%%%%%%%%%%%%%%%%%%%%%%%%%%%%%%%%%%%%%%%%%%
% Generate the figure of Relative Neighborhood Density and the extra figure(s) of multiple species
%%%%%%%%%%%%%%%%%%%%%%%%%%%%%%%%%%%%%%%%%%%%%%%%%%%%%%%%%%%%%%%%%%%%%%%%%%%%%%%%%
%
% Author: Michael Kalyuzhny
% Date created: 22/01/2021
% Date last modified: 11/02/2022
%
%%%%%%%%%%%%%%%%%%%%%%%%%%%%%%%%%%%%%%%%%%%%%%%%%%%%%%%%%%%%%%%%%%%%%%%%%%%%%%%%%%%%%%%%%%%%

%% Fig. 2: setup

% Load required data and set some settings:

load('species_data3.mat','sp_dat')
load('Results/spatial_distributions.mat')
load('Results/Null1_2008_stats.mat') 
bin_centers = 5:10:195;

species_inds = [45 57 74 51]; %the second and fourth species will be presented with their distribution maps
inds_in_2008 = [22 30 37 26]; % indexes for 2008 subset of data, to be used for the R(r) plots where
%extra_labels = ['a' 'b' 'd' 'e']; 
species_names = {'O. mapora', 'R. armata', 'T. tuberculata', 'P. reticulata'};
%R_locations = [1 2 5 6]; %the locations of the subplots of R(r)

species_tot = length(inds_in_2008);

%% Obtain data from nulls:

samp_null_take = 1;

points_nulls = cell(1,species_tot);

for ss = 1:species_tot
    
    sp_abd = length(point_2015{species_inds(ss)}); %abundance

    load(['Results/null1_2008_' sp_dat.SpeciesCode{species_inds(ss)} '.mat'],'com_samp'); %load samples

    %cut edges:
    for s = 1:size(com_samp,3)
       com_samp(((com_samp(:,1,s) < 100) | (com_samp(:,1,s) > 1100) | (com_samp(:,2,s) < 350) | (com_samp(:,2,s) > 850)), :, s) = nan; %remove data on tree with at least one coordinate out of bounds
       com_samp(:,1:2,s) = com_samp(:,1:2,s) - [100 350];
    end
    
    %find appropriate null sample:
    cs = sum_pop(com_samp);
    [row,col] = find(cs == sp_abd);
    null1_col = col(samp_null_take); 
    null1_row = row(samp_null_take);
    inds_in_null1 = com_samp(:, 3, null1_row) == null1_col; %the indexes of the reference in the data file of samples
    
    points_nulls{ss} = [com_samp(inds_in_null1, 1, null1_row), com_samp(inds_in_null1, 2, null1_row)];

end
%%

f = figure();

% LOOP OVER ALL SPECIES

for ss = 1:species_tot
    
    % Plot observed distribution:
    s1 = subplot(species_tot,5,[((ss-1)*5 + 1) ((ss-1)*5 + 2)]);
    hold on
    set(s1,'FontSize',16,'FontWeight','bold','LineWidth',1.5,'XGrid','on','YGrid','on');
    scatter(point_2015{species_inds(ss)}(:,1), point_2015{species_inds(ss)}(:,2), 12, ...
        'MarkerEdgeColor',[0 0.2 1], 'MarkerFaceColor',[0.4 .85 0])
    xlim([0 1000])
    ylim([0 500])
    
    xticks(0:100:1000)
    yticks(0:100:500)
    xticklabels(repmat("",1,11));
    yticklabels(repmat("",1,6));
    text(-40,250,['\it' species_names{ss}],'FontSize',20,'FontWeight','bold','Rotation',90)
    
    if ss == 1
        title('Observed','FontSize',20)
        ylabel('Y (m.)')
    elseif ss == 4
        xlabel('X (m.)')
    end
    
    
    % Plot null:
    s1 = subplot(species_tot,5,[((ss-1)*5 + 3) ((ss-1)*5 + 4)]);
    hold on
    set(s1,'FontSize',16,'FontWeight','bold','LineWidth',1.5,'XGrid','on','YGrid','on');
    scatter(points_nulls{ss}(:,1), points_nulls{ss}(:,2), 12, ...
        'MarkerEdgeColor','black', 'MarkerFaceColor',[0.4 .85 0])
    xlim([0 1000])
    ylim([0 500])
    xticks(0:100:1000)
    yticks(0:100:500)
    xticklabels(repmat("",1,11));
    yticklabels(repmat("",1,6));
    
    if ss == 1
        title('Dispersal Limitation null','FontSize',20)
        ylabel('Y (m.)')
    elseif ss == 4
        xlabel('X (m.)')
    end
    
    % Plot EA(r):
    s1 = subplot(species_tot,5,(ss-1)*5 + 5);
    hold on
    set(s1,'FontSize',16,'FontWeight','bold','LineWidth',1.5);

    x_fill=[bin_centers fliplr(bin_centers)];
    y_fill=[R_sim_envelope(2, :, inds_in_2008(ss)) fliplr(R_sim_envelope(1, :, inds_in_2008(ss)))];
    fill(x_fill, y_fill , 1,'facecolor',[0.95 0.7 0],'edgecolor','none','facealpha', 0.5);
    plot(bin_centers, R(inds_in_2008(ss),:),'-o','MarkerEdgeColor',[0 0.45 0.74], 'MarkerFaceColor', [.3 .75 .93], 'Color', [0 0.45 0.74],'LineWidth',1.5)
    line([0 200],[0 0], 'LineStyle','--', 'Color','black','LineWidth',1.5)

    xlim([0 200])
    ybounds = [min(min(R(inds_in_2008(ss),:)) - 0.1, -0.2), max(.2, max(R(inds_in_2008(ss),:)) + .1)];
    ylim(ybounds) %upper limit: 1.2 or the maximal R + 0.1, whichever is greater. Lower: 
    xticks([0 100 200])
    text(5,0.1, 'Aggregated','FontSize',12)
    text(5,-0.1, 'Overdispersed','FontSize',12)
    %title(['\it' species_names{ss}])
    if ss == 4
        xlabel('Distance (m.)')
    elseif ss == 1
        ylabel('Excess Abundance (EA(r))')
        title('EA(r)','FontSize',20)
    end
    
end

%%
set(f,'PaperSize',[16 16]); %set the paper size to what you want
print('-painters',f,'Fig2','-dpdf')

%%%%%%%%%%%%%%%%%%%%%%%%%%%%%%%%%%%%%%%%%%%%%%%%%%%%%%%%%%%%%%%%%%%%%%%%%%%
%% REPEAT WITH EXTRAORDINARY SPECIES
%%%%%%%%%%%%%%%%%%%%%%%%%%%%%%%%%%%%%%%%%%%%%%%%%%%%%%%%%%%%%%%%%%%%%%%%%%%
%%
species_inds = [13 82 14 37]; %the second and fourth species will be presented with their distribution maps
inds_in_2008 = [6 41 7 20]; % indexes for 2008 subset of data, to be used for the R(r) plots where
species_names = {'C. curvigemmia', 'Z. ekmanii', 'C. billbergianus', 'J. copaia'};

species_tot = length(inds_in_2008);

%% Obtain data from nulls:

samp_null_take = 2;

points_nulls = cell(1,species_tot);

for ss = 1:species_tot
    
    sp_abd = length(point_2015{species_inds(ss)}); %abundance

    load(['Results/null1_2008_' sp_dat.SpeciesCode{species_inds(ss)} '.mat'],'com_samp'); %load samples

    %cut edges:
    for s = 1:size(com_samp,3)
       com_samp(((com_samp(:,1,s) < 100) | (com_samp(:,1,s) > 1100) | (com_samp(:,2,s) < 350) | (com_samp(:,2,s) > 850)), :, s) = nan; %remove data on tree with at least one coordinate out of bounds
       com_samp(:,1:2,s) = com_samp(:,1:2,s) - [100 350];
    end
    
    %find appropriate null sample:
    cs = sum_pop(com_samp);
    [row,col] = find(cs == sp_abd);
    null1_col = col(samp_null_take); 
    null1_row = row(samp_null_take);
    inds_in_null1 = com_samp(:, 3, null1_row) == null1_col; %the indexes of the reference in the data file of samples
    
    points_nulls{ss} = [com_samp(inds_in_null1, 1, null1_row), com_samp(inds_in_null1, 2, null1_row)];

end
%%

f2 = figure();

% LOOP OVER ALL SPECIES

for ss = 1:species_tot
    
    % Plot observed distribution:
    s1 = subplot(species_tot,5,[((ss-1)*5 + 1) ((ss-1)*5 + 2)]);
    hold on
    set(s1,'FontSize',16,'FontWeight','bold','LineWidth',1.5,'XGrid','on','YGrid','on');
    scatter(point_2015{species_inds(ss)}(:,1), point_2015{species_inds(ss)}(:,2), 12, ...
        'MarkerEdgeColor',[0 0.2 1], 'MarkerFaceColor',[0.4 .85 0])
    xlim([0 1000])
    ylim([0 500])
    
    xticks(0:100:1000)
    yticks(0:100:500)
    xticklabels(repmat("",1,11));
    yticklabels(repmat("",1,6));
    text(-40,250,['\it' species_names{ss}],'FontSize',20,'FontWeight','bold','Rotation',90)
    
    if ss == 1
        title('Observed','FontSize',20)
        ylabel('Y (m.)')
    elseif ss == 4
        xlabel('X (m.)')
    end
    
    
    % Plot null:
    s1 = subplot(species_tot,5,[((ss-1)*5 + 3) ((ss-1)*5 + 4)]);
    hold on
    set(s1,'FontSize',16,'FontWeight','bold','LineWidth',1.5,'XGrid','on','YGrid','on');
    scatter(points_nulls{ss}(:,1), points_nulls{ss}(:,2), 12, ...
        'MarkerEdgeColor','black', 'MarkerFaceColor',[0.4 .85 0])
    xlim([0 1000])
    ylim([0 500])
    xticks(0:100:1000)
    yticks(0:100:500)
    xticklabels(repmat("",1,11));
    yticklabels(repmat("",1,6));
    
    if ss == 1
        title('Dispersal Limitation null','FontSize',20)
        ylabel('Y (m.)')
    elseif ss == 4
        xlabel('X (m.)')
    end
    
    % Plot EA(r):
    s1 = subplot(species_tot,5,(ss-1)*5 + 5);
    hold on
    set(s1,'FontSize',16,'FontWeight','bold','LineWidth',1.5);

    x_fill=[bin_centers fliplr(bin_centers)];
    y_fill=[R_sim_envelope(2, :, inds_in_2008(ss)) fliplr(R_sim_envelope(1, :, inds_in_2008(ss)))];
    fill(x_fill, y_fill , 1,'facecolor',[0.95 0.7 0],'edgecolor','none','facealpha', 0.5);
    plot(bin_centers, R(inds_in_2008(ss),:),'-o','MarkerEdgeColor',[0 0.45 0.74], 'MarkerFaceColor', [.3 .75 .93], 'Color', [0 0.45 0.74],'LineWidth',1.5)
    line([0 200],[0 0], 'LineStyle','--', 'Color','black','LineWidth',1.5)

    xlim([0 200])
    ybounds = [min(min(R(inds_in_2008(ss),:)) - 0.1, -0.2), max(.2, max(R(inds_in_2008(ss),:)) + .1)];
    ylim(ybounds) %upper limit: 1.2 or the maximal R + 0.1, whichever is greater. Lower: 
    xticks([0 100 200])
    text(5,0.1, 'Aggregated','FontSize',12)
    text(5,-0.1, 'Overdispersed','FontSize',12)
    %title(['\it' species_names{ss}])
    if ss == 4
        xlabel('Distance (m.)')
    elseif ss == 1
        ylabel('Excess Abundance (EA(r))')
        title('EA(r)','FontSize',20)
    end
    
end

%%
set(f2,'PaperSize',[16 16]); %set the paper size to what you want
print('-painters',f2,'SI_fig_EA','-dsvg')

%% Supplamentary figure with H2008 kernel:

inds_to_show = find((~isnan(sp_dat{:,'HML2008DistanceFit'})) & (sp_dat{:,'avg_log_abd'} >= log(30))); %which species should be analyzed?
x_subplots = 4; %numbers of subplots in each dimension
y_subplots = 8;
dists_2008 = sp_dat{~isnan(sp_dat{:,'HML2008DistanceFit'}),'HML2008DistanceFit'};

figure()

for ss = 1:length(inds_to_show)
    
    %Find index including only 2008 data:
    ind_2008_now = find((dists_2008==sp_dat{inds_to_show(ss),'HML2008DistanceFit'}) ...
        & (sp_dat{~isnan(sp_dat{:,'HML2008DistanceFit'}),'avg_log_abd'} == sp_dat{inds_to_show(ss),'avg_log_abd'}));
    
    
    s1 = subplot(y_subplots, x_subplots, ss);
    hold on
    set(s1,'FontSize',14,'FontWeight','bold','LineWidth',1.5);
    x_fill=[bin_centers fliplr(bin_centers)];
    y_fill=[R_sim_envelope(2, :, ind_2008_now) fliplr(R_sim_envelope(1, :, ind_2008_now))];
    fill(x_fill, y_fill , 1,'facecolor',[0.95 0.7 0],'edgecolor','none','facealpha', 0.5);
    plot(bin_centers, R(ind_2008_now,:),'-o','MarkerEdgeColor',[0 0.45 0.74], 'MarkerFaceColor', [.3 .75 .93], 'Color', [0 0.45 0.74],'LineWidth',1.5)
    line([0 200],[0 0], 'LineStyle','--', 'Color','black','LineWidth',1.5)

    xlim([0 200])
    ybounds = [min(min(R(ind_2008_now,:)) - 0.1, -0.2), max(.2, max(R(ind_2008_now,:)) + .1)];

    %ybounds = [0 max(1.2, max(R(ind_2008_now,:)) + .1)];
    ylim(ybounds) %upper limit: 1.2 or the maximal R + 0.1, whichever is greater. Lower: 
    xticks([0 100 200])
    text(150,-0.3,['N=' num2str(sp_dat{inds_to_show(ss),'abd_adults_2015'})],'FontSize',12)

    title([ '\it' sp_dat{inds_to_show(ss),'Genus'}{1} ' ' sp_dat{inds_to_show(ss),'Species'}{1}])
    %text(10,450,'f','FontSize',16,'FontWeight','bold')
    
    if ss == 13
        ylabel('Excess Abundance(r)')
    end
    
    if ss == 30
        xlabel('Distance (m.)')
    end
end

%% Supplamentary figure(s) with H2001 kernel:

load('species_data3.mat','sp_dat')
load('Results/spatial_distributions.mat')
load('Results/Null1_2001_stats.mat')
bin_centers = 5:10:195;

inds_to_show = find((~isnan(sp_dat{:,'HML2001Distance'})) & (sp_dat{:,'avg_log_abd'} >= log(30)));
x_subplots = 5; %numbers of subplots in each dimension
y_subplots = 6;

for ss = 1:length(inds_to_show)

    if rem(ss,30)==1
        figure()
    end

    s1 = subplot(y_subplots, x_subplots, ss - 30*fix(ss/31));
    hold on
    set(s1,'FontSize',14,'FontWeight','bold','LineWidth',1.5);
    x_fill=[bin_centers fliplr(bin_centers)];
    y_fill=[R_sim_envelope(2, :, inds_to_show(ss)) fliplr(R_sim_envelope(1, :, inds_to_show(ss)))];
    fill(x_fill, y_fill , 1,'facecolor',[0.95 0.7 0],'edgecolor','none','facealpha', 0.5);
    plot(bin_centers, R(inds_to_show(ss),:),'-o','MarkerEdgeColor',[0 0.45 0.74], 'MarkerFaceColor', [.3 .75 .93], 'Color', [0 0.45 0.74],'LineWidth',1.5)
    line([0 200],[0 0], 'LineStyle','--', 'Color','black','LineWidth',1.5)

    xlim([0 200])
    %ybounds = [0 max(1.2, max(R(inds_to_show(ss),:)) + .1)];
    ybounds = [min(min(R(inds_to_show(ss),:)) - 0.1, -0.2), max(.2, max(R(inds_to_show(ss),:)) + .1)];
    ylim(ybounds) %upper limit: 1.2 or the maximal R + 0.1, whichever is greater. Lower: 
    xticks([0 100 200])

    title([ '\it' sp_dat{inds_to_show(ss),'Genus'}{1} ' ' sp_dat{inds_to_show(ss),'Species'}{1}])
    text(150,-0.3,['N=' num2str(sp_dat{inds_to_show(ss),'abd_adults_2015'})],'FontSize',12)


    if (ss == 16) || (ss == 46)
        ylabel('Excess Abundance(r)')
    end

    if (ss == 28) || (ss == 58)
        xlabel('Distance (m.)')
    end
end

%%
f1 = gco;
set(f1,'PaperSize',[16 12]); %set the paper size to what you want
set(f1,'Renderer','Painters')
print(f1,'ED Figure 4_2','-dpdf')
```
