## Supplementary material for "Pervasive species-specific repulsion among adult tropical trees": Code appendix: generate_fig_EA_dist1.m

```
 %%%%%%%%%%%%%%%%%%%%%%%%%%%%%%%%%%%%%%%%%%%%%%%%%%%%%%%%%%%%%%%%%%%%
% Generate figures of EA(r) and RND(r)
%%%%%%%%%%%%%%%%%%%%%%%%%%%%%%%%%%%%%%%%%%%%%%%%%%%%%%%%%%%%%%%%%%%%

% Author: Michael Kalyuhny
%
% First Written: 17/07/21
% Last modified: 18/09/22


%% 
cd('./CNDD_HNDD')

%% Set  up parameters to use:

% each row of the matrix is a parameter regime with order:
% [CNDD magnitude , HNDD magnitude , dispresal dist.] (cc,hh,dd)

load('inp_CNDD_HNDD1.mat') 

regs_use = [1 1 2; 3 1 2; 3 3 2];
reg_num = size(regs_use,1);

%% import_null:

ref = cell(1,2);
for dd = 1:length(dist_vals)
    ref{dd} = load(['ref_HNDD_D_' sprintf('%0.4g',dist_vals(dd)) '_analysis_results.mat']);
end

%% import data:

% The analysis_result files for these regimes are provided

dist_axis = ref{1}.dist_bin_centers;

EA = nan(reg_num,length(dist_axis));

for rr = 1:reg_num %run over the different regimes
    dat = load([inp_all(regs_use(rr,1),regs_use(rr,2),regs_use(rr,3)).output_file '_analysis_results.mat']);
    
    EA(rr,:) = nansum( (dat.log_ND - ref{regs_use(rr,3)}.log_ND).*(dat.samps_per_bin') ) / sum(dat.samps_per_bin);
end


%% plot data:
% [CNDD magnitude , HNDD magnitude , dispresal dist.]
%regs_use = [1 1 2; 3 1 2; 3 3 2];

cols = [.5 .85 0 ; 0 .1 .9 ; 0 .1 .9]; %Magnitude of CNDD; Q_C = 0: green; Q_C = 6: blue, QC =12: purple
syms = {'-x', '-x', '-o'}; % %Magnitude of HNDD; Q_H = 0: x; Q_C = 6: o
legend_entries = {'Q_C=0, Q_H=0 (no DD)', 'Q_C=6, Q_H=0', 'Q_C=6, Q_H=6', 'DD range'};
marker_sizes = [10 10 8];

f = figure();

s1 = subplot(1,1,1);
hold on
set(s1,'FontSize',16,'FontWeight','bold','LineWidth',2.5);
for rr = 1:reg_num
    plot(dist_axis,EA(rr,:),syms{rr},'color',cols(rr,:),'MarkerFaceColor',cols(rr,:),'LineWidth',2,'MarkerSize',marker_sizes(rr))
end

plot([0 7],[0.15 0.15], '-', 'color', [.85 .15 .0], 'LineWidth', 6); 
line([0 150],[0 0], 'LineStyle', ':', 'color', 'black', 'LineWidth', 2)
ylabel('Excess neighborhood Abundance')
xlabel('Distance (m.)')
xlim([0 150])
%ylim([-2 1.1])
text(20,0.2,'Aggregated w.r.t. DL','FontSize',14,'FontWeight','bold')
text(20,-0.2,'Overdispersed w.r.t. DL','FontSize',14,'FontWeight','bold')
legend(legend_entries);

%%

set(f,'PaperSize',[10 12]); %set the paper size to what you want
print(f,'Fig_SI_EAr','-dpdf')

cd('..')
```
