## Supplementary material for "Pervasive species-specific repulsion among adult tropical trees": Code appendix: generate_sp_dat_table1.m

```
%%%%%%%%%%%%%%%%%%%%%%%%%%%%%%%%%%%%%%%%%%%%%%%%%%%%%%%%%%%%%%%%%%%
%%% Generation species data table  - first step
%%%%%%%%%%%%%%%%%%%%%%%%%%%%%%%%%%%%%%%%%%%%%%%%%%%%%%%%%%%%%%%%%%%
%
% Author: Michael Kalyuzhny
%
% Date created: 21/08/2020
% Date last modified: 20/01/2022
%
%%%%%%%%%%%%%%%%%%%%%%%%%%%%%%%%%%%%%%%%%%%%%%%%%%%%%%%%%%%%%%%%%%%
% Description:
%
% In this step, info. on species is calculated and obtained from various sources, before computing the main spatial
% statistics of interest.
%
%% Load and prepare the species data table:

%Load species data and preserve only needed:
sp_dat = readtable('Tree_data_reduced.xlsx');
Rep_thresh_use = sp_dat{:,'Rep_Threshold'}*2/3; %use 2/3 of reproductive threshold sensu Foster, like most works
sp_dat = addvars(sp_dat,Rep_thresh_use,'After','Rep_Threshold');

%% Merge full BCI data:

% Input: set of csv files downloaded from Condit et al (2019). They were imported into R and transformed into CSV format with the names 'bci_1985.csv', 'bci_1990.csv',.... They are joined into one cell array. 
% This part can be commented out since it only needs to run once to generate the unified BCI dataset:
% 
bci = cell(1,7);
years_bci = 1985:5:2015;

for yy = 1:7
    bci{yy} = readtable(['../species metadata/bci_' num2str(years_bci(yy)) '.csv']);
end

save('bci_trees_data.mat','bci')

%%
load('bci_trees_data.mat','bci')

% Correct error in importing: broken code for 'anacex' in two censuses:

bci{6}(strcmp(bci{6}{:,'sp'},'a cex'),'sp') = {'anacex'};
bci{7}(strcmp(bci{7}{:,'sp'},'a cex'),'sp') = {'anacex'};

%% Analyze BCI data and add to sp_dat:

%indexes of living trees in every survey
alive = false(size(bci{1},1),7); %the indexes of dead and alive trees. missing (with no coordinates) are listed as dead
for yy=1:7
    alive(:,yy) = strcmp(bci{yy}{:,'status'},'A') & ~isnan(bci{yy}{:,'gx'}) & ~isnan(bci{yy}{:,'gy'});
end

% Create variables:
abd_adults_1985 = nan(size(sp_dat,1), 1); %abundance of adults in 1985
abd_adults_2015 = nan(size(sp_dat,1), 1); %abundance of adults in 2015
comparable_trees = nan(size(sp_dat,1), 1); %individuals with similar or larger size than the reproductive threshold of this species
avg_log_abd = nan(size(sp_dat,1), 1); %average log abundance (over time)
mor_events = zeros(size(sp_dat,1), 1); %number of demographic events
rec_events = zeros(size(sp_dat,1), 1);
point_1985 = cell(1,size(sp_dat,1)); %the point pattern in 1985
point_2015 = cell(1,size(sp_dat,1));
point_all_by_species = false(size(alive,1), size(sp_dat,1)); %data on all the trees that are comparable to this species

% Compute variables for species:

for sp = 1:size(sp_dat,1)
    
    sp_code_now = sp_dat{sp,'SpeciesCode'}{1}; %what is the current species code

    inds_in_bci = (ismember(bci{1}{:,'sp'},sp_dat{sp,'SpeciesCode'})); %index is consistent in all data tables. Vector of binary indexes of individuals of this species in BCI
    
    %%% Compute abundance: %%%%%%%%%%%%%%%%%%%%%%%%
    
    comp_trees_years = zeros(1,7); %how many comparable trees in each year? comparables = with similar reproductive threshols
    abds = zeros(1,7); %what are the abundances, every year?
    
    for yy = 1:7 %all surveys
        comp_trees_years(yy) = sum((bci{yy}{:,'dbh'} >= (10*sp_dat{sp,'Rep_thresh_use'})) & alive(:,yy));
        abds(yy) = sum(inds_in_bci & alive(:,yy) & (bci{yy}{:,'dbh'} >= (10*sp_dat{sp,'Rep_thresh_use'})));
    end
    
    % avoid zeroes (happens once with one species):
    abds(abds == 0) = 0.5; %half an individual
    
    comparable_trees(sp) = round(mean(comp_trees_years));
    avg_log_abd(sp) = mean(log(abds));
    abd_adults_1985(sp) = abds(1);
    abd_adults_2015(sp) = abds(7);
    
    %%% Compute number of recruitment and mortality events (and spatial distributions): %%%%%%%%%%%
        
    % Identify living adults of this species in each survey:
    living_adults = false(size(bci{1},1),7); %the indexes of living_adults
    for yy = 1:7
        living_adults(:,yy) = (inds_in_bci & alive(:,yy) & (bci{yy}{:,'dbh'} >= (10*sp_dat{sp,'Rep_thresh_use'})));
    end
    
    %count events:
    for yy = 1:6
        mor_events(sp) = mor_events(sp) + sum(living_adults(:,yy) & ~living_adults(:,yy+1));
        rec_events(sp) = rec_events(sp) + sum(~living_adults(:,yy) & living_adults(:,yy+1));
    end
    
    point_1985{sp} = [bci{1}{living_adults(:,1),'gx'} bci{1}{living_adults(:,1),'gy'}];
    point_2015{sp} = [bci{7}{living_adults(:,7),'gx'} bci{7}{living_adults(:,7),'gy'}];
    
    % Export all trees with size threshold at or above the threshold of this species:
    
    point_all_by_species(:,sp) = ((bci{7}{:,'dbh'} >= (10*sp_dat{sp,'Rep_thresh_use'})) & alive(:,7));
end

points_all = bci{7}{:,{'gx', 'gy'}};

sp_dat = addvars(sp_dat, comparable_trees, avg_log_abd, abd_adults_1985, abd_adults_2015, mor_events, rec_events);

save('Results/spatial_distributions.mat','point_1985','point_2015')
save('Results/spatial_distribution_all_trees.mat', 'point_all_by_species', 'points_all');

%% save main output:
save('species_data1.mat','sp_dat')
```
