## Supplementary material for "Pervasive species-specific repulsion among adult tropical trees": Code appendix: README.docx

### How to use the code appendix?

Author: Michael Kalyuzhny

Code last updated: 12/01/2023

This code appendix is aimed at reproducing all the analyses, figures and tables. The code is written in the Matlab™. The master script that calls all other scripts is ‘Full_analysis.m’, which calls all other scripts and generates all results. Some graphics saving pieces of the code may be sensitive to the version of Matlab or screen resolution. Also, some parts may depend on the stochastic realization that is used (i.e., what sample of the simulation to visualize?).

**Code input**

The code requires two inputs: one is ‘Tree_data_reduced.xlsx’, a data file on tree characteristics that is provided. The second is a set of .csv files, each containing data from one BCI survey. The R data for that can be downloaded from <https://datadryad.org/stash/dataset/doi:10.15146/5xcp-0d46> and it needs to be separated into separate CSV files as explained in the code.

**Code output**

The code generates many intermediate data files. Because they may take long to generate (many days using a 16 core CPU), some intermediate result files (with extension ‘.mat’) are included. The final output is a set of figures and tables.

**Folder structure**

The provided folder contains three sub folders: ‘Results’, which contains several scripts and intermediate results, ‘Clustered_disp’, which performs the analysis of the clustered dispersal and ‘CNDD_HNDD’, which is used to perform the theoretical analysis of the magnitudes of CNDD and HNDD and contains intermediate results as well.

**Structure of specific scripts**

The master script, ‘Full_analysis.m’, calls many scripts and their children. The bullets below presents the direct children of the master script and their own children, in hierarchy. If a script is called by (“child of”) more than one script, it appears here only under the first parent in the linear order in which the results are generated.

- ‘generate_sp_dat_table1.m’ – produces the basic data for each species required for their analysis
- ‘run_sims_null1_2008.m’ – runs the standard null model with the HML2008 dispersal kernel. This script sets the parameters for the run and calls the function of the null. The output (as well as the output of other simulations) is generated in the ‘Results’ folder.
  - ‘Null1.m’ – The script of the Standard null model
- ‘run_sims_null1_2001.m’ – runs the standard null model with the HML2001 dispersal kernel. This script sets the parameters for the run and calls the function of the null.
  - ‘Null1_exp.m’ – the script of the standard null model with an exponential distance distribution.
- ‘run_sims_null2_lag_2008.m’ - runs the ‘Lagged’ model with the HML2008 dispersal kernel. This script sets the parameters for the run and calls the function of the null.
  - ‘Null2_lag.m’ – the script of the ‘Lagged’ null model
- ‘run_sims_null3_LDD_2008.m’ - runs the Standard null model with the LDD dispersal kernel. This script sets the parameters for the run and calls the function of the null.
  - ‘Null3_LDD.m’ – the script of the standard null model with LDD
- ‘run_sims_null4_initial_2008.m’ – runs the transient null model, initiating with the observed distribution and randomizing the observed number of demographic events. This script sets the parameters for the run and calls the function of the null.
  - ‘Null4_init.m’ - the script of the transient null model with LDD.
- ‘run_sims_null6_2008.m’ – runs the fixed null model, where the dynamics happens only on the observed distribution of trees of all species. This script sets the parameters for the run and calls the function of the null.
  - ‘Null6_Fixed_Trees.m’ - the script of the Fixed null model with LDD.
- ‘Analyze_all_stats3_log.m’ – Performs the analysis of all the statistics of all the nulls. Saves them separately also updates the main data table.
  - ‘Analyze_nulls3_1.m’ – performs the analysis for a single variant of the null model for all species, saves resulting stats.
    - ‘sum_pop.m’ – generates a data structure of species composition by samples (time) from the samples in space and time
    - ‘dist_pvals.m’ – performs a two sided test comparing observed statistic to reference distribution
  - ‘Analyze_nulls_dem3_1.m’ – performs the analysis of the transient null model, where only new trees are analyzed.
- ‘Analysis_CNDD_HNDD_Delta7.m’ – runs all theoretical simulations considering the effects of CNDD and HNDD, assuming Delta (range of CNDD and HNDD) is 7 meters. Also generates Figure 3 of the main text and related supplementary figures.
  - ‘sim3.m’ – runs a simulation model with CNDD and HNDD. There are many unused functionalities in this script.
  - ‘sim2_N.m’ – runs a null simulation, without CNDD or HNDD.
  - ‘Stats_regime3.m’ – analyzes a simulation regime (null/with DD) computing many summary statistics.
  - ‘Compose_pieces.m’ – service script to assemble simulation results if simulations were stopped/crashed.
  - ‘complete_sims.m’ – service script to complete all simulations that finished prematurely
    - ‘sim3_continue.m’ – continues the stopped simulations to completion (hopefully).
- ‘Analysis_CNDD_HNDD_Delta20.m’ – runs supplementary simulations considering the effects of CNDD and HNDD, assuming Delta (range of CNDD and HNDD) is 20 meters. Also generates appropriate supplementary figure.
- ‘generate_fig1_figS1_ver2.m’ – Generates Figure 1 and S1
- ‘Create_tab1_S1_2.m’ – Generates result tables
- ‘Generate_Fig2_and_SI_2.m’ – Generates Figure 2 and SI Figures
- ‘generate_fig_EA_dist1.m’ – generates the figure of the theoretical EA(r)
- ‘analyze_sim_clustered_dispersal2.m’ – master script for all analyses of clustered dispersal
  - ‘Run_sim_clustered_dispersal1.m’ – runs the simulations of clustered dispersal, their respective null models, and also runs their analysis
    - ‘Sim_clustered.m’ – function of the actual simulation with cluster dispersal
    - ‘Stats_regime_clustered_disp.m’ – analyzes the clustered simulation results
    - ‘sim2_NC.m’ – function of the actual null model simulation incorporating two stages of dispersal
