## Supplementary material for "Pervasive species-specific repulsion among adult tropical trees": Code appendix: Null2_lag.m

```
%%%%%%%%%%%%%%%%%%%%%%%%%%%%%%%%%%%%%%%%%%%%%%%
% Null model 2 - recruits are added after a time lag
%%%%%%%%%%%%%%%%%%%%%%%%%%%%%%%%%%%%%%%%%%%%%%%
%
% Author: Michael Kalyuzhny
%
% First written: 21/08/2020
% Last used: 06/01/2021
%
%%%%%%%%%%%%%%%%%%%%%%%%%%%%%%%%%%%%%%%%%%%%%% 
%% Description
%
%  Main community matrix - 3 columns: x,y,species ID Seeds are generated
% (with same attributes) and dispersed with torus BC. immigrants are
%  added from a species pool with equal probability for every species. 
%
% All time is in sweeps. Fixed number of samples is taken, after equilibration. 
% In this version of the null model, recruited individuals "wait" for inp.stack_size time steps before becoming adults.
%%%%%%%%%%%%%%%%%%%%%%%%%%%%%%%%%%%%%%%%%%%%%%%%%%%
%%
function [com_samp, time_samp, runtime] = Null2_lag(inp)

% %% Input patameters
% 
% inp.J = 1600; %number of individuals in the forest
% inp.L = 200; %landscape edge (meters)
% inp.S_reg = 30; %number of species in regional pool
% inp.b = 2; %'b' parameter of the 2DT kernel (sensu Nathan 2012), = p+1 (sensu Clark 1999), representing heavy tailedness
% inp.a = 25; %second parameter of the 2DT
% inp.imm_prob = 0.01; %number of immigrating seeds to add
% inp.stack_size = inp.J;

% % Time and sampling parameters (time in sweeps):
% inp.samps_tot = 100; %how many samples to take?
% inp.samp_freq = 10; %how many sweeps between samples?
% inp.first_samp = 0; %after how many sweeps to take first sample?
% inp.print_freq = 10;
% inp.output_file = 'sim2x_1.mat';

%% Initialization

%all time is in sweeps (J events)
t_now = 0;
next_samp_time = inp.first_samp;
time_samp = NaN(1,inp.samps_tot,'single'); %when were the samples taken?
next_samp_ind = 1; %index of next sample
stop_samp_ind = inp.samps_tot + 1; %when this is the next sample - stop the run
next_print = inp.print_freq;
mkdir(inp.output_file) % create a temporary directory for the samples
time_add = 1/inp.J; %the amount of time (in sweeps) that = a time step

% Generate initial community
com = zeros(inp.J,3,'single'); %this is the community composition, main dynamic variable. first two columns are coordinates. 3rd is species ID, 4th is quadrat (using linear indexing)
com(:,1:2) = rand(inp.J,2,'single')*inp.L; %set uniform random coordinates
com(:,3) = randi(inp.S_reg,inp.J,1,'single'); %draw species from uniform dist.

% Generate initial stack:
stack = zeros(inp.stack_size,3,'single');
stack(:,1:2) = rand(inp.stack_size,2,'single')*inp.L; %set uniform random coordinates
stack(:,3) = randi(inp.S_reg,inp.stack_size,1,'single'); %draw species from uniform dist.
stack_next_replace = 1; %next index to replace

%% Main Loop:
tic

while next_samp_ind < stop_samp_ind

    
    if rand>inp.imm_prob % No immigration: generate and disperse seed:
        seed = com(randi(inp.J,'single'),:);
        u = rand; %temporary variable, to be transformed into distance:
        r = sqrt(((1-u)./(inp.a.^(2*inp.b-2))).^(1./(1-inp.b)) - inp.a^2 ); %transform to 2DT distance distribution
        theta = rand*2*pi; %direction
        seed(1) = seed(1) + r*cos(theta); %perform seed displacement - x coord.
        seed(2) = seed(2) + r*sin(theta); %perform seed displacement - y coord.

        % Enforce torus boundaries:
        seed(1:2) = rem(seed(1:2),inp.L); %for coordinates > L - performs wrapping, also shortens negative values to be > -L
        seed(seed<0) = inp.L + seed(seed<0); %for negative values
    else % Immigration:
        seed = [rand(1,2,'single')*inp.L randi(inp.S_reg,'single')]; %generate immigrating seeds
    end
    
    %replace a random tree with the next tree in the stack and assign the new seed in it's stead
    
    com(randi(inp.J,'single'),:) = stack(stack_next_replace,:); % replace a random tree with a seed from the stack
    stack(stack_next_replace,:) = seed; %replace the seed in the stack with the produced seed
    stack_next_replace = rem(stack_next_replace,inp.stack_size) + 1; %next time use next stack index
        
    %print time:
    if t_now >= next_print
        disp(['Runtime: ' sprintf('%0.8g',toc) ' sec., Time: ' sprintf('%0.8g',t_now) ' gen.'])
        next_print = next_print + inp.print_freq;
    end
  
    if t_now >= next_samp_time %if it's time to sample
        time_samp(next_samp_ind) = t_now; %record time
        
        save([inp.output_file '/s_' num2str(next_samp_ind) '_' inp.output_file '.mat'],'com');
        
        %next sampling:
        next_samp_time = next_samp_time + inp.samp_freq;
        next_samp_ind = next_samp_ind + 1;
    end
    
    t_now = t_now + time_add; %progress time
end

runtime = toc;

% Compose the pieces and save:

com_samp = NaN(inp.J,3,inp.samps_tot,'single');

for ss = 1:inp.samps_tot
    l = load([inp.output_file '\s_' num2str(ss) '_' inp.output_file '.mat'],'com');
    com_samp(:,:,ss) = l.com;
end

save([inp.output_file '.mat'], '-v7.3')
rmdir(inp.output_file, 's') % delete temporary directory for the samples


disp(['Finished! Runtime: ' sprintf('%0.8g',toc) ' sec., Time: ' sprintf('%0.8g',t_now) ' gen.'])
end
```
