## Supplementary material for "Pervasive species-specific repulsion among adult tropical trees": Code appendix: Null4_init.m

```
%%%%%%%%%%%%%%%%%%%%%%%%%%%%%%%%%%%%%%%%%%%%%%%
% Null model 4 - fix initial state and number of events
%%%%%%%%%%%%%%%%%%%%%%%%%%%%%%%%%%%%%%%%%%%%%%%
%
% Author: Michael Kalyuzhny
% First written: 21/10/2020
% Last modified: 06/01/2020
%
% In this version the initial point pattern of the species is preserved, as well as the number of mortality and
% recruitment events. the order of the events and the identity of individuals are randomized and the dispersal distances
% are drawn from the kernel. Iterations with extinctions are ignored (recalculated).
%
% input: initial point pattern, sizes of the landscape (Lx, Ly), the 'a' paramter of the 2DT kernel, the number of
% recruitment and mortality events and the number of resamples to generate. Filename is location to save the results. If
% is 'nan' then no file is saved
% output: a cell array final point patterns, for each realization - the indexes of new individuals

%%
function [points_final, is_new] = Null4_init(points_initial, Lx, Ly, a, mor_events, rec_events, resamps, filename)

% %example input:
% points_initial = [200:10:300;200:10:300]';
% Lx = 1000;
% Ly = 500;
% a=25;
% mor_events = 3;
% rec_events = 7;
% resamps = 100;

%precalculations:
tot_events = mor_events + rec_events;
points_final = cell(1,resamps); %save final spatial patterns
is_new = cell(1,resamps); %save for each point in each pattern - is it a new recruit?
ii = 1;

while ii <= resamps
    points = [points_initial ; nan(tot_events,2)]; % dynamic point pattern - preallocation
    pop_now = size(points_initial,1); %initial abundance
    
    next_event = [ones(1,rec_events) zeros(1,mor_events)];
    next_event = next_event(randperm(tot_events)); %reshuffle events! This is their order

    for tt = 1:tot_events

        if next_event(tt)==1 %recruitment
            seed = points(randi(pop_now,'single'),:);
            u = rand; %temporary variable, to be transformed into distance:
            r = sqrt(((1-u)./(a.^(2))).^(-1) - a^2 ); %transform to 2DT distance distribution
            theta = rand*2*pi; %direction
            seed(1) = seed(1) + r*cos(theta); %perform seed displacement - x coord.
            seed(2) = seed(2) + r*sin(theta); %perform seed displacement - y coord.

            % Enforce torus boundaries:
            seed(1) = rem(seed(1),Lx); %for coordinates > L - performs wrapping, also shortens negative values to be > -L
            seed(2) = rem(seed(2),Ly); %for coordinates > L - performs wrapping, also shortens negative values to be > -L
            if seed(1) < 0 
                seed(1) = Lx + seed(1); %for negative values
            end
            if seed(2) < 0 
                seed(2) = Ly + seed(2); %for negative values
            end

            %establish:
            pop_now = pop_now + 1;
            points(pop_now,:) = seed; %save the seed at the last index

        else %mortality
            points(randi(pop_now,'single'),:) = points(pop_now,:); %transform a random individual (who dies) into the last invididual
            points(pop_now,:) = nan(1,2); %eliminate last individual
            pop_now = pop_now-1;
            
            if pop_now == 0 %if species is extinct - stop this loop
                break
            end
        end
    end

    
    if (tt==tot_events) && (pop_now>0) %if itteration was completed without an extinction
        %save results:
        points_final{ii} = points(1:pop_now,:); 
        is_new{ii} = ~ismember(points_final{ii}, points_initial,'rows');
        
        ii=ii+1; %promote ii only if the simulation finished!
    end

end

if ~isnan(filename)
    save(filename,'points_final','is_new')
end

end
```
