## Supplementary material for "Pervasive species-specific repulsion among adult tropical trees": Code appendix: Null6_Fixed_Trees.m

```
%%%%%%%%%%%%%%%%%%%%%%%%%%%%%%%%%%%%%%%%%%%%%%%
% Null model 6 - using the observed distribution of trees
%%%%%%%%%%%%%%%%%%%%%%%%%%%%%%%%%%%%%%%%%%%%%%%
%
% Author: Michael Kalyuzhny
%
% First written: 21/08/2020
% Last used: 02/03/2021
%
%%%%%%%%%%%%%%%%%%%%%%%%%%%%%%%%%%%%%%%%%%%%%% 
%% Description
%
% This is the standard form of the Null model, a spatialy explicit version of Neutral Theory on a Torus
%
%  Main community matrix - 4 columns: x,y,species ID, quadrat destinations are generated
% (with same attributes) and dispersed with torus BC. immigrants are
%  added from a species pool with equal probability for every species. 
%
% All time is in sweeps. Fixed number of samples is taken, after equilibration. 
% Dispersal is performed "backwards" and the nearest tree is found to be the parent of an extant tree.
% To save computation of nearest neighbor distances, we perform computation only within quadrats.
%%%%%%%%%%%%%%%%%%%%%%%%%%%%%%%%%%%%%%%%%%%%%%%%%%%
%%
function [com_samp, time_samp, runtime] = Null6_Fixed_Trees(inp)

% %% Input patameters
% 
% inp.J = 1600; %number of individuals in the forest
% inp.L = 200; %landscape edge (meters)
% inp.S_reg = 30; %number of species in regional pool
% inp.b = 2; %'b' parameter of the 2DT kernel (sensu Nathan 2012), = p+1 (sensu Clark 1999), representing heavy tailedness
% inp.a = 25; %second parameter of the 2DT
% inp.imm_prob = 0.01; %number of immigrating destinations to add
% 
% % Time and sampling parameters (time in sweeps):
% inp.samps_tot = 100; %how many samples to take?
% inp.samp_freq = 10; %how many sweeps between samples?
% inp.first_samp = 0; %after how many sweeps to take first sample?
% inp.print_freq = 10;
% inp.output_file = 'sim2x_1.mat';

%% Initialization

%all time is in sweeps (J events)
t_now = 0;
next_samp_time = inp.first_samp;
time_samp = NaN(1,inp.samps_tot,'single'); %when were the samples taken?
next_samp_ind = 1; %index of next sample
stop_samp_ind = inp.samps_tot + 1; %when this is the next sample - stop the run
next_print = inp.print_freq;
mkdir(inp.output_file) % create a temporary directory for the samples
time_add = 1/inp.J; %the amount of time (in sweeps) that = a time step
LD = inp.Ly/inp.D; %how many quadrats along the y edge?

% Generate initial community
com = zeros(inp.J,4,'single'); %this is the community composition, main dynamic variable. first two columns are coordinates. 3rd is species ID, 4th is quadrat (using linear indexing)
com(:,1:2) = inp.trees; %set uniform random coordinates
com(:,3) = randi(inp.S_reg,inp.J,1,'single'); %draw species from uniform dist.
com(:,4) = ceil(max((inp.Ly - com(:,2)),1)/inp.D) + LD*floor(max(com(:,1)-0.001,0)/inp.D); %compute quadrat for every tree. the min/max and 0.00001 are ment to deal with extreme cases of exactly 0 or L.

% Compute the indexes of trees in each quadrat:
quad_trees = cell(1,LD*(inp.Lx/inp.D));
for ii = 1:length(quad_trees)
    quad_trees{ii} = [find(com(:,4) == ii) com(find(com(:,4) == ii),1:2)]; %first column: index in com, 2nd&3rd column: x,y coordinates
end

%% Main Loop:
tic

while next_samp_ind < stop_samp_ind

    to_rep = randi(inp.J,'single'); %Tree to replace
    if rand>inp.imm_prob % No immigration: find parent tree:
        destination = com(to_rep,1:2); %begin with parent tree
        u = rand; %temporary variable, to be transformed into distance:
        r = sqrt(((1-u)./(inp.a.^(2*inp.b-2))).^(1./(1-inp.b)) - inp.a^2 ); %transform to 2DT distance distribution
        theta = rand*2*pi; %direction
        destination(1) = destination(1) + r*cos(theta); %perform destination displacement - x coord.
        destination(2) = destination(2) + r*sin(theta); %perform destination displacement - y coord.

        % Enforce torus boundaries:
        destination(1) = rem(destination(1),inp.Lx); %for coordinates > L - performs wrapping, also shortens negative values to be > -L
        destination(2) = rem(destination(2),inp.Ly); %for coordinates > L - performs wrapping, also shortens negative values to be > -L
        if destination(1) < 0 
            destination(1) = inp.Lx + destination(1); %for negative values
        end
        if destination(2) < 0 
            destination(2) = inp.Ly + destination(2); %for negative values
        end
        
        %find the quadrat of the destination:
        destination_quadrat = ceil(max((inp.Ly - destination(:,2)),1)/inp.D) + LD*floor(max(destination(:,1)-0.001,0)/inp.D); %compute linear index of quadrat 
        
        % Find nearest tree:        
        xdists = abs(quad_trees{destination_quadrat}(:,2)-destination(1));
        ydists = abs(quad_trees{destination_quadrat}(:,3)-destination(2));
        dd = sqrt(xdists.^2+ydists.^2); %distances
        
        [~, ind_min_dist] = min(dd); %this is the index WITHIN THE QUDRAT of the closest tree!
        
        com(to_rep,3) = com(quad_trees{destination_quadrat}(ind_min_dist, 1), 3); %perform replacement

    else % Immigration - replace by random sample from the pool:
        com(to_rep,3) = randi(inp.S_reg,1,'single');
    end
        
    %print time:
    if t_now >= next_print
        disp(['Runtime: ' sprintf('%0.8g',toc) ' sec., Time: ' sprintf('%0.8g',t_now) ' gen.'])
        next_print = next_print + inp.print_freq;
    end
  
    if t_now >= next_samp_time %if it's time to sample
        time_samp(next_samp_ind) = t_now; %record time
        
        save([inp.output_file '/s_' num2str(next_samp_ind) '_' inp.output_file '.mat'],'com');
        
        %next sampling:
        next_samp_time = next_samp_time + inp.samp_freq;
        next_samp_ind = next_samp_ind + 1;
    end
    
    t_now = t_now + time_add; %progress time
end

runtime = toc;

% Compose the pieces and save:

com_samp = NaN(inp.J,3,inp.samps_tot,'single');

for ss = 1:inp.samps_tot
    l = load([inp.output_file '\s_' num2str(ss) '_' inp.output_file '.mat'],'com');
    com_samp(:,:,ss) = l.com(:,1:3);
end

save([inp.output_file '.mat'], '-v7.3')
rmdir(inp.output_file, 's') % delete temporary directory for the samples
pause(3); %wait until delete is done, just in case

disp(['Finished! Runtime: ' sprintf('%0.8g',toc) ' sec., Time: ' sprintf('%0.8g',t_now) ' gen.'])
end
```
