## Supplementary material for "Pervasive species-specific repulsion among adult tropical trees": Code appendix: run_sims_null1_2001.m

```
%%%%%%%%%%%%%%%%%%%%%%%%%%%%%%%%%%%%%%%%%%%%%%%%%%%%%%%%%%%%%%%%%%%
%%% Run simulation of null model 1 for every species with dispersal distance in Muller-Landau 2001
%%%%%%%%%%%%%%%%%%%%%%%%%%%%%%%%%%%%%%%%%%%%%%%%%%%%%%%%%%%%%%%%%%%
%
% Author: Michael Kalyuzhny
%
% Date created: 23/08/2020
% Date last modified: 04/01/2021
%
% Load the species table and run the simulations for the parameters of each species. The simulations are run on a
% lanscape of 1200X1200 meters but with the same density as in BCI, which ia assounted for in the calculation of J.
% Immigration is small, only to maintain some diversity.
%% set up:

load('species_data1.mat','sp_dat')

cd('Results') %generate all results in this folder

species_run = find(~isnan(sp_dat{:,'HML2001Distance'})); %these are the species to run the simulation for

disp('Running Null1 with H2001 distances: ')
tic
parfor spp = 1:length(species_run)
    sp = species_run(spp)
    inp = struct;
    inp.J = round(1200*1200/(1000*500) * sp_dat{sp,'comparable_trees'}); %number of individuals in the forest
    inp.L = 1200; %landscape edge (meters)
    inp.S_reg = 300; %number of species in regional pool
    inp.a = sp_dat{sp,'HML2001Distance'}; %dispersal distance
    inp.imm_prob = 10/inp.J; %number of immigrating seeds to add
    
    % Time and sampling parameters (time in sweeps):
    inp.samps_tot = 1000; %how many samples to take?
    inp.samp_freq = 10; %how many sweeps between samples?
    inp.first_samp = 5000; %after how many sweeps to take first sample?
    inp.print_freq = 1000;
    inp.output_file = ['null1_2001_' sp_dat{sp,'SpeciesCode'}{1}];
    
    [~, ~, ~] = Null1_exp(inp)
end
toc

cd('..') %Go back to parent folder
```
