## Supplementary material for "Pervasive species-specific repulsion among adult tropical trees": Code appendix: run_sims_null4_initial_2008.m

```
%%%%%%%%%%%%%%%%%%%%%%%%%%%%%%%%%%%%%%%%%%%%%%%%%%%%%%%%%%%%%%%%%%%
%%% Run simulation of null model 4 (conditioned on initial state) for every species
%%%%%%%%%%%%%%%%%%%%%%%%%%%%%%%%%%%%%%%%%%%%%%%%%%%%%%%%%%%%%%%%%%%
%
% Author: Michael Kalyuzhny
%
% Date created: 23/08/2020
% Date last modified: 06/01/2021
%
% In this version of the null model, the initial state (coordinates) of the species, and the number of mortality and recruitment
% events are preserved
%% set up:

load('species_data1.mat','sp_dat')
load('Results/spatial_distributions.mat','point_1985','point_2015')

cd('Results') %generate all results in this folder

species_run = find(~isnan(sp_dat{:,'HML2008AlphaFitted'})); %these are the species to run the simulation for
resamps = 20000; %for every species

disp('Running Null4 (Initial state) with H2008 distances: ')
tic
parfor spp = 1:length(species_run)
    sp = species_run(spp)

    a = sqrt(exp(sp_dat{sp,'HML2008AlphaFitted'}));
    rec_events = sp_dat{sp,'rec_events'};
    mor_events = sp_dat{sp,'mor_events'};
    filename = ['null4_2008_' sp_dat{sp,'SpeciesCode'}{1} '.mat'];
    
    
    [~, ~] = Null4_init(point_1985{sp}, 1000, 500, a, mor_events, rec_events, resamps, filename)

end
toc

cd('..') %Go back to parent folder
```
